# Evolution of the insect PPK gene family

**DOI:** 10.1101/2021.03.30.437681

**Authors:** Jose Manuel Latorre-Estivalis, Francisca Cunha Almeida, Gina Pontes, Hernán Dopazo, Romina Barrozo, Marcelo Gustavo Lorenzo

## Abstract

Insect *Pickpocket* (PPK) receptors mediate the detection of stimuli of diverse sensory modalities, therefore having a relevant role for environmental sounding. Notwithstanding their relevance, studies on their evolution are scarce. We have analyzed the genomes of 26 species belonging to 8 insect orders (Blattodea, Orthoptera, Hemiptera, Phthiraptera, Hymenoptera, Lepidoptera, Coleoptera, and Diptera) to identify their PPK repertoires and study the evolution of this gene family. PPKs were detected in all genomes analyzed, with a total of 578 genes identified that distributed in 7 subfamilies. Our phylogenetic analysis allowed clarifying that the *ppk17* gene appears to be the most divergent family member, composing a new group designed as subfamily VII. According to our analysis, PPKs evolved under a birth-and-death model that generated lineage-specific expansions usually located in clusters and the effect of strong purifying selection was seen for several orthogroups. Subfamily V was the largest one, presenting half of all PPKs studied, including a mosquito-specific expansion that can be considered a new target for pest control. Consistently with their sensory role, PPKs present a high gene turnover that generated considerable variation in the size of insect repertoires: *Musca domestica* (59), *Blattella germanica* (41), *Culex quinquefasciatus* (48), and *Aedes albopictus* (51) presented the largest PPK repertoires, while *Pediculus humanus* (only *ppk17*), bees and ants (6-9) had the smallest ones. The expansions identified in *M. domestica* and *Bl. germanica* also show promise as specific targets for controlling these nuisance insects. Our phylogenetic analysis revealed a subset of prevalent PPKs across insect genomes, suggesting a very conserved function that resembles the case of *antennal* ionotropic receptors. Finally, we identified new highly conserved residues in the second transmembrane domain that may be key for receptor function. Besides, more than a hundred PPK sequences presented calmodulin binding motifs, suggesting that at least some members of this family may amplify sensory responses as previously proposed for *D. melanogaster ppk25*. Overall, our study is a first attempt to characterize the evolutionary history of this family of sensory receptors, revealing relevant unknown features and clade-specific expansions.

## INTRODUCTION

The insect pickpocket (PPK) family belongs to the much larger Degenerin/Epithelial Sodium Channel (Deg/ENaC) gene superfamily, first described when the genetic bases of mechanosensory pathways were studied in the nematode *Caenorhabditis elegans* (García-Añoveros et al. 1995). This gene superfamily includes seven families, three of which were first described in vertebrates: ENaC, Acid-Sensing Ion Channels (ASICs), and Brain-Liver-Intestine Sodium Channel (BLINaC)/Human Intestine Sodium Channel (hINaC). Other Deg/ENaC families were reported in invertebrates: the Degenerins from *C. elegans*, the *Drosophila* PPK channels, the FMRFamide-gated Sodium Channel (FaNaC) from mollusks, and the FLR-1 receptor that was only identified in *C. elegans* (Kellenberger and Schild 2002). Deg/ENac members encode a diverse array of epithelial Na^+^ channel proteins related to fundamental functions such as Na^+^ absorption, neuron membrane potential control, detection of pH variation, and touch (Kellenberger and Schild 2002). Deg/ENaCs form channels through the union of hetero- or homotrimeric subunits (Canessa et al. 1994), whose identities have an important effect on the pharmacological and kinetic properties of the channel (Benson et al. 2002; Xie et al. 2003). A property of many Deg/ENaCs is their sensitivity to amiloride, an antagonist drug that blocks channel activity in a transient way (Schild et al. 1997).

The sensory abilities of insects have been shaped along their evolution to adapt to a diverse array of ecological conditions. Insects explore their environment through diverse sensory modalities, and chemoreception is probably the best studied to date. Odorant, gustatory, and ionotropic receptors (ORs, GRs, and IRs, respectively) constitute the three main gene families related to insect chemoreception and have received most of the attention in the last two decades (Carey and Carlson 2011; Hansson and Stensmyr 2011; Breer et al. 2019). Insects present a fourth gene family that encodes receptor proteins that can relate to chemosensory processes, i.e. that of PPKs, which has received less attention. The PPKs have a fundamental role in mediating the perception of stimuli of diverse modalities, including water, salts, osmotic potential, pheromones, and mechanosensory properties of their environment (Liu et al. 2003; Zhong et al. 2010; Thistle et al. 2012; Matthews et al. 2019; Pontes et al. 2017; Masagué et al. 2020)

The sequences of PPK proteins are characterized by one highly conserved cysteine-enriched domain in their extracellular loop and two transmembrane domains (Liu et al. 2003; Zelle et al. 2013). According to Zelle et al., (2013), the PPKs of *Drosophila* spp can be phylogenetically divided into six subfamilies, but our knowledge about the PPKs of other insect species is scarce. The lack of comparative genomics and phylogenetic studies on the PPK gene family hinders our understanding of the extent of their conservation and diversification across different insect orders. This gap of knowledge prevents predicting whether the functional roles described for model organisms like *Drosophila* can be extrapolated to other insects. As PPKs are considered exclusive of insects, a proper phylogenetic and functional characterization can have practical implications for developing novel tools to interfere with key functions of pests causing sanitary or economic impact. Here, we study the PPK sequences found in 26 insect genomes belonging to 8 orders to characterize the evolution of the PPK family across insects.

## MATERIAL AND METHODS

### Candidate sequences

Amino acid sequences of PPKs from *D. melanogaster* (Zelle et al. 2013), *Aedes aegypti*, and *Anopheles gambiae* (Matthews et al. 2019) were used as queries in BLASTp searches (with e-value threshold = 0.0001 and the low complexity regions filter deactivated) in different databases (details in Supplementary Table S1) to identify their putative orthologs in: Blattodea (*Blattella germanica*), Orthoptera (*Locusta migratoria*), Hemiptera (*Acyrthosiphon pisum*, *Myzus persicae,* and *Cimex lectularius*), Phthiraptera (*Pediculus humanus)*, Hymenoptera (*Atta cephalotes, Apis mellifera, Bombus impatiens*, and *Camponotus floridanus*), Lepidoptera (*Bombyx mori, Danaus plexippus, Plutella xylostella*, and *Spodoptera frugiperda*), Coleoptera (*Anoplophora glabripennis*, *Dendroctonus ponderosae, Leptinotarsa decemlineata*, and *Tribolium castaneum*), and Diptera (*Aedes albopictus*, *Culex quinquefasciatus, Glossina morsitans*, and *Musca domestica*). Iterative searches were conducted with each new PPK protein sequence as a query until no new genes could be identified for each PPK subfamily or lineage. Translated sequences of PPK genes previously described for *R. prolixus* (Latorre-Estivalis et al. 2017) were also included in the analysis.

### Analysis of PPK sequences

Protein sequences obtained in the similarity searches were characterized at functional and structural levels. The Pfam v.27.0 (Finn et al. 2016) database was used to check for the presence of the amiloride-sensitive sodium channel domain (PFAM00858) characteristic of PPKs using the Batch Web CD-Search tool from the Conserved Domain Database at NCBI (Lu et al. 2020). The presence and number of transmembrane domains predicted were established using TOPCONS (Bernsel, Viklund et al. 2009). The Calmodulin Target Database (http://calcium.uhnres.utoronto.ca/ctdb/ctdb/home.html) was used to identify Calmodulin Binding Motifs (CBMs) in the N- and C- terminal intracellular regions of PPK sequences. In order to provide the best candidates, this analysis was exclusively performed on PPKs with 2 transmembrane domains, CBMs longer than 7 amino acids and normalized scores ≥ 8. Finally, protein sequences were aligned using the G-INS-I strategy in MAFFT v.7 (mafft.cbrc.jp/alignment/server) and different functional and conserved motifs, previously characterized for *D. melanogaster* (Liu et al. 2003) were identified in the alignment.

### Phylogenetic analysis

Sequences were aligned with MAFFT using the G-INS-i strategy, and the following settings: *unaligned level* =0.1; *offset value* = 0.12; *maxiterate* = 1000 and the option “*leave gappy regions*”. The alignment was trimmed using trimAl v1.2 (Capella-Gutierrez et al. 2009) with default parameters except for the gap threshold that was fixed at 0.3. Following trimming, two phylogenetic trees were built. The first one was based on the maximum-likelihood approach by using IQ-tree v 1.6.12 (Nguyen et al. 2015). The branch support was estimated using both the approximate Likelihood Ratio Test based on the Shimodaira-Hasegawa (aLRT-SH) and the ultrafast bootstrap or UFBoot (Hordijk and Gascuel 2005; Minh et al. 2013) procedures. The best-fit amino acid substitution model, reached by IQ-tree, was WAG+F+R9. This model was chosen according to the Bayesian Information Criterion. The second maximum-likelihood phylogenetic tree was obtained using RAxML v8 (Stamatakis 2014) with the PROTCATWAG amino-acid substitution model for tree search and 200 replicates in the bootstrap analysis. Both phylogenetic trees were displayed and edited with FigTree (http://tree.bio.ed.ac.uk/software/figtree). The PPK candidates were annotated based on their relation with those of *D. melanogaster*. The nomenclature adopted in our study implies that candidates without a clear relationship with *D. melanogaster* PPK genes retained the original codes from their corresponding databases.

### Selection analysis

Codeml package from PALM4 (Yang 2007) was used to estimate the non-synonymous to synonymous substitution rate (dN/dS) ratios (*ω*) across several PPK lineages following the methodology implemented in Almeida et al., (2014). Lineages bearing genes found in *D. melanogaster* and having more than 4 orthologs for each of them were included in the analysis. Exceptionally the *ppk2* and *ppk19* lineages of *M. domestica* and the expansion identified for *B. germanica* in Subf-III were also included due to their corresponding expansions even though the previous criteria were not met for these lineages. We fitted different codon-based substitution models (M) using a maximum-likelihood approach. Initially, M0 was used to obtain branch lengths and mean ω. Then, we estimated model parameters and the log likelihood (*L*) of M7 and M8 and compared the *L* of these two models with the likelihood ratio test (LRT; α = 0.05). To reduce the proportion of false positives of the M7 *vs.* M8 comparison, we also used the M8a, which is considered as an alternative null hypothesis of the M8 where the highest *ω* of the distribution across sites is fixed to *ω* = 1. We assumed 2 and 1 degrees of freedom for the M8 *vs.* M7, and the M8 *vs.* M8a comparisons, respectively. Finally, the Bayes Empirical Bayes analysis under M8 (Yang 2005) was used to identify codons under positive selection. Sites were considered under positive selection when the posterior probability (PP) of belonging to the class with ω > 1 was PP > 0.5.

### Family size evolution

The evolution of PPKs was further analyzed by estimating gene birth and death rates. Because this analysis relies on well-curated gene sets, we opted to include the best-annotated insect genomes only. Therefore, we obtained phylogenetic trees for each subfamily (using RAxML with 20 independent runs) using a matrix including the sequences of *D. melanogaster*, *M. domestica*, *G. morsitans*, *Ae. aegypti*, *Ae. albopictus*, *Cx. quinquefasciatus*, *An. gambiae*, *Bo. mori*, *De. plexippus*, *T. castaneum*, *Ap. mellifera*, *Ca. floridanus*, *P. humanus*, *Ac. pisum*, and *R. prolixus*. Besides using these phylogenetic trees for family size analyses, they were also used to resolve the evolution of PPK subfamilies (complementing the information offered by both trees built for the Phylogenetic Analysis section).

We estimated the number of gene duplications and losses by applying the gene tree *vs.* species tree reconciliation method for each orthogroup as in Almeida et al., (2014). To obtain gene birth and death rates, we applied equations 1 (“total time” approach) and 2 (“branch average” approach) from (Almeida et al. 2014), using previously published node divergence times (Vieira et al. 2007; Misof et al. 2014; Logue et al. 2013). Total rates comprise an average across branches, and take into account the number of duplications/losses *per* number of genes in the ancestral node, *per* branch length in time.

## RESULTS

### Analysis of PPK sequences

A total of 578 PPKs were identified in the genomes of the insect species analyzed (Supplementary Table S2). All PPKs presented the PFAM domain 00858 characteristic of the Degenerin/Epithelial Sodium Channel superfamily. The average length of PPKs was 455 amino acids (aa) and Subf- I to VII presented 464, 402, 523, 548, 627, 522, and 436 aa, respectively. Structural analysis revealed the presence of transmembrane domains in most cases, although their numbers were variable (149 PPKs (26%) showed 1 transmembrane domains, 323 PPKs sequences (56%) had two, 46 PPKs (8%) presented between 3 to 5, while for 60 PPKs (10%) no transmembrane domains were identified). A total of 274 sequences (47.4 %) presented between 13 and 14 cysteine residues, 170 (29.4%) between 10 and 12, and 134 (23.2%) 9 or less (Supplementary Table S2), a characteristic feature of the extracellular loop of PPKs.

Based on Liu et al. (2003) findings of *D. melanogaster* PPK sequences, highly conserved amino acid residues were looked for in the PPK sequences found for the species here analyzed (Supplementary Table S2). Firstly, 43.4% of the sequences presented the “T/S-X-h-H-G” motif (where “h” indicates a hydrophobic residue) anteceding the first transmembrane domain (132 exhibiting an initial threonine and 119 a serine). Secondly, the first transmembrane domain presented a conserved tryptophan residue in 381 (66%) PPK sequences (Figure 1). The second transmembrane domain of 395 (68.3%) PPK sequences presented the “GxS” motif (Figure 1). Interestingly, this region of the receptor was consistently conserved in other positions such as the glycine at position 3 (58%), the leucine at position 5 (66%), the serine at position 13 (59%), the glutamic acid at position 16 (69%), and the tyrosine at position 19 (59%). Additionally, before the first cysteine, a set of 256 sequences (44.2%) presented the “F-P-h-h-T-h-C” motif and 335 sequences (60%) had the “G-X-C-X-X-F-N” motif associated with the fourth cysteine. Alanine was the most common residue (i.e. found in 180 candidates representing 31.1% of the sequences) located in the Degenerin (Deg) site. This residue was followed by valine (137), serine (79), glycine (23), isoleucine (7), threonine (6), cysteine (4), asparagine (3), methionine (3), proline (3), arginine (2), glutamic acid (2) and lysine (2). Leucine and histidine were identified in one sequence each. All PPK protein sequences are listed in Supplementary File S1.

**Figure 1.**
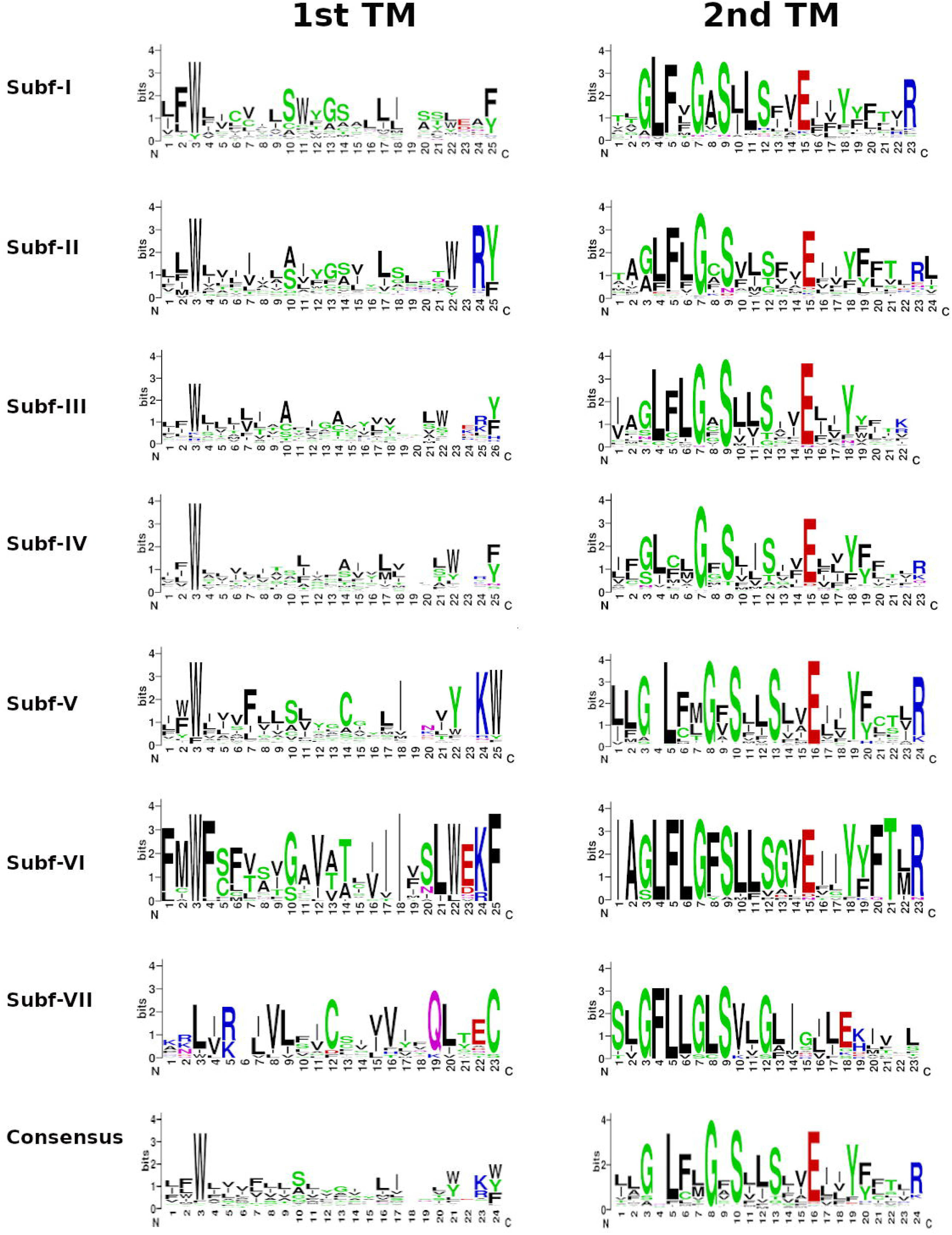
Graphical representation of amino acid sequence alignments of the first (left) and second (right) transmembrane domains of the PPK subfamilies and its consensus. These images were generated using WebLogo Version 2.8.2.

Finally, the analysis of the Calmodulin Binding Motifs (CBMs) revealed that 116 candidates out of 323 sequences with two transmembrane domains had this motif in their sequences. The CBM was located in the N-terminal in 63.7% of the sequences (74 sequences) and in 36.3 % of the sequences (42 sequences) were at the C-terminal regions (Supplementary Table S3). CBMs were identified in PPKs from the seven subfamilies. Some PPK orthogroups, including *ppk15*, *ppk9*, *ppk6*, and in particular Subf-V expansions, such as those of *Cx. quinquefasciatus* (16 sequences) and *T. castaneum* (8 sequences), had a higher prevalence of CBMS in their sequences. The alignment of CBM sequences a certain degree of conservation in PPKs belonging to the same subfamily (Supplementary Figure S1) and presence of positively charged amino acids (red residues in Supplementary Figure S1) interspersed among hydrophobic residues (blue residues on Supplementary Figure S1), a typical features of CBMs (Rhoads and Friedberg 1997).

### PPK evolution in insects

Phylogenetic analyses are summarized in Figure 2, which presents a reduced version of the tree depicted in Supplementary Figure S2. A second extended phylogenetic tree is presented in Supplementary Figures S3. These trees were almost identical, and only a few PPKs belonging to Subf-I and II showed slight changes within the same clades when both trees were compared. In addition, phylogenetic trees (one *per* subfamily) obtained using well-curated genomes (more details in the “Family size evolution” section in Material and Methods) allowed reinforcing the evolutionary relations observed within Sub-III, Subf-V, Sub-VI and Sub-VII and resolving ambiguities in the remaining PPK subfamilies (Supplementary Figures S4-S10).

**Figure 2.**
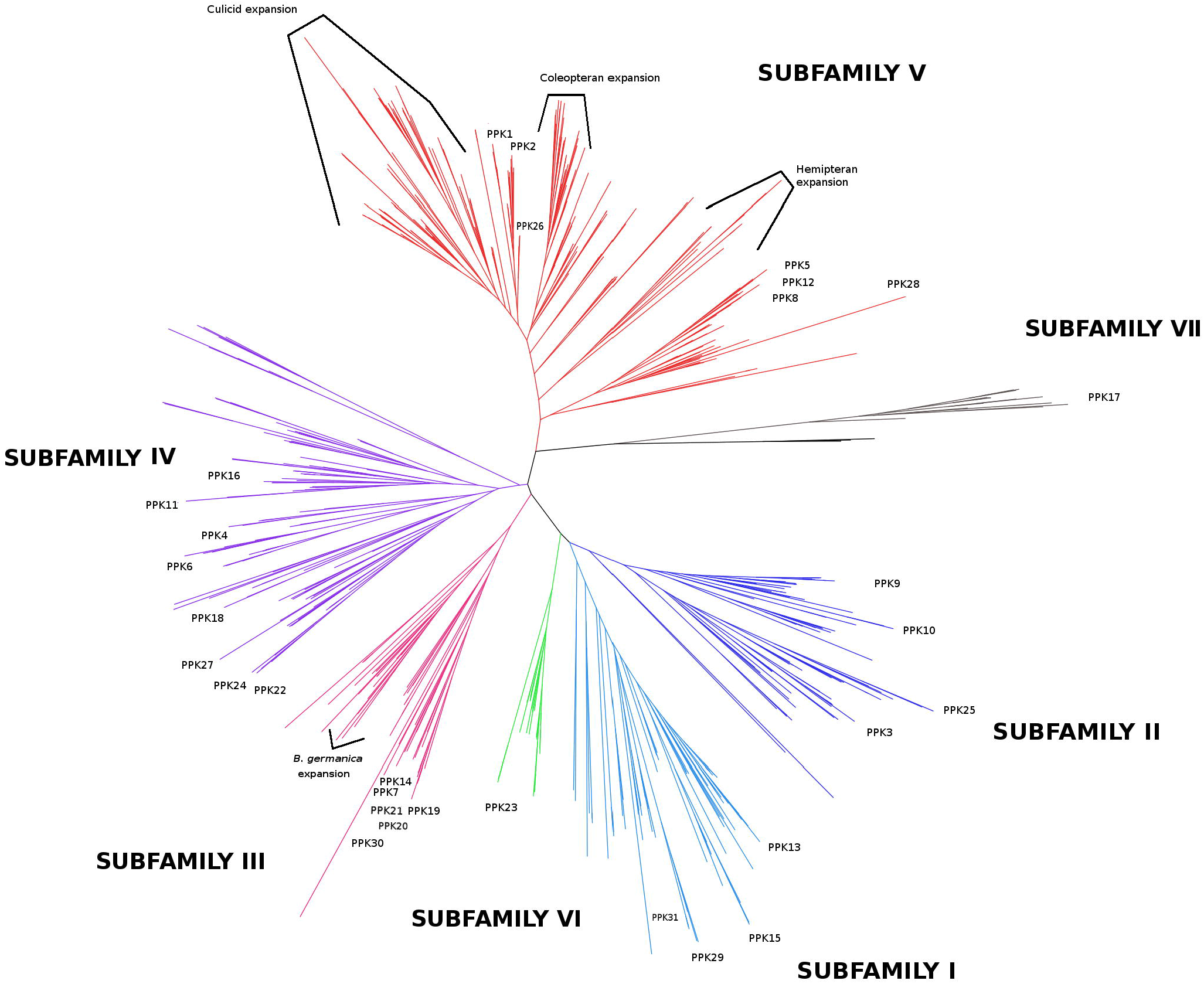
Phylogeny of the PPK receptor gene family. The phylogenetic tree based on a MAFFT alignment was obtained with IQ-Tree using the WAG+F+R9 best-fit model according to Bayesian Information Criterion. The tree was unrooted and three amiloride-sensitive sodium channels (SCNN) from *Rattus norvegicus* were used as an outgroup. PPK subfamilies are displayed in different colors. Branch tags and support values were eliminated to facilitate visualization and extended trees are presented in Supplementary Figures S2 and S3.

Our phylogenetic reconstructions distributed PPKs into seven subfamilies, each being recovered in a well-supported clade (Figure 2), showing that the previous PPK classification into subfamilies (Zelle et al. 2013) is mostly in agreement with the evolutionary history of this family. The PPKs of *D. melanogaster* distributed as previously described Zelle et al., (2013), with the exception of *Dmelppk17*. Indeed, *ppk17* came out as the most divergent PPK and the current analysis suggests that it constitutes the sole member of a novel subfamily named Subf-VII (Figure 2, Supplementary Figures S2 and S4).

The second gene clade to split off the PPK tree was Subf-V (Figure 2). This subfamily presented many duplications and split into two main orthogroups (Supplementary Figure S5). One of these orthogroups included *ppk28*, *ppk8*, *ppk5*, and *ppk12*, while the other was composed by *ppk1* and *ppk2* and a large mosquito expansion.

Two main clades were identified in Subf-II, one including *ppk9* and *ppk10*, and the other including *ppk25* and *ppk3* (Figure 2; Supplementary Figures S2 and S6). Interestingly, one-to-one orthologs of *ppk9* were found for the different insect orders, the phylogenetic branching pattern resembling the phylogeny of the species studied here.

Three orthogroups were found for Subf-I, a first including the orthologs of *ppk15*, a second including those of *ppk13*, and a third group including both the orthologs of *ppk29* and *ppk31* (Figure 2 and Supplementary Figures S2 and S7). As shown by the phylogenetic tree, this subfamily exhibited a consistently dynamic evolutionary history along with duplications in different insect groups (Supplementary Figure S7). This contrasts with our findings for *Dmelppk23* orthologs forming Subf-VI in which less duplications or losses seemed to occur (Figure 2, Supplementary Figures S2 and S8).

Subf-III was basally subdivided into two main clades (Figure 2 and Supplementary Figures S2 and S9). The first included hemipteran genes, an expansion of 11 paralogs in *Bl. germanica* and one PPK of *Ca. floridanus* (the only Subf-III gene found for hymenopterans). The other orthogroup included several *D. melanogaster* PPKs *(Dmelppk7*, *Dmelppk14*, *Dmelppk19*, *Dmelppk20*, *Dmelppk21*, and *Dmelppk30*), their corresponding orthologs identified in *M. domestica* and *G. morsitans*, and representatives of other insects, mostly hemimetabolous. Notably, no Subf-III PPK was found across lepidopterans, mosquitoes, and beetles (Figure 3a).

**Figure 3.**
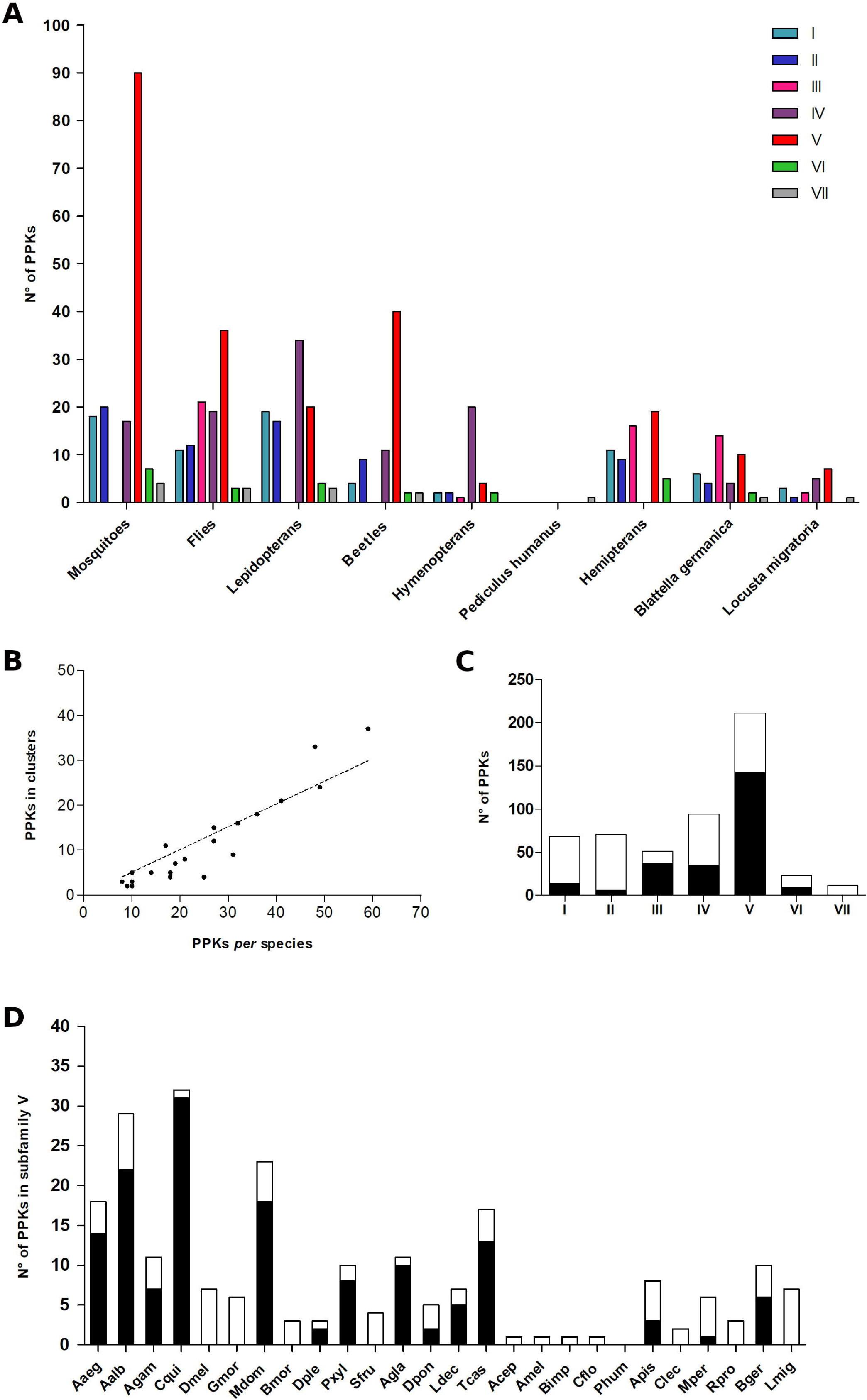
Number of PPKs *per* subfamily in the different taxonomic groups and species analyzed (A); Correlation between the number of PPKs in clusters and number of PPK *per* species (B); Number of clustered (black) and non-clustered (white) PPKs in each subfamily (C); Number of clustered (black) and non-clustered (white) PPKs in the analyzed species for Subf-V (D). Abbreviations are described in Table 2

Subf-IV encompassed 4 orthogroups and many lineage losses and duplications were identified (Figure 2 and Supplementary Figures S2 and S10). A first clade included orthologs of *ppk18*, *ppk27*, *ppk24* and *ppk22*, a second comprised those of *ppk4* and *ppk6*, a third included genes with no known orthologs in *D. melanogaster*, while a fourth clade included orthologs of *ppk11* and *ppk16.*

### PPK distribution along different insect orders

*Musca domestica* and *Bl. germanica* had the largest PPK repertoires, with 59 and 41 genes respectively, followed by mosquitoes (Table 1). Our analysis identified *Aaegppk29*, an additional gene to be included in the *Ae. aegypti* PPK repertoire (Matthews et al. 2019). In contrast, the lower end of the table included hymenopteran species that presented six to nine PPKs and *P. humanus*, for which only *ppk17* was found (Table 1).

**Table 1.**
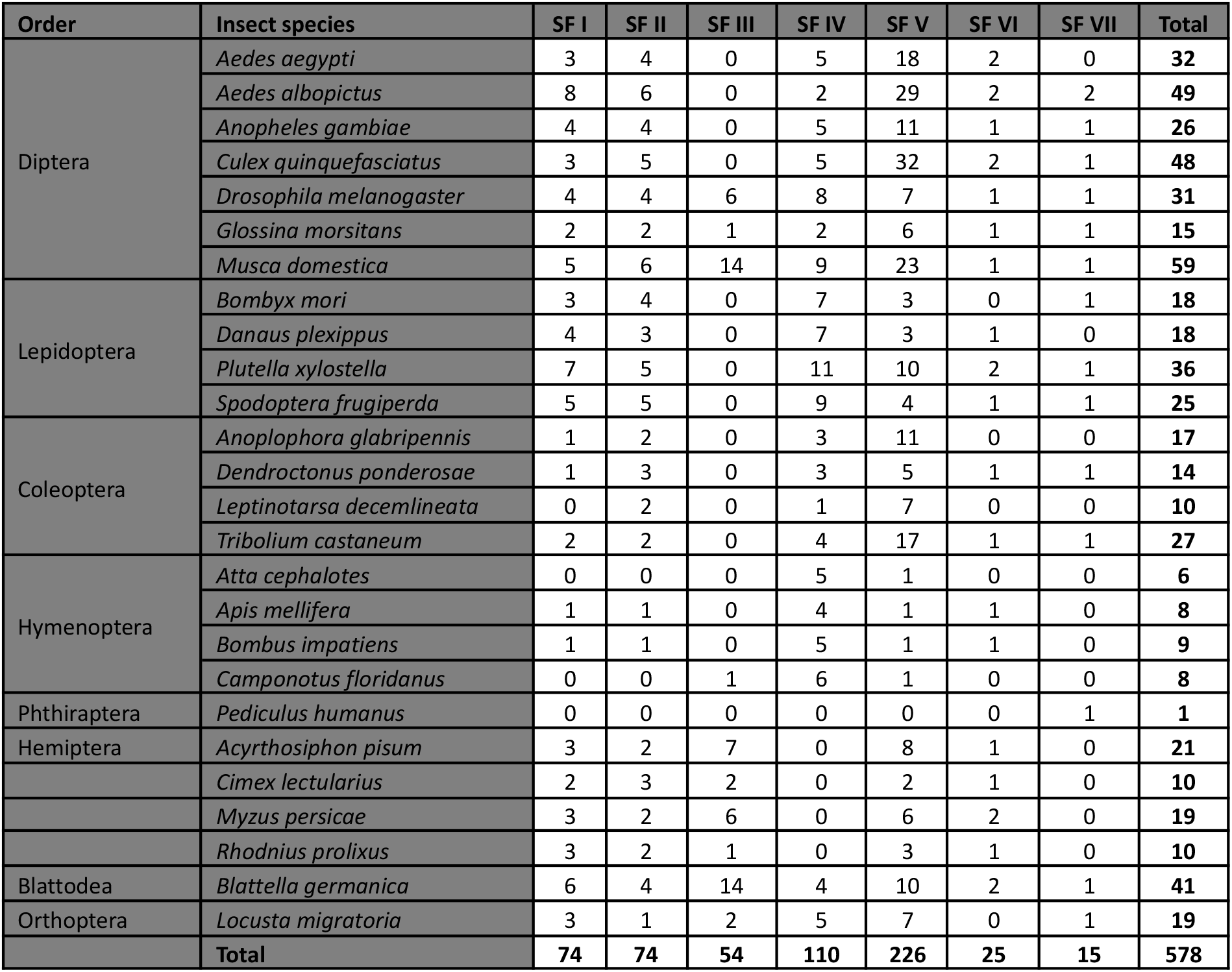
Composition of PPK subfamilies in the different insect orders analyzed. SF: Subfamily

**Table 2.** Number of PPKs identified in the different insect genomes and corresponding clustering patterns. (−) Families with any PPK; SF: Subfamily.

| Order | Insect species | PPKs | PPKs in clusters (%) | N° of clusters | Average of PPKs per cluster | N° of PPKs clustered per subfamily |  |  |  |  |  |  |
| --- | --- | --- | --- | --- | --- | --- | --- | --- | --- | --- | --- | --- |
|  |  |  |  |  |  | SF I | SF II | SF III | SF IV | SF V | SF VI | SF VI I |
| Diptera | *Aedes aegypti* | 32 | 16 (50%) | 5 | 3.2 | 0 | 0 | - | 0 | 14 (78%) | 2 (100%) | - |
|  | *Aedes albopictus* | 49 | 24 (49%) | 7 | 3.4 | 0 | 0 | - | 0 | 22 (76%) | 2 (100%) | 0 |
|  | *Anopheles gambiae* | 26 | 12 (46%) | 4 | 3 | 2 (50%) | 1 (20%) | - | 2 (40%) | 7 (64%) | 0 | 0 |
|  | *Culex quinquefasciatus* | 48 | 33 (69%) | 7 | 3.6 | 0 | 0 | - | 0 | 31 (97%) | 2 (100%) | 0 |
|  | *Drosophila melanogaster* | 31 | 9 (29%) | 3 | 2.9 | 0 | 0 | 6 (100%) | 3 (37%) | 0 | 0 | 0 |
|  | *Glossina morsitans* | 15 | 0 | 0 | 0 | 0 | 0 | 0 | 0 | 0 | 0 | 0 |
|  | *Musca domestica* | 59 | 37 (63%) | 11 | 3.4 | 0 | 0 | 12 (86%) | 7 (78%) | 18 (78%) | 0 | 0 |
| Lepidoptera | *Bombyx mori* | 18 | 5 (27%) | 2 | 2.5 | 2 (67%) | 0 | - | 5 (43%) | 0 | - | 0 |
|  | *Danaus plexippus* | 18 | 4 (22%) | 2 | 2 | 0 | 0 | - | 2 (29%) | 2 (67%) | 0 | - |
|  | *Plutella xylostella* | 36 | 18 (50%) | 9 | 2 | 2 (29%) | 0 | - | 7 (64%) | 8 (70%) | 1 (50%) | 0 |
|  | *Spodoptera frugiperda* | 25 | 4 (16%) | 2 | 2 | 3 (60%) | 0 | - | 1 (11%) | 0 | 0 | 0 |
| Coleoptera | *Anoplophora glabripennis* | 17 | 11 (65%) | 3 | 3.3 | 0 | 0 | - | 0 | 10 (90%) | - | - |
|  | *Dendroctonus ponderosae* | 14 | 5 (35%) | 2 | 2.5 | 0 | 0 | - | 3 (100%) | 2 (40%) | 0 | 0 |
|  | *Leptinotarsa decemlineata* | 10 | 5 (50%) | 2 | 2.4 | - | 0 | - | 0 | 5 (71%) | - | - |
|  | *Tribolium castaneum* | 27 | 15 (55.5%) | 3 | 5 | 0 | 0 | - | 2 (50%) | 13 (76%) | 0 | 0 |
| Hymenoptera | *Atta cephalotes* | 6 | 0 | 0 | 0 | - | - | - | 0 | 0 | - | - |
|  | *Apis mellifera* | 8 | 0 | 0 | 0 | 0 | 0 | - | 0 | 0 | 0 | - |
|  | *Bombus impatiens* | 9 | 2 (22%) | 1 | 2 | 0 | 0 | - | 2 (40%) | 0 | 0 | - |
|  | *Camponotus floridanus* | 8 | 3 (38%) | 1 | 3 | - | - | 0 | 3 (50%) | 0 | - | - |
| Phthiraptera | *Pediculus humanus* | 1 | 0 | 0 | - | - | - | - | - | - | - | 0 |
| Hemiptera | *Acyrtosiphon pisum* | 21 | 8 (38%) | 3 | 2.6 | 0 | 0 | 5 (71%) | - | 3 (37%) | 0 | - |
|  | *Cimex lectularius* | 10 | 3 (33%) | 1 | 3 | 1 (50%) | 2 (67%) | 0 | - | 0 | 0 | - |
|  | *Myzus persicae* | 19 | 7 (37%) | 3 | 2 | 1 (33%) | 2 (50%) | 3 (67%) | - | 1 (17%) | 0 | - |
|  | *Rhodnius prolixus* | 10 | 2 (20%) | 1 | 2 | 2 (67%) | 0 | - | 0 | 0 | 0 | - |
| Blattodea | *Blattella germanica* | 41 | 21 (51%) | 6 | 3.5 | 1 (17%) | 1 (25%) | 11 (79%) | 0 | 6 (60%) | 2 (100%) | 0 |
| Orthoptera | *Locusta migratoria* | 19 | 0 | 0 | - | 0 | 0 | 0 | 0 | 0 | - | 0 |

Subf-I and Subf-II genes were found in all insect orders studied, except for Phthiraptera (Figure 3a). Genes belonging to Subf-III were restricted to flies, hemipterans, the German cockroach, the migratory locust and the ant *Ca. floridanus* (Table 1 and Figure 3a). Interestingly, flies presented several duplications and one *M. domestica*-specific expansion including 6 paralogs which are orthologs to *ppk19*. Few one-to-one orthologs were identified across orders in Subf-IV (Figure 2), which presented no hemipteran members (Figure 3a). Presenting almost 40% of all PPKs analysed, Subf-V was the largest one, including a total of 226 genes (Table 1 and Figure 3a). This family included four lineages representing taxon-specific expansions. The first lineage included 37 beetle members, while the second corresponded to 85 mosquito members (*An. gambiae*: 10, *Ae. aegypti*: 17, *Ae. albopictus*: 27 and *Cx. quinquefasciatus*: 31). The third lineage included 13 genes belonging to hemipterans. Finally, the fourth lineage presented a *M. domestica* expansion related to *ppk2* that included 11 paralogs. Subf-VI was composed of a single orthogroup that includes *Dmelppk23* and the corresponding orthologs of 19 other insects (Table 1). The *ppk17* gene was the only member of Subf-VII and was absent in hymenopterans and hemipterans (Figure 3a).

### Genomic organization of PPKs in insect genomes

We examined the location of PPK genes along the different insect genomes and found a non-random distribution of these genes: 57% of all PPKs (329) shared a scaffold or chromosome with another PPK, most of which (86%) were closely related, i.e. members of the same subfamily (Supplementary Table S4). Next, we studied the distance between PPKs located in the same genomic region and observed that 70% of them were separated by less than 100,000 bp (Supplementary Table S4). This distance was used as a cut-off reference defining that two or more PPKs belonged to the same genomic cluster (Engsontia et al. 2014). According to this criterion, the PPKs of *G. morsitans, At. cephalotes, Ap. mellifera, L. migratoria,* and *P. humanus* were not arranged in clusters and thus, were not considered in subsequent analyses. Ten out of 21 remaining insect genomes presented more than 40% of their PPKs into clusters (Table 3). Among them, mosquitoes, the house fly, the cockroach and the moth *Pl. xylostella* exhibited a higher number of clusters (between 5 to 11 genes *per* cluster) in comparison with the rest of the genomes analyzed. A positive correlation was observed between the number of PPKs and the number of genes in clusters (Spearman R=0.91, P<0.0001, Figure 3b).

**Table 3.** Gene birth and death rate estimates for the entire PPK family and for each subfamily.

| Subfamily | “Total time” rates |  | “Branch average” rates |  |
| --- | --- | --- | --- | --- |
|  | Birth | Death | Birth | Death |
| Subf-I | 0.0008 | 0.0016 | 0.0006 | 0.0007 |
| Subf-II | 0.0007 | 0.0011 | 0.0019 | 0.0014 |
| Subf-III | 0.0028 | 0.0026 | 0.0059 | 0.0040 |
| Subf-IV | 0.0027 | 0.0016 | 0.0083 | 0.0028 |
| Subf-V | 0.0069 | 0.0011 | 0.0102 | 0.0018 |
| Subf-VI | 0.0008 | 0.0008 | 0.0021 | 0.0011 |
| Subf-VII | 0.0004 | 0.0016 | 0.0015 | 0.0029 |
| All | 0.0024 | 0.0015 | 0.0047 | 0.0020 |

Most clusters were formed by genes of the same subfamily (72 out of 80), while the remaining eight presented genes coming from more than one subfamily (Supplementary Table S4). Most genes belonging to Subf-III and Subf-V were distributed in clusters (73% and 67%, respectively) (Figure 3c). This high degree of clustering is explained by gene expansions seen in several taxonomic groups for both subfamilies, e.g. in *M. domestica* and *Bl. germanica* for Subf-III (Supplementary Table S4), and in mosquitoes, beetles and hemipterans for Subf-V (Figure 3d). A second group including Subf-IV and Subf-VI presented an intermediate clustering profile (40%). Finally, most of the PPKs belonging to Subf-I and Subf-II were not included in clusters (21% and 9% of PPKs, respectively).

### Family size dynamics

The PPK gene family evolved according to a birth and death evolutionary model (Nei and Rooney 2005). Overall, the PPK family is expanding, as inferred from the comparison of birth and death rates (Table 3). Furthermore, Subf-III, Subf-IV, and Subf-V were the most dynamic, presenting birth/death rates compatible with gene expansion (Table 3). Although Subf-II and Subf-VI had different rates depending on the estimation methods, their birth and death rates were always similar to each other, suggesting that their sizes are stable across the groups analyzed (Table 3). On the other hand, Subf-I and Subf-VII appeared to be contracting, with the latter contracting at a faster rate (Table 3).

Higher rate estimates obtained with the “branch average” method reflected the multiple duplication and loss events that occurred in short branches of the phylogeny, i.e. within a specific order or taxonomic group, as observed for the expansions in *M. domestica* (Figure 4).

**Figure 4.**
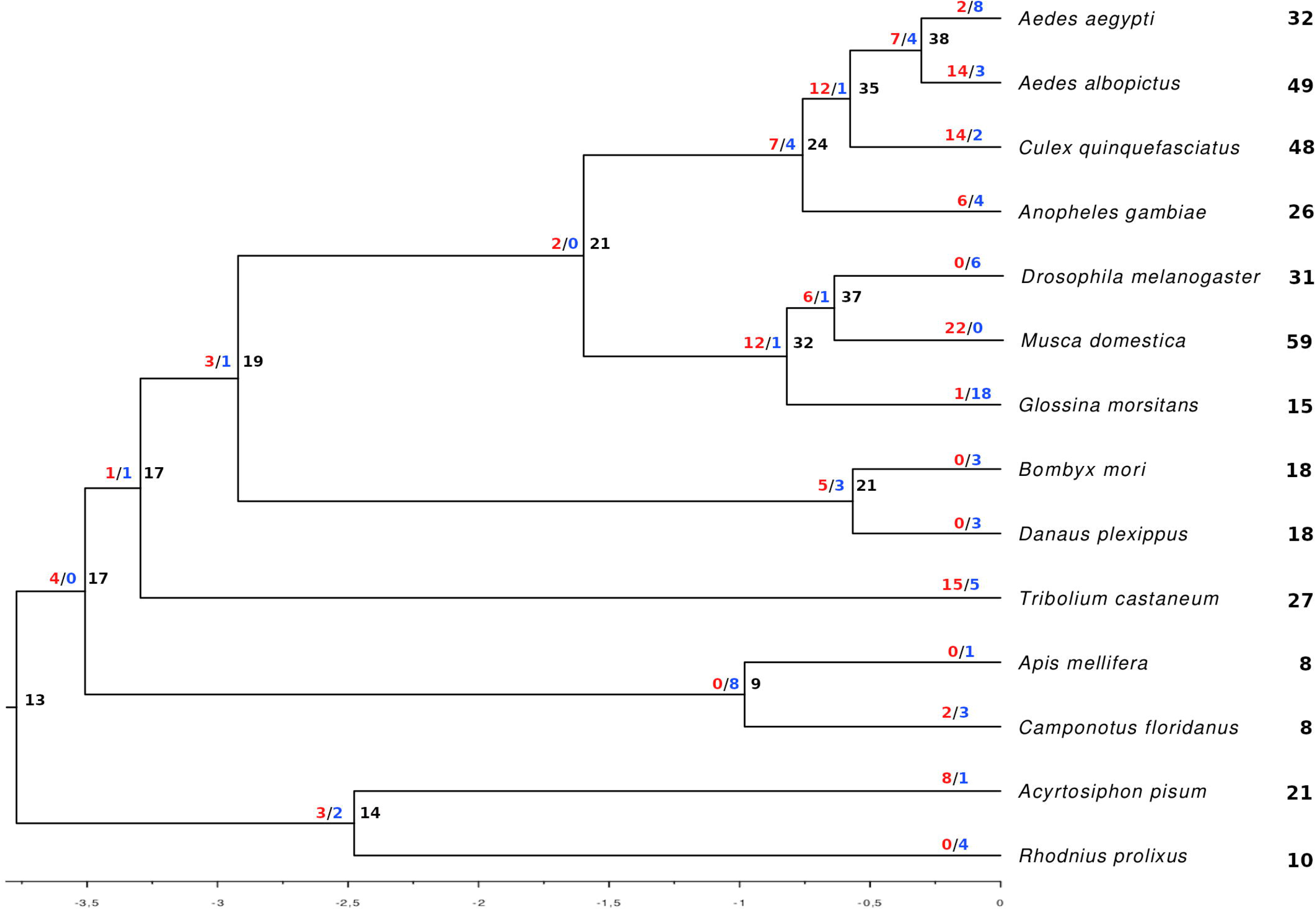
Species tree showing the number of gene gains (blue) and losses (red) in each branch and the total number of genes for each terminal.

### Selective pressure analysis

A total of 20 orthogroups, i.e. those composed by orthologs of 16 *D. melanogaster* PPKs and 4 taxon-specific expansions, were included in the analysis of selective pressure. The role of natural selection in the evolution of these 20 orthogroups was evaluated by means of a phylogeny based analysis (Supplementary Table S5). The ω-values calculated by means of the M0 were lower than 0.16, except for the hemipteran and *Bl. germanica* orthogroups, which exhibited ω-values of 0.20 and 0.21, respectively. Interestingly, the *ppk17* orthogroup, the most divergent member of the PPK family, presented the lowest ω = 0.01.

Only three genes (*ppk6*, *ppk25* and *ppk27*) were found to be under positive selection (M7 *vs.* M8 model comparison, Supplementary Table S5). The *ppk6* orthogroup showed two predicted codons in the C-terminal region with posterior probabilities higher than 0.95 of being under positive selection. The *ppk25* orthogroup presented three codons under positive selection: 1 codon at the N-terminal region (329) and 2 codons (565 and 567) at the extracellular loop (i.e. 11 residues downstream of the “G-X-C-X-X-F-N” motif). Finally, the *ppk27* orthogroup showed three positions under positive selection that were located at the N-terminal region before the first transmembrane domain.

## DISCUSSION

We have performed a large-scale genomic data mining analysis to inquire into the evolution of the PPK gene family across 26 species spanning four hemimetabolous and four holometabolous insect orders. PPKs were detected in all genomes studied and assigned to 7 subfamilies, including the new Subf-VII formed by orthologs of the *ppk17* gene that seems to be the basal PPK family member. Our analysis shows that PPKs evolved under a birth-and-death model that generated lineage-specific expansions by gene duplication events and the effect of strong purifying selection. The size of PPK repertoires displayed high variability, while *M. domestica, Bl. germanica* and several mosquito species presented the largest gene sets, *P. humanus*, bees and ants had very small ones. Whether these important differences relate to specific ecological needs and constraints of these insects, or to clade-specific genetic properties favoring or limiting gene duplication events is not clarified by our analyses. Subfamily V is revealed as the largest and most dynamic one, presenting a specific mosquito expansion that deserves attention as a source of potential targets for pest control. Species-specific expansions identified in *Bl. germanica* and *M. domestica* also show promise for developing specific tools for controlling these nuisance insects. Our phylogeny displayed a set of conserved PPKs across insect genomes suggesting a very conserved function that resembles the case of antennal ionotropic receptors. The new highly conserved residues found in PPK sequences, especially for the second transmembrane domain, indicate that they possess additional fundamental roles critical for in receptor function. The calmodulin binding motifs reported for more than a hundred PPKs suggest that a relevant proportion of the members of this family may modulate sensory neuron responses as proposed by Ng. et al. (2019).

### *Ppk17* formed a new and divergent PPK subfamily

Our phylogenetic analysis revealed that the PPK family is composed of seven subfamilies, each one located in well-supported clades (Figure 2). In the case of the *ppk17* orthogroup, our analysis indicated that it is an independent lineage within the PPK family and should not be included in Subf-V as previously suggested (Zelle et al. 2013). Furthermore, the clade formed by proteins of the *ppk17* orthogroup represented the most divergent member of the PPK family (Figure 2 and Supplementary Figures S2 and S3). George et al., (2019) used loss-of-function mutant and RNAi lines to study the role of ion channels in wing development in *D. melanogaster* and found that *Dmelppk17*, *Dmelppk1*, *Dmelppk2*, *Dmelppk25,* and *Dmelppk30*, among others, are likely involved in this pathway. Interestingly, *ppk17* was identified in all insect orders, except for hymenopterans, that apparently lost it (Figure 2 and Table 1). This gene presented the strongest level of purifying selection (ω-value = 0.01), similar to that reported for the odorant receptor co-receptor, *OrCo* (Soffan et al. 2018), suggesting that its function is likely conserved and fundamental for insects. Functional genetic studies in other experimental models are necessary to confirm this hypothesis and provide further details on its function.

### Genetic mechanisms driving PPK evolution

Detailed examination of the relationships of PPKs across several insect orders revealed the expected evolutionary pattern of birth-and-death typical of environmentally relevant genes (Nei and Rooney 2005; McBride and Arguello 2007; Vieira and Rozas 2011; Almeida et al. 2014; Sánchez-Gracia et al. 2011), with lineage-specific expansions (through duplication) and contractions (through pseudogenization). Consistently with this evolution model, 329 PPKs (57%) shared the same scaffold or chromosome, 284 being of the same subfamily, and 230 located in clusters. This pattern of related genes located in nearby chromosomal regions suggests that they are products of relatively recent gene duplications. A similar distribution pattern was previously reported for ORs or GRs (Robertson et al. 2003; Engsontia et al. 2008; Wanner and Robertson 2008; Smadja et al. 2009). Gene duplication is fundamental for the diversification of chemoreceptor repertoires as it can be followed by neofunctionalization or subfunctionalization processes leading to evolutionary innovation (Sánchez-Gracia et al. 2009). It has been observed that genes within clusters, like ORs and GRs, are usually co-regulated, which can lead to joint gene expression (Robertson et al. 2003; Nozawa and Nei 2007; Guo and Kim 2007). Therefore, we would expect that PPKs follow a similar pattern: clusters might be subject to common regulatory mechanisms and deal with common or related stimuli.

Estimates of gene birth and death rates suggest that the PPK family is generally expanding in the set of species we analyzed, although it could be contracting in particular species or groups. No estimate of birth and death rates were available across insect orders, therefore we used estimates obtained for the *Drosophila* genus for comparisons (see Sánchez-Gracia et al., (2009) and references therein). It is important to acknowledge that our gene birth and death rates might be underestimated due to lack of information on pseudogenes and because we used a conservative approach when counting duplication and loss events by gene tree *vs.* species tree reconciliation. The “total time” method was our choice for all comparisons among families discussed herein, because it was the method used to obtain previous estimates for other chemosensory families (Sánchez-Gracia et al. 2009), even though it may lead to underestimates. The “branch average” method estimated higher rates, reflecting the availability of genomes of closely related species (i.e. flies, mosquitos) and the large number of duplication and loss events in these groups, as it takes into account each branch length (in units of time). This last approach is more accurate, but also more dependent on the species set used in analysis. Our rate estimates for the PPK family as a whole are in the lower range of rates obtained for other chemosensory gene families, but yet it is substantially higher than the genomic average. Nevertheless, there was considerable variation in gene birth and death rate estimates among PPK subfamilies (Table 3). While some had very low rates (e.g. Subf-II and Subf-VI), others had rates comparable to those of the most dynamic chemosensory gene families, the ORs and GRs (e.g. Subf-V).

Despite the differences in turnover rates and gene repertoires, many PPK subfamilies were represented in most insect orders (Table 2). PPK subfamilies presented different evolutionary patterns, probably as a consequence of different selection pressures. For example, Subf-III was among the most dynamic, together with Subf-IV and V, and it is almost exclusive of flies and hemimetabolous insects (Table 2). The available functional information, which is restricted to few *D. melanogaster* members, and the absence of one-to-one orthology between fly PPKs and other Subf-III members impedes understanding why this subfamily was lost in several insect orders. Subf-IV also had high estimates of gene turnover rates (Table 3), but with a larger difference between birth and death rates pointing to an expansion trend of this subfamily. It had 110 members identified in all orders and was only exceeded in size by Subf-V, the most dynamic and expanded subfamily with almost half of the PPKs. Subfamilies VI and VII presented an opposite pattern, as both were conserved across different insect orders, with few gene duplication events. This was reflected in their low turnover rates, as both were restricted to one or two members *per* species in the ortholog groups represented by *ppk23* and *ppk17*, respectively (Table 1 and 4).

PPK evolution is mostly driven by purifying selection, as ω-values estimated for selected orthogroups were generally low. Besides, the presence of positively selected sites was very limited, only identified in three orthogroups without observing a common distribution pattern along the protein sequences (Supplementary Table S5). As the orthogroups included followed two criteria (conforming an expansion or having one *D. melanogaster* PPK to grant functional data), we cannot discard that excluded PPK lineages could resemble this pattern. Estimated ω-values were low even when we compared genes within an expansion, contrary to the suggestion that purifying selection is relaxed after duplication events. Future analyses, including more closely related species and specific gene lineages belonging to the expansions described for Subf-V, may clarify which evolutionary forces shaped PPK evolution. Re-annotations of several insect genomes will allow identifying additional PPKs, as seen with *Ae. aegypti* (Matthews et al. 2019). Besides, improving the annotation of the PPKs identified herein by means of RNA-Seq and performing subsequent functional experiments, as well as extending the analysis to other arthropods, would help further characterizing the PPK family. Indeed, preliminary searches using the Deg/ENaC PFAM domain on several arthropod genomes identified a set of candidate PPK sequences, suggesting that it is possible that the PPK family is not exclusive of insects. Further work under development, including new searches on non-arthropod genomes followed by sequence and phylogenetic analyses will evaluate the origin, as well as the complete evolutionary history of PPKs.

### Subf-V represents the largest gene lineage

Sánchez-Gracia et al., (2009) reported the turnover rates of diverse chemosensory gene families among which GRs were the only showing a turnover rate higher than that seen for Subf-V in our study (Table 3). Three large expansions made it the largest subfamily (226 genes), including almost half of the PPKs identified in this study (Figure 2). A similar pattern was described for the 9-exon subfamily of ant ORs that is extremely expanded (Engsontia et al. 2015). The expansions of Subf-V sat into genomic clusters (Supplementary Table S3) and were taxon-specific: flies, mosquitoes, beetles and hemipterans (Figure 2). The origin of these expansions may have been a response to rapid adaptation to changing chemical environments, as suggested for other sensory receptor families (Smadja et al. 2009; Engsontia et al. 2015). However, specific information about cognate ligands or function is lacking for these PPKs, limiting our capacity to understand the factors causing these expansions.

In the case of mosquitoes, the increase in the number of PPKs is mainly due to the Subf-V expansion that is related to the *ppk1-ppk2-ppk26* clade (Figure 2). The functional information available for this set of PPKs is restricted to *D. melanogaster* larvae (see Supplementary Table S6), where they are co-expressed mediating mechanosensation, acid sensing (pH between 5 and 9), and proper locomotion (Adams et al. 1998; Zelle et al. 2013; Gorczyca et al. 2014; Boiko et al. 2012; Tsubouchi et al. 2012) Whether this expansion was driven to mediate mosquito pH sensing in aquatic larval environments deserves experimental assessment. Tissue-specific expression analyses in larvae and adults (through RNA-Seq or *in situ* hybridization) and functional genetic studies should clarify the role of this mosquito lineage. In the end, this large exclusive lineage emerges as a potential target to manipulate mosquito behaviour through genetic tools.

Two other lineages belonging to Subf-V represent important expansions deserving attention. These are the beetle and hemipteran-specific expansions that include a large set of genes each. No functional information from orthologs is available to interpret their potential roles, as both are composed of gene sets that seem exclusive of insects of those orders for which no experiments tested functional hypotheses regarding PPKs. In spite of this, these large, order-specific expansions seem appealing for developing functional analyses intending to search for targets directed to control pests belonging to these insect orders.

### Extreme variations in PPK repertoires

Consistently with their sensory role and fast evolution, the size and composition of insect PPK repertoires was shown to be very diverse, although phylogenetic proximity is associated with consistently similar repertoire sizes (Table 1). The impressive species diversity that insects have reached through their evolution shows a high capacity to adapt according to environmental heterogeneities and changing scenarios. In this sense, the dramatically different PPK repertoires reported here may reflect specific sensory abilities shaped by the particular niches where these animals thrive (McBride and Arguello 2007; Smadja et al. 2012; Sánchez-Gracia et al. 2011; Robertson and Wanner 2006). The extreme examples of 59 PPK family members found in the genome of *M. domestica*, and only one in that of *P. humanus* capture the extent of the plasticity shown by these gene repertoires to adapt to environmental pressures. In the case of the human lice, the reduction observed for the PPK family follows the pattern seen for other families of sensory receptors (only 10 ORs, 8 GRs, and 12 IRs), which is assumed to be related with a very limited foraging range (Kirkness et al. 2010). The hymenopteran species analyzed herein are another example showing relatively small PPK sets in their genomes (6-9 genes), in coincidence with those observed for their GR and IR families (Supplementary Table S7). On the other hand, the large OR sets known for hymenopterans suggest that they rely mostly on olfaction to detect and discriminate floral/plant/prey-emitted compounds (Robertson and Wanner 2006; Zhou et al. 2015).

Dipterans, except for *G. morsitans* that had a global reduction of sensory receptor families, had large PPK repertoires (Table 1). Interestingly, *Musca domestica* (59) had the largest PPK gene set of all insects analyzed, clearly overcoming other fly species: *G. morsitans* (15) and *D. melanogaster* (31). This receptor diversity is similar for other chemosensory families for which *M. domestica* also has specific expansions (Scott et al. 2014), suggesting that the house fly is an extreme example of chemosensory environmental sounding. Interestingly, its IR and GR expansions were most frequent for orthologs of *D. melanogaster* genes mediating the detection aversive tastants, and reported to facilitate its adaptation to septic environments (Scott et al. 2014). A similar case can be considered for *Bl. germanica*, which presented the second largest set of PPKs and is known to possess the largest sets of IRs and GRs described to date (Robertson et al. 2018). Therefore, this insect is another example of extreme chemosensory abilities developed to sound the chemical complexity of the environment. Whether these extreme cases were driven during the development of their synanthropy deserves consideration. Indeed, anthropic environments offer a broad diversity of nutrient sources, as well as toxins, and might have driven the development of these highly diversified chemosensory repertoires necessary to tell apart and exploit the complex features of anthropic chemistry, gaining access to the resources gathered by the most cosmopolitan of animal species.

A total of 41 PPK genes belonging to all subfamilies were identified in *B. germanica*, the largest repertoire among those of non-dipteran insects. This insect is an extreme omnivore (Schal et al. 1984) with an immense sensory gene repertoire (Supplementary Table S7, GRs= 545; IRs= 897) (Robertson et al. 2018). Indeed, this species shows remarkably high rates of gene turnover in several families, including chemoreceptors, xenobiotic defense proteins, and metabolism proteins, which has been correlated to its capacity of rapid adaptation to new environments (Thomas et al. 2018). Therefore, the enlarged set of PPKs identified in *Bl. germanica* could also contribute to this ability to exploit any available sources of nutrients and survive in a large diversity of environments.

Expanded species-specific lineages for *Bl. germanica* and *M. domestica* were identified in Subf-III and Subf-V. In the case of the Subf-III, two expansions were identified for *Bl. germanica* (11 members) and another for *M. domestica* (6 members). The latter is related to *Dmelppk19*, which has been shown to be involved in sensing low salt concentrations, acting together with *Dmelppk11* (Liu et al. 2003). The expansion of this PPK lineage would probably increase house fly sensitivity or capacity to detect different types of salts or to discriminate concentration ranges. The position of the *Bl. germanica* expansion in the tree depicting Subf-III prevents proposing a putative function for this lineage due to their lack of orthology with *D. melanogaster* PPKs. A *M. domestica* expansion including 11 paralogues related to *ppk2,* a gene mediating gentle touch in *D. melanogaster* larvae, was identified in Subf-V (Supplementary Table S6). Future studies should evaluate whether this expansion provides enhanced mechanosensory abilities to *M. domestica* larvae (Tsubouchi et al. 2012). In sum, these species-specific expansions represent new specific targets to control these highly synatropic insects.

### A subset of conserved PPKs

Functional and structural studies of DEG/ENaC channels have shown that these receptors probably act as hetero or homotrimetric complexes for which subunit composition determines function and pharmacological properties (Ben-Shahar 2011). Indeed, expression and functional experiments with *D. melanogaster* PPKs indicate that they may also function as complexes, e.g., *ppk23* is co-expressed with *ppk29* and *ppk25* in sensory organs mediating cuticular hydrocarbon pheromone detection (Supplementary Table S6).

Our data mining analysis evinced a pattern of prevalent *vs* divergent PPKs that could be useful to understand their functional properties. We observed that most PPKs are divergent, impeding to establish a one-to-one orthology across insect orders. This is similar to the patterns observed for OR, OBP, and GR gene families, for which most genes present low degrees of orthology. In some of these gene families, a few known exceptions, such as the odorant receptor co-receptor (*OrCo*) and the GRs involved in carbon dioxide detection, are highly conserved (Jones et al. 2007; Larsson et al. 2004; Jones et al. 2005; Robertson and Kent 2009). Coincidently, several PPK orthogroups were prevalent among the insect orders studied here. Members of the *ppk28* orthogroup were identified in 92% of the genomes studied (Figure 2). This gene was only absent in *P. humanus,* and *Le. decemlineata.* It is possible that the latter case was due to an error in the automated gene prediction, as *ppk28* orthologs have been identified for the other beetle genomes. *Ppk28* has been reported to mediate water detection in *D. melanogaster*, while it is also reported to mediate responses to water and low salt concentrations in *Ae. aegypti* during egg-laying initiation (Supplementary Table S6). Water sensing could indeed be considered a mechanism detecting osmolarity changes in the environment, a fundamental function that could explain why *ppk28* has been conserved through the evolution of most insects. Functional genetic studies in other insect species are lacking and would be necessary to confirm the role of *ppk28* as an osmolarity sensor in non-dipteran insects.

Another prevalent gene, *ppk23* is co-expressed with *ppk29* and *ppk25* in the antennae, proboscis and legs of *D. melanogaster*, where this complex acts as a contact chemoreceptor of cuticular hydrocarbon pheromones controlling the sexual behavior of both sexes (Supplementary Table S6). Cuticular hydrocarbons mediate sexual, colony recognition, and aggregation behaviors in diverse insect orders (Pavković-Lučić et al. 2012), suggesting that the pheromone receptor role seen for *D. melanogaster* can be potentially found in other cases. *Dmelppk23* is also expressed in GRNs from the labellum, where, together with *Gr64f*, *Gr66a*, *Ir94e*, *Ir76b*, and *ppk28*, mediates salt taste (Jaeger et al. 2018). It has been proposed that the GRs provide a baseline level of salt attraction or avoidance, whereas *Dmelppk23* adjusts insect responses depending on internal stage (Jaeger et al. 2018). Considering the importance of salt taste for feeding behavior (Jaeger et al. 2018; Pontes et al. 2017) and the presence of *ppk23* in diverse insect orders (Figure 2), it is possible that its functionalities have been maintained throughout insect evolution.

Notwithstanding the fact that *ppk9* was as prevalent as *ppk23* (both present in 73% of the genomes analyzed), raising functional hypotheses is not feasible because its role is unknown to date. Expression of this gene has been detected in several tissues of larvae and adults, including different sections of the digestive tract, salivary glands, and fat body of *D. melanogaster* (Zelle et al. 2013). A fourth example of prevalence is *Dmelppk16* (65% of the genomes had orthologs for this gene), which was reported to mediate presynaptic homeostatic plasticity together with *Dmelppk1* and *Dmelppk11* (Younger et al. 2013; Orr et al. 2017). However, it is difficult to extrapolate a prevalent role for *ppk16* in other insects because *ppk1* and *ppk11* are exclusive of flies. *Dmelppk13* (58% of the genomes had orthologs for this gene) was found to be expressed in hairs of legs, tarsi and wings, suggesting that this receptor has a role in gustation (Liu et al. 2003). Finally, *Dmelppk17*, orthologs of which were detected in the genomes of 14 species (54%), was reported to be involved in wing morphogenesis (George et al. 2019). Interestingly, *ppk17* is the only gene of this family present in the genome of wingless lice. Whether its conserved role relates to that reported for *D. melanogaster* does not seem probable based on this observation. This pattern of conserved *vs.* divergent PPKs resembles the characteristic division between antennal and divergent receptors known for the IR family (Croset et al. 2010). Expression data and functional experiments, especially studying conserved PPKs, should clarify whether these features are related to a particular functional architecture that governs this protein family.

### Conserved features of PPK sequences

The degenerin site (DEG or *d* position) is a residue located in the segment that precedes the second transmembrane domain and plays a fundamental role in the open state and ion selectivity of the channel (Eastwood and Goodman 2012). This position is most commonly occupied by glycine, alanine or serine in DEG/ENaC proteins (Eastwood and Goodman 2012). In the PPK sequences analyzed herein, alanine and serine also occupied the DEG site, 180 and 79 sequences being observed in this position, respectively. Interestingly, valine, a bulkier residue, was identified in the DEG site in 137 PPK sequences. This PPK feature could have functional implications, as the presence of an amino acid with bigger side chains in the DEG site has been implicated in changes in channel activity of other DEG/ENaC superfamily members (Eastwood and Goodman 2012). Furthermore, a substitution of serine by phenylalanine increased the activity of the DEG site of *Dmelppk1* (Boiko et al. 2012). Residues present in the DEG site varied across PPK subfamilies (Supplementary Table S2), suggesting that they may affect channel activity and functionality.

Liu et al., 2003) and Zelle et al., (2013) reported conserved features in PPK sequences of several *Drosophila* species and *An. gambiae* that according to our analysis are conserved in insects in general. First, the cysteine-rich domain, which seems fundamental in mammals for an efficient transport of assembled channels to the cell membrane (Firsovt et al. 1999) is highly conserved (77% of sequences had more than 10 cysteine residues). Second, a tryptophan residue in the first transmembrane domain that is critical to Na^+^ sensitivity was detected in 66% of the PPK sequences (Figure 1; Pochynyuk et al., (2009). Finally, the “GxS” motif of the second transmembrane domain (Figure 1), which contributes to the channel pore and is fundamental for ion selectivity and surface expression (Eastwood and Goodman 2012), was identified in 68% of the sequences. Our analysis revealed a higher conservation of the second transmembrane domain compared to the first one (Figure 1), which is consistent with its fundamental role in pore formation in DEG/ENaCs (Hong and Driscoll 1994). Besides, new highly conserved residues, a leucine in position 5 and a glutamic acid in position 16, were identified in the second transmembrane domain (Figure 1), suggesting that they may contribute to pore function.

Recently, Ng et al., (2019) have described the involvement of *Dmelppk25* in the amplification of olfactory responses in *D. melanogaster*. These authors reported that after odor stimulation of *Or47b* and *Ir84a*, Ca^2+^ influx increases and activates *Dmelppk25* via an intracellular Calmodulin-Binding Motif (CBM). They identified CBMs in other *D. melanogaster* PPKs, suggesting that they function as Ca^2+^-activated amplification channels. Our results revealed CBMs in more than a hundred PPKs from other insect species (Supplementary Table S3), reinforcing the hypothesis that these receptors may amplify sensory responses through Ca^2+^ activation.

## CONCLUSIONS

Our phylogenetic and evolutionary characterization of PPKs across different insect orders has allowed identifying specificities that turn many of these genes into potential candidates to manipulate insect behavior and physiology. First, PPKs control fundamental insect physiological processes, like water or salt detection. Second, they are particularly expanded for several synanthropic insects, including *M. domestica* and *Bl. germanica*, suggesting that these receptors have been relevant for their adaptation to anthropic habitats. Third, the large culicid expansion observed for PPK Subf-V suggests an important role that grants them as very specific targets for developing specific mosquito control tools. Finally, the identification of CBMs in PPK sequences suggests that a large subset of these receptors may be related to the amplification of sensory responses, and tools targeting them could affect a diversity of sensory modalities.

## Authors’ contributions

J.M.L.E, F.C.A. and M.G.L conceived the project and designed the experiments. J.M.L.E and F.C.A performed data analyses. All authors wrote the manuscript and provided comments on versions, read and approved the final manuscript.

## Acknowledgments

The authors wish to thank the Program for Technological Development in Tools for Health-PDTIS-FIOCRUZ for having facilitated the use of its facilities.

## Supplementary information

**Table S1. Source of peptide datasets used to identify PPK gene candidates in the insect genomes.**

**Table S2. Analysis of PPK sequences.** Columns 9-15 display the presence of conserved residues and motifs described by Liu et al. (2003) for the PPK sequences of *D. melanogaster*. SF: Subfamily, Aa: amino acids; TM: transmembrane domains; Cs: cysteines; DEG: degenerin residue; “x” indicates any residue; “h” indicates a hydrophobic residue; Abs: Absent. Length is expressed in number of amino acids.

**Table S3. Calmodulin Binding Motif (CBM) analysis.** SF: Subfamily; TM: transmembrane domains. Length is expressed in number of amino acids.

**Table S4. Genomic location of PPK genes and cluster characteristics.** PPKs located in clusters are depicted in red. Exp: Expansion.

**Table S5. Results of likelihood ratio test and parameter estimates under the best-fitting model for each orthogroup.** ω-values were obtained with the M0 model. LR = Likelihood ratio (2ΔL). Estimated parameters were obtained under M8 model (when significant): P_0_ = proportion of sites that follow a beta distribution with 10 omega classes (0 ≤ ω ≥ 1); P_1_ = proportion of sites in the extra class with ω ≥ 1; location of predicted sites under positive selection and their posterior probability in parentheses.

**Tabla S6. Functional information reported for PPKs**

**Table S7. Number of sensory receptor families from different insect species.** Data of OR, GR and IR genes from *Myzus persicae* are not available. ND: No data,

**Figure S1. Alignment of Calmodulin Binding Motifs from the PPK subfamilies.**

**Figure S2. Molecular phylogenetic analysis of PPK genes across different insect orders.** The evolutionary history of PPK genes is based on a MAFFT alignment and constructed on IQ-Tree using the WAG+F+R9 (Best-fit model according to Bayesian Information Criterion) as a model of amino-acid substitution. Maximum-likelihood tree is based on 1,000 replicates and branch support values were estimated using the approximate Likelihood Ratio Test based on the Shimodaira-Hasegawa (aLRT-SH) and the ultrafast bootstrap or UFBoot procedures (Hordijk and Gascuel 2005; Minh et al. 2013). The tree was rooted on three amiloride-sensitive sodium channels (SCNN) from *Rattus norvegicus*, which were used as an outgroup. The PPK sequences from *D. melanogaster* were displayed in blue. Species abbreviations: Aaeg: *Aedes aegypti*, Aalb: *Aedes albopictus*, Agam: *Anopheles gambiae,* Cqui: *Culex quinquefasciatus,* Dmel: *Drosophila melanogaster,* Gmor: *Glossina morsitans,* Mdmo: *Musca domestica*, Bmor: *Bombyx mori,* Dplex: *Danaus plexippus*, Pxyl: *Plutella xylostella*, Sfru: *Spodoptera frugiperda*, Agla: *Anoplophora glabripennis;* Dpon: *Dendroctonus ponderosae,* Ldec: *Leptinotarsa decemlineata*, Tcas: *Tribolium castaneum*, Amel: *Apis mellifera*, Acep: *Atta cephalotes*, Bimp: *Bombus impatiens*, Cflo: *Camponotus floridanus,* Phum: *Pediculus humanus*, Apis: *Acyrthosiphon pisum,* Clec: *Cimex lectularius*, Mper: *Myzus persicae*, Rpro: *Rhodnius prolixus*, Bger: *Blattella germanica* and Lmig: *Locusta migratoria.* The sequences used are presented in Supplementary Data File S1.

**Figure S3. Molecular phylogenetic analysis of PPK genes across different insect orders.** The Maximum-likelihood tree is based on a MAFFT alignment and was constructed using RAxML v8 (Stamatakis 2014). Phylogenetic tree was generated with the PROTCATWAG amino-acid substitution model for tree search step and 200 replicates in the bootstrapping analysis. The tree was rooted on three amiloride-sensitive sodium channels (SCNN) from *Rattus norvegicus*, which were used as an outgroup. The PPK sequences from *D. melanogaster* were displayed in blue. Species abbreviations: Aaeg: *Aedes aegypti*, Aalb: *Aedes albopictus*, Agam: *Anopheles gambiae,* Cqui: *Culex quinquefasciatus,* Dmel: *Drosophila melanogaster,* Gmor: *Glossina morsitans,* Mdmo: *Musca domestica*, Bmor: *Bombyx mori,* Dplex: *Danaus plexippus*, Pxyl: *Plutella xylostella*, Sfru: *Spodoptera frugiperda*, Agla: *Anoplophora glabripennis;* Dpon: *Dendroctonus ponderosae,* Ldec: *Leptinotarsa decemlineata*, Tcas: *Tribolium castaneum*, Amel: *Apis mellifera*, Acep: *Atta cephalotes*, Bimp: *Bombus impatiens*, Cflo: *Camponotus floridanus,* Phum: *Pediculus humanus*, Apis: *Acyrthosiphon pisum,* Clec: *Cimex lectularius*, Mper: *Myzus persicae*, Rpro: *Rhodnius prolixus*, Bger: *Blattella germanica* and Lmig: *Locusta migratoria.* The sequences used are presented in Supplementary Data File S1.

**Figure S4. Phylogenetic tree for Subf-VII.** It was generated using RAxML with 20 independent runs.

**Figure S5. Phylogenetic tree for Subf-V.** It was generated using RAxML with 20 independent runs,

**Figure S6. Phylogenetic tree for Subf-II.** It was generated using RAxML with 20 independent runs.

**Figure S7. Phylogenetic tree for Subf-I.** It was generated using RAxML with 20 independent runs.

**Figure S8. Phylogenetic tree for Subf-VI.** It was generated using RAxML with 20 independent runs.

**Figure S9. Phylogenetic tree for Subf-III.** It was generated using RAxML with 20 independent runs.

**Figure S10. Phylogenetic tree for Subf-IV.** It was generated using RAxML with 20 independent runs.

**Data file S1. PPK sequences from target insects in fasta format**

