## Supplementary tables for "Evolution of the insect PPK gene family"

**Table S1. Source of peptide datasets used to identify PPK gene candidates in the insect genomes.**

| Insect species | Source | File |
|---|---|---|
| *Acyrthosiphon pisum* | bipaa.genouest.org/is/aphidbase/ | aphidbase_2.1b_pep.fasta |
| *Aedes aegypti* | Matthews et al. 2019 | Aedes-aegypti-LVP_AGWG_PEPTIDES_AaegL5.2.fa.gz |
| *Aedes albopictus* | ncbi.nlm.nih.gov/genome/?term=txid7160 | GCF_006496715.1_Aalbo_primary.1_protein.faa |
| *Anopheles gambiae* | Matthews et al. 2019 | Anopheles-gambiae-PEST_PEPTIDES_AgamP4.12.fa.gz |
| *Anoplophora glabripennis* | metazoa.ensembl.org/Anoplophora_glabripennis/Info/Index | Anoplophora_glabripennis.Agla_1.0.pep.all.fa.gz |
| *Apis mellifera* | hymenopteragenome.org/?q=hymenopteramine_datasets | Amel_OGSv3.2_consortium_pep.fa.gz |
| *Atta cephalotes* | hymenopteragenome.org/atta/?q=genome_consortium_datasets | acephalotes_OGS1.1_protein.fa.gz |
| *Blattella germanica* | ncbi.nlm.nih.gov/genome/?term=txid6973[Organism:noexp] | GCA_003018175.1_Bger_1.1_protein.faa.gz |
| *Bombus impatiens* | hymenopteragenome.org/?q=hymenopteramine_datasets | BIMP_2.0 (GCF_000188095.1) Protein - NCBI RefSeq Annotation Release 101 |
| *Bombyx mori* | lepbase.org/v4/sequence/ | Bombyx_mori_ASM15162v1_-_proteins.fa |
| *Camponotus floridanus* | hymenopteragenome.org/?q=hymenopteramine_datasets | Cflo_v1.0_refseq_pep.fa.gz |
| *Cimex lectularius* | www.vectorbase.org | Cimex-lectularius-Harlan_PEPTIDES_ClecH1.3.fa.gz |
| *Culex quinquefasciatus* | www.vectorbase.org | Culex-quinquefasciatus-Johannesburg_PEPTIDES_CpipJ2.4.fa.gz |
| *Danaus plexippus* | lepbase.org/v4/sequence/ | Danaus_plexippus_v3_-_proteins.fa |
| *Dendroctonus ponderosae* | metazoa.ensembl.org/Dendroctonus_ponderosae/Info/Index | Dendroctonus_ponderosae.DendPond_male_1.0.pep.all.fa.gz |
| *Drosophila melanogaster* | Liu et al. 2003 | - |
| *Glossina morsitans* | www.vectorbase.org | Culex-quinquefasciatus-Johannesburg_PEPTIDES_CpipJ2.4.fa.gz |
| *Leptinotarsa decemlineata* | data.nal.usda.gov/dataset/leptinotarsa-decemlineata-official-gene-set-v12 | Leptinotarsa decemlineataOGSv1.2 lepdec_OGSv1.2_GCF_000500325.1_pep.fa |
| *Locusta migratoria* | i5k.nal.usda.gov/locusta-migratoria | Locust.OGS_preQC.gff3.pep.gz |
| *Musca domestica* | ncbi.nlm.nih.gov/genome/?term=musca+domestica | GCF_000371365.1_Musca_domestica-2.0.2_protein.faa |
| *Myzus persicae* | bipaa.genouest.org/is/ myzus_persicae/ | Myzus_persicae_Clone_G006b_scaffolds.gff.pep.fa |
| *Pediculus humanus* | www.vectorbase.org | Pediculus-humanus-USDA_PEPTIDES_PhumU2.4.fa.gz |
| *Plutella xylostella* | lepbase.org/v4/sequence/ | Plutella_xylostella_pacbiov1_-_proteins.fa |
| *Rhodnius prolixus* | Latorre-Estivalis et al. 2017 | Rhodnius-prolixus-CDC_PEPTIDES_RproC3.3.fa.gz |
| *Spodoptera frugiperda* | bipaa.genouest.org/sp/spodoptera_frugiperda_pub/download/annotation/rice/OGS2.3_20151204/ | OGS2.3_20151204 |
| *Tribolium castaneum* | beetlebase.org | BeetleBase3.0 |

**Table S2. Analysis of PPK sequences.** Columns 9-15 display the presence of conserved residues and motifs described by (L. Liu, Leonard, et al., 2003) for the PPK sequences from *D. melanogaster*. SF: Subfamily, Aa: amino acids; TM: transmembrane domains; Cs: cysteines; DEG: degenerin residue; “x” indicates any residue; “h” indicates a hydrophobic residue; Abs: Absent. Length is expressed in number of amino acids.

| Database code | Annotation | SF | Location of PFAM0056 | | E-Value of BLAST search against PFAM database | Length | Nº TM | Nº Cs | GxS | W | T/SxhHG | FPhhThC | DEG | GxCxxFN | N-Terminus length | Extracel. Loop length | C-Terminus length |
|---|---|---|---|---|---|---|---|---|---|---|---|---|---|---|---|---|---|
| | | | From | To | | | | | | | | | | | | | |
| AAEL014009-PA | *AaegPPK101* | VI | 51 | 473 | 2.41E-79 | 618 | 2 | 14 | GFS | W | STLHG | FPTVMVC | V | GFCYSFN | 73 | 354 | 140 |
| AAEL014010-PB | *AaegPPK102* | VI | 25 | 431 | 8.84E-67 | 576 | 2 | 14 | GFS | W | STLHG | FPTVLVC | V | GFCYGFN | 43 | 342 | 140 |
| AAEL002575-PB | *AaegPPK103* | IV | 66 | 541 | 3.33E-84 | 575 | 2 | 14 | GLS | W | ITTHC | FPAISLC | S | GFCCTFN | 86 | 409 | 29 |
| AAEL022385-PA | *AaegPPK201* | II | 16 | 416 | 1.21E-39 | 438 | 2 | 12 | GAS | W | SSLHG | FPSLTVC | V | GICYSFN | 36 | 337 | 14 |
| AAEL019946-PB | *AaegPPK202* | II | 4 | 206 | 2.98E-24 | 223 | 2 | 10 | GKD | - | Abs. | Abs. | V | Abs. | 1 | 187 | 9 |
| AAEL003714-PB | *AaegPPK203* | IV | 25 | 355 | 1.34E-54 | 454 | 1 | 13 | GGS | - | Abs. | Abs. | A | GFCCSFN | - | 334 | 94 |
| AAEL010779-PA | *AaegPPK204* | IV | 56 | 518 | 1.39E-90 | 548 | 2 | 14 | GFS | W | TALHG | FPAVTIC | A | GLCCSFN | 77 | 395 | 25 |
| AAEL022016-PA | *AaegPPK205* | II | 36 | 386 | 2.28E-31 | 399 | 4 | 7 | GFS | W | GALHG | LPSLTIC | V | GICYTTN | 58 | 282 | 8 |
| AAEL006613-PA | *AaegPPK206* | I | 10 | 432 | 3.19E-65 | 453 | 2 | 14 | GAS | W | STIHG | FPAIVVC | V | GLCYAVN | 32 | 354 | 16 |
| AAEL000582-PC | *AaegPPK301* | V | 40 | 502 | 1.10E-96 | 612 | 2 | 14 | GFS | F | TTIHG | FPAVTIC | S | GLCCTFN | 62 | 357 | 105 |
| AAEL006258-PA | *AaegPPK302* | I | 18 | 451 | 2.34E-64 | 481 | 2 | 14 | GAS | W | CSLVG | IPSLGIC | V | GSCYLLN | 40 | 394 | 25 |
| AAEL008809-PB | *AaegPPK303* | II | 23 | 341 | 1.25E-32 | 353 | 1 | 7 | Abs. | W | TSAHG | FPSLTVC | - | GLCLSVN | 44 | 365 | - |
| AAEL003470-PB | *AaegPPK305* | IV | 9 | 325 | 6.22E-42 | 373 | 1 | 13 | GMS | - | Abs. | Abs. | A | GACCSFN | - | 283 | 43 |
| AAEL014228-PA | *AaegPPK306* | V | 59 | 536 | 4.39E-95 | 571 | 2 | 14 | GAS | W | TPIHG | FPAVTIC | A | GFCYTFN | 82 | 304 | 30 |
| AAEL026145-PA | *AaegPPK307* | V | 1 | 129 | 5.51E-12 | 178 | 1 | 5 | GSS | - | Abs. | Abs. | A | Abs. | - | 408 | 44 |
| AAEL014230-PA | *AaegPPK308* | V | 33 | 515 | 1.62E-110 | 557 | 2 | 14 | GVS | W | TSIHG | FPAVTIC | A | GFCYTFN | 55 | 108 | 37 |
| AAEL011005-PA | *AaegPPK309* | V | 12 | 491 | 1.11E-77 | 515 | 2 | 14 | GAS | W | SSVNA | FPAVTIC | A | GLCYTFN | 34 | 414 | 19 |
| AAEL011002-PB | *AaegPPK310* | V | 22 | 121 | 7.03E-14 | 255 | 1 | 10 | GAS | - | Abs. | Abs. | A | Abs. | 1 | 411 | 39 |
| AAEL026983-PA | *AaegPPK311* | V | 17 | 493 | 2.42E-85 | 516 | 2 | 14 | GAS | W | CSIYA | FPAVTVC | A | GVCYTFN | 38 | 189 | 18 |
| AAEL010995-PB | *AaegPPK312* | V | 20 | 511 | 3.93E-77 | 543 | 2 | 14 | GAS | W | NSIRA | FPAITIC | A | GICYTFN | 37 | 409 | 27 |
| AAEL019676-PA | *AaegPPK313* | V | 20 | 556 | 3.21E-74 | 599 | 2 | 14 | GTS | W | SSVHG | FPSFTIC | A | GLCVNFN | 42 | 428 | 38 |
| AAEL019675-PA | *AaegPPK314* | V | 6 | 428 | 1.94E-45 | 471 | 1 | 13 | GTS | - | Abs. | Abs. | A | GLCVNFN | - | 468 | 38 |
| AAEL000863-PA | *AaegPPK315* | V | 21 | 433 | 7.15E-50 | 471 | 2 | 11 | GAS | W | TSVHG | FPGMTIC | A | STCHTIN | 45 | 407 | 33 |
| AAEL000926-PB | *AaegPPK316* | V | 20 | 510 | 1.49E-100 | 550 | 2 | 14 | GVS | W | SSVHG | FPAVTIC | A | GICVSFN | 42 | 342 | 35 |
| AAEL000873-PB | *AaegPPK317* | V | 20 | 509 | 2.26E-104 | 544 | 2 | 14 | GMS | W | TSIHG | FPAVTIC | A | GVCLTFN | 42 | 422 | 30 |
| AAEL008053-PA | *AaegPPK318* | V | 22 | 504 | 5.46E-85 | 550 | 2 | 13 | GVS | W | SSIHG | FPAVTIC | A | GVCYTFN | 44 | 421 | 41 |
| AAEL000534-PB | *AaegPPK319* | V | 2 | 368 | 3.12E-50 | 398 | 1 | 13 | GVS | - | Abs. | Abs. | S | GICFTIN | 1 | 414 | 25 |

| | | | | | | | | | | | | | | | | | |
|---|---|---|---|---|---|---|---|---|---|---|---|---|---|---|---|---|---|
| AAEL000552-PB | *AaegPPK320* | V | 64 | 430 | 5.29E-63 | 460 | 1 | 13 | GVS | - | Abs. | Abs. | A | GICFTFN | 19 | 346 | 25 |
| AAEL000547-PA | *AaegPPK321* | V | 50 | 540 | 2.45E-99 | 569 | 2 | 14 | GVS | W | SSVHG | FPAITIC | A | GICYTYN | 71 | 388 | 24 |
| AAEL004091-PB | *AaegPPK322* | V | 39 | 523 | 4.16E-114 | 561 | 2 | 14 | GVS | W | STIHG | FPAVTIC | A | GICYTFN | 61 | 423 | 33 |
| AAEL027684-PA | *AaegPPK323* | IV | 9 | 250 | 2.95E-20 | 298 | 0 | 10 | GMS | - | Abs. | Abs. | S | GACCSFN | - | 416 | 43 |
| AAEL023544-PA | *AaegPPK29* | I | 1 | 389 | 1.07E-13 | 396 | 1 | 11 | GVS | L | Abs. | FPATSIC | V | GLCFISN | 1 | 229 | 2 |
| XP_019525011.2 | *Aalb019525011.2* | V | 34 | 524 | 1.4524E-101 | 568 | 2 | 14 | GVS | W | SSVHG | FPAVTI | A | GICVSFN | 56 | 422 | 39 |
| XP_019536370.2 | *Aalb019536370.2* | V | 39 | 523 | 1.5226E-113 | 561 | 3 | 14 | GVS | W | STIHG | FPAVTI | A | GICYTFN | 61 | 416 | 33 |
| XP_019537219.2 | *Aalb019537219.2* | V | 22 | 504 | 9.46131E-83 | 550 | 2 | 13 | GVS | W | SSIHG | FPAVTI | A | GLCYTFN | 44 | 414 | 41 |
| XP_019539770.2 | *Aalb019539770.2* | V | 51 | 542 | 4.17856E-95 | 573 | 2 | 14 | GVS | W | SSIHG | FPAITI | A | GICYTYN | 73 | 423 | 26 |
| XP_019556935.2 | *Aalb019556935.2* | V | 16 | 492 | 2.20104E-83 | 516 | 2 | 14 | GAS | W | GSIYA | FPAVTV | A | GVCYTFN | 37 | 409 | 19 |
| XP_019556948.2 | *Aalb019556948.2* | V | 16 | 492 | 2.32689E-84 | 515 | 4 | 14 | GAS | W | CSIYA | FPAVTV | A | GVCYTFN | 37 | 409 | 18 |
| XP_019557794.2 | *Aalb019557794.2* | II | 16 | 415 | 2.02337E-42 | 436 | 3 | 12 | GAS | W | SSLHG | FPSLTV | V | GICYSFN | 36 | 336 | 13 |
| XP_029712101.1 | *Aalb029712101.1* | V | 43 | 576 | 4.57599E-60 | 617 | 2 | 12 | GTS | W | SSVHG | FPSFTI | A | GFCVNFN | 65 | 465 | 36 |
| XP_029713859.1 | *Aalb029713859.1* | II | 16 | 415 | 8.5535E-44 | 436 | 3 | 12 | GAS | W | SSLHG | FPSLTV | V | GICYSFN | 36 | 336 | 13 |
| XP_029720868.1 | *Aalb029720868.1* | V | 41 | 525 | 2.91014E-97 | 561 | 2 | 14 | GVS | W | TPIHG | FPAVTL | A | GFCYTFN | 64 | 415 | 31 |
| XP_029720869.1 | *Aalb029720869.1* | V | 33 | 515 | 3.3425E-111 | 558 | 2 | 14 | GVS | W | TSIHG | FPAVTI | A | GFCYTFN | 55 | 414 | 38 |
| XP_029721586.1 | *Aalb029721586.1* | V | 28 | 507 | 6.50184E-69 | 518 | 2 | 14 | GSS | W | NFVYA | FPAITV | A | GICYTFN | 36 | 425 | 6 |
| XP_029721587.1 | *Aalb029721587.1* | V | 45 | 197 | 1.4856E-16 | 285 | 1 | 4 | Abs. | W | ATKDA | FPAITI | - | GICFTFN | 38 | 221 | - |
| XP_029721588.1 | *Aalb029721588.1* | V | 1 | 129 | 3.25403E-10 | 178 | 1 | 5 | GSS | - | Abs. | Abs. | A | Abs. | - | 108 | 44 |
| XP_029721589.1 | *Aalb029721589.1* | V | 12 | 491 | 1.67567E-74 | 515 | 2 | 14 | GAS | W | SSINA | FPAVTI | S | GVCYTFN | 34 | 411 | 19 |
| XP_029721590.1 | *Aalb029721590.1* | V | 1 | 360 | 1.28171E-44 | 387 | 0 | 14 | Abs. | - | Abs. | FPAVTV | - | GICYTFN | 1 | 381 | - |
| XP_029721592.1 | *Aalb029721592.1* | V | 112 | 341 | 4.40405E-47 | 364 | 1 | 10 | GAS | - | Abs. | Abs. | A | Abs. | 13 | 300 | 18 |
| XP_029731012.1 | *Aalb029731012.1* | V | 20 | 509 | 8.6209E-107 | 546 | 2 | 14 | GMS | W | TSIHG | FPAVTI | A | GVCLTFN | 42 | 421 | 32 |
| XP_029731018.1 | *Aalb029731018.1* | V | 21 | 433 | 7.48268E-52 | 476 | 2 | 9 | GAS | W | TSVHG | FPGMTI | A | RSSWDLD | 45 | 342 | 38 |
| XP_029731325.1 | *Aalb029731325.1* | V | 22 | 504 | 5.27614E-84 | 550 | 2 | 13 | GVS | W | SSIHG | FPAVTI | A | GLCYTFN | 44 | 414 | 41 |
| XP_029731661.1 | *Aalb029731661.1* | V | 45 | 434 | 1.18536E-61 | 464 | 1 | 13 | GVS | - | Abs. | SRHRIL | A | GICFTFN | 19 | 392 | 25 |
| XP_029731663.1 | *Aalb029731663.1* | V | 6 | 367 | 4.5214E-66 | 397 | 1 | 13 | GVS | - | Abs. | Abs. | S | GICFTFN | 1 | 345 | 25 |
| XP_029731672.1 | *Aalb029731672.1* | V | 52 | 434 | 7.83978E-62 | 464 | 1 | 13 | GVS | - | Abs. | SRHRIL | A | GICFTFN | 19 | 392 | 25 |
| XP_029732095.1 | *Aalb029732095.1* | V | 54 | 547 | 4.5275E-102 | 577 | 2 | 14 | GVS | W | STIHG | FPAVTI | S | GICFTFN | 76 | 425 | 25 |
| XP_029732096.1 | *Aalb029732096.1* | V | 58 | 431 | 3.44378E-63 | 461 | 1 | 13 | GVS | - | Abs. | Abs. | A | GICFTFN | 14 | 394 | 25 |
| XP_029732188.1 | *Aalb029732188.1* | V | 48 | 440 | 9.68534E-43 | 448 | 2 | 11 | GVS | W | TSFLP | FPAITI | A | GICFTFN | 57 | 337 | 3 |
| XP_029734091.1 | *Aalb029734091.1* | V | 149 | 522 | 3.36062E-66 | 552 | 1 | 13 | GVS | C | GVLVT | WGCRIL | A | GICFTFN | 105 | 394 | 25 |
| XP_029734092.1 | *Aalb029734092.1* | V | 2 | 429 | 1.06046E-75 | 459 | 1 | 14 | GVS | - | Abs. | FPAVTI | S | GICFTFN | 1 | 407 | 25 |
| XP_029734094.1 | *Aalb029734094.1* | V | 53 | 435 | 5.03517E-61 | 465 | 1 | 13 | GVS | - | Abs. | SHHRIL | A | GICFTFN | 19 | 393 | 25 |
| XP_029734497.1 | *Aalb029734497.1* | IV | 35 | 503 | 1.37909E-76 | 585 | 2 | 14 | GGS | W | TSLHG | FPAVTV | A | GFCCSFN | 63 | 394 | 77 |
| XP_029735277.1 | *Aalb029735277.1* | IV | 25 | 355 | 1.40085E-54 | 437 | 1 | 13 | GGS | - | Abs. | Abs. | A | GFCCSFN | - | 334 | 77 |

| | | | | | | | | | | | | | | | | | |
|---|---|---|---|---|---|---|---|---|---|---|---|---|---|---|---|---|---|
| XP_029715164.1 | *AalbPPK10* | II | 27 | 360 | 1.66181E-40 | 373 | 4 | 11 | GFS | W | GALHG | LPSLTI | V | MFEFIES | 49 | 265 | 8 |
| XP_029720145.1 | *AalbPPK13a* | I | 10 | 432 | 6.29455E-65 | 453 | 2 | 14 | GAS | W | STIHG | FPAIVV | V | GLCYAVN | 32 | 354 | 16 |
| XP_029729201.1 | *AalbPPK13b* | I | 10 | 432 | 1.27009E-64 | 453 | 2 | 14 | GAS | W | STIHG | FPAIVV | V | GLCYAVN | 32 | 354 | 16 |
| XP_019558124.2 | *AalbPPK15a* | I | 18 | 451 | 4.20818E-65 | 482 | 2 | 14 | GAS | W | CSLVG | FPSLGI | V | GSCYLLN | 40 | 365 | 26 |
| XP_029726981.1 | *AalbPPK15b* | I | 18 | 468 | 2.02193E-62 | 499 | 2 | 14 | GAS | W | CSLVG | FPSLGI | V | GSCYLLN | 40 | 382 | 26 |
| XP_019555869.2 | *AalbPPK17a* | VII | 15 | 367 | 5.70291E-07 | 443 | 2 | 8 | GLS | R | LCYD | YPAITF | A | GFSILLQ | 15 | 312 | 51 |
| XP_029729547.1 | *AalbPPK17b* | VII | 15 | 367 | 3.7138E-08 | 443 | 2 | 8 | GLS | R | LCYD | YPAITF | A | GFSILLQ | 15 | 312 | 51 |
| XP_029711542.1 | *AalbPPK23a* | VI | 39 | 446 | 6.65572E-69 | 587 | 2 | 13 | GFS | W | STLHG | FPTVLV | V | GFCYGFN | 57 | 343 | 136 |
| XP_019530003.2 | *AalbPPK23b* | VI | 53 | 475 | 6.16714E-79 | 623 | 2 | 13 | GFS | W | STLHG | FPTVMV | V | GFCYSFN | 75 | 354 | 143 |
| XP_019540188.2 | *AalbPPK25* | II | 8 | 320 | 5.0714E-17 | 350 | 2 | 7 | GCS | W | SSVHG | RPAATV | V | GVCFSSS | 29 | 256 | 14 |
| XP_029709904.1 | *AalbPPK29a* | I | 1 | 397 | 3.60667E-18 | 404 | 1 | 11 | GVS | - | Abs. | FPATSI | V | GFCFISN | 1 | 357 | 2 |
| XP_029733934.1 | *AalbPPK29b* | I | 1 | 397 | 3.27778E-18 | 404 | 1 | 11 | GVS | - | Abs. | FPATSI | V | GFCFISN | 1 | 357 | 2 |
| XP_029733926.1 | *AalbPPK29c* | I | 29 | 376 | 3.07208E-14 | 383 | 1 | 10 | GVS | - | Abs. | ATTHTI | V | GFCFISN | 1 | 353 | 2 |
| XP_019551763.2 | *AalbPPK301A* | V | 40 | 502 | 6.13034E-96 | 613 | 2 | 14 | GFS | F | TTLHG | FPAVTI | S | GLCCTFN | 62 | 394 | 106 |
| XP_029707777.1 | *AalbPPK301b* | V | 40 | 502 | 2.21739E-95 | 613 | 2 | 14 | GFS | F | TTLHG | FPAVTI | S | GLCCTFN | 62 | 394 | 106 |
| XP_029715424.1 | *AalbPPK31* | I | 39 | 315 | 5.91039E-30 | 332 | 2 | 13 | GIS | - | Abs. | Abs. | I | GTCYSIN | 1 | 298 | 7 |
| XP_029730550.1 | *AalbPPK3* | II | 86 | 364 | 3.02287E-26 | 384 | 2 | 11 | GKD | R | Abs. | LSAPLL | V | IECTYAK | 1 | 343 | 12 |
| XP_029735319.1 | *AalbPPK9* | II | 23 | 331 | 1.65941E-31 | 378 | 1 | 7 | VIF | W | TSAHG | FPSLTV | H | GLCLSVN | 44 | 308 | - |
| ACEP16277-PA | *Acep16277* | IV | 1 | 325 | 4.17703E-32 | 340 | 0 | 14 | Abs. | - | Abs. | FPALTIC | A | GICCSFN | 2 | 327 | 3 |
| ACEP17448-PA | *Acep17448* | IV | 3 | 423 | 2.36293E-74 | 466 | 2 | 12 | GGS | W | Abs. | FPAVTIC | A | GFCCAFN | 9 | 368 | 38 |
| ACEP19650-PA | *Acep19650* | IV | 109 | 494 | 7.41077E-30 | 513 | 2 | 4 | GFS | W | SSINS | FPGIAIC | S | GYCCTFN | 106 | 343 | 12 |
| ACEP21061-PA | *Acep21061* | IV | 40 | 412 | 5.04595E-32 | 415 | 0 | 11 | Abs. | - | SSFNR | Abs. | - | GYCCTFN | 31 | 383 | - |
| ACEP19465-PA | *AcepPPK16* | IV | 3 | 291 | 1.69912E-42 | 292 | 0 | 9 | Abs. | - | Abs. | FPAITVC | - | GLCCSFN | 1 | 285 | - |
| ACEP22748-PA | *AcepPPK28* | V | 5 | 406 | 6.21888E-68 | 458 | 1 | 14 | GLQ | - | Abs. | FPSVTIC | C | GICCNFN | 1 | 403 | 26 |
| AGAP001631-PA | *Agam001631* | IV | 50 | 519 | 8.28822E-80 | 549 | 2 | 14 | GMS | W | ITAHC | FPAISLC | S | GFCCTFN | 71 | 402 | 25 |
| AGAP006703-PA | *Agam006703* | IV | 25 | 441 | 8.94196E-65 | 529 | 2 | 9 | GGS | W | SSLHG | FPAVTLC | A | GYCCSFN | 50 | 349 | 79 |
| AGAP006704-PA | *Agam006704* | IV | 21 | 477 | 4.36962E-82 | 565 | 2 | 12 | GGS | W | TSLHG | FPAVTVC | A | GFCCSYN | 51 | 380 | 83 |
| AGAP007084-PA | *Agam007084* | II | 18 | 435 | 1.65456E-36 | 481 | 2 | 12 | GFS | W | TSLHG | FPSLTVC | V | GICYTFN | 36 | 353 | 41 |
| AGAP011103-PA | *Agam011103* | V | 16 | 520 | 4.74027E-78 | 562 | 2 | 14 | GMS | W | STCHG | FPAITIC | A | GVCYSFN | 38 | 436 | 37 |
| AGAP011433-PA | *Agam011433* | V | 38 | 462 | 1.94695E-36 | 471 | 2 | 11 | GVS | W | STFPI | FPAVTLC | G | GICYTFN | 60 | 356 | 4 |
| AGAP011610-PA | *Agam011610* | V | 42 | 525 | 3.8137E-108 | 575 | 2 | 14 | GVS | W | STIHG | FPAVTIC | A | GICYTFN | 64 | 415 | 45 |
| AGAP011611-PA | *Agam011611* | V | 44 | 524 | 7.71521E-94 | 568 | 2 | 14 | GVS | W | SSIHG | FPAVTIC | A | GICYTFN | 66 | 412 | 39 |
| AGAP012279-PA | *Agam012279* | V | 9 | 493 | 1.0125E-107 | 511 | 2 | 14 | GVS | W | STVHG | FPAVTIC | A | GICFTFN | 52 | 395 | 13 |
| AGAP028699-PB | *Agam028699* | V | 10 | 467 | 3.86022E-93 | 502 | 2 | 14 | GVS | W | Abs. | FPAITIC | A | GFCYTFN | 9 | 412 | 30 |

| | | | | | | | | | | | | | | | | | |
|---|---|---|---|---|---|---|---|---|---|---|---|---|---|---|---|---|---|
| AGAP028700-PA | *Agam028700* | V | 20 | 499 | 2.1997E-100 | 507 | 2 | 14 | GVS | W | SSIHG | FPAITIC | A | GFCYTFN | 42 | 411 | 3 |
| AGAP028701-PA | *Agam028701* | V | 10 | 470 | 1.90552E-90 | 505 | 2 | 14 | GVS | W | Abs. | FPAVTIC | A | GICYTFN | 9 | 415 | 30 |
| AGAP028702-PA | *Agam028702* | V | 10 | 470 | 5.5964E-94 | 510 | 2 | 14 | GVS | W | Abs. | FPAITIC | A | GICYTFN | 9 | 415 | 35 |
| AGAP028703-PA | *Agam028703* | V | 15 | 399 | 3.8174E-53 | 424 | 1 | 14 | Abs. | W | Abs. | FPAITIC | - | GICYSFN | 14 | 367 | 17 |
| AGAP009789-PA | *AgamPPK10* | II | 47 | 496 | 5.0189E-58 | 509 | 2 | 12 | GFS | W | GAVHG | MPSLTIC | V | GICYTTN | 69 | 381 | 8 |
| AGAP007945-PA | *AgamPPK13* | I | 10 | 432 | 1.26534E-65 | 455 | 2 | 14 | GAS | W | STFHG | FPSIVVC | V | GLCYAVN | 32 | 354 | 18 |
| AGAP008378-PA | *AgamPPK15a* | I | 17 | 451 | 2.99493E-61 | 489 | 2 | 14 | GAS | W | CSLAG | FPSVGVC | V | GSCFLLN | 39 | 366 | 33 |
| AGAP008380-PA | *AgamPPK15b* | I | 18 | 460 | 2.56002E-55 | 503 | 3 | 14 | GAS | W | ASLAG | FPAVGVC | V | GTCYLLN | 40 | 374 | 38 |
| AGAP009590-PA | *AgamPPK16* | IV | 45 | 508 | 6.13354E-88 | 533 | 2 | 14 | GFS | W | TALHG | FPAVTVC | A | GLCCSFN | 66 | 396 | 20 |
| AGAP010146-PA | *Agamppk17* | VII | 18 | 361 | 2.13164E-08 | 443 | 2 | 10 | GLS | F | Abs. | YPAITFC | S | GRCYTLN | 15 | 312 | 51 |
| AGAP000840-PA | *AgamPPK23* | VI | 64 | 502 | 1.72124E-80 | 656 | 2 | 14 | GFS | W | STLHG | FPTVLVC | V | GFCYAFN | 86 | 370 | 149 |
| AGAP005516-PA | *AgamPPK25* | II | 7 | 409 | 1.33456E-44 | 431 | 2 | 12 | GCS | W | SSIHG | PPAATVC | V | GVCYSST | 28 | 335 | 17 |
| AGAP006720-PA | *AgamPPK3* | II | 6 | 400 | 1.10546E-27 | 445 | 2 | 12 | GKN | - | Abs. | MPGLTFC | V | GICYALN | 1 | 373 | 9 |
| AGAP001602-PA | *AgamPPK301* | V | 39 | 485 | 1.0722E-76 | 575 | 2 | 14 | Abs. | F | TTIHG | FPAVTIC | C | GLCCTFN | 61 | 400 | 28 |
| AGAP000657-PA | *AgamPPK31* | I | 2 | 396 | 1.09292E-38 | 417 | 1 | 14 | GWS | - | Abs. | FPTVTLC | I | GRCFSVN | 1 | 360 | 69 |
| AGAP010430-PA | *AgamPPK6* | IV | 76 | 525 | 2.58635E-70 | 557 | 2 | 14 | GIS | W | SKIHG | FPGITLC | S | GACCSFN | 98 | 381 | 27 |
| AGAP004474-PA | *AgamPPK9* | II | 22 | 452 | 1.33856E-39 | 459 | 2 | 12 | GCS | W | TSAHG | FPSLTIC | V | GLCLAVN | 43 | 363 | 2 |
| AGLA006011 | *AglaPPK27* | IV | 15 | 376 | 1.0363E-46 | 377 | 1 | 11 | Abs. | W | TTIHA | HPGVAIC | - | GVCCVFN | 37 | 314 | - |
| AGLA006813 | *Agla006813* | IV | 118 | 398 | 8.6337E-39 | 537 | 2 | 10 | Abs. | W | Abs. | FPGVTIC | A | GICCSFN | 117 | 257 | 112 |
| AGLA012530 | *Agla012530* | V | 21 | 194 | 6.8456E-27 | 216 | 0 | 8 | Abs. | - | Abs. | Abs. | - | Abs. | 6 | 206 | 2 |
| AGLA012532 | *Agla012532* | V | 73 | 462 | 2.35509E-60 | 482 | 2 | 12 | GFS | - | Abs. | FPAVTIC | A | ICPETKS | 71 | 345 | 15 |
| AGLA012536 | *Agla012536* | V | 52 | 524 | 6.51518E-94 | 544 | 2 | 14 | GFS | W | TGIHG | FPAVTIC | S | GNCYSFN | 73 | 405 | 15 |
| AGLA013191 | *Agla013191* | V | 43 | 483 | 2.24709E-93 | 514 | 2 | 14 | GFS | W | TGIHG | FPAVTIC | A | GLCFTFN | 65 | 372 | 26 |
| AGLA013193 | *Agla013193* | V | 39 | 275 | 5.84695E-48 | 305 | 1 | 10 | GFS | - | LH | Abs. | A | Abs. | 1 | 253 | 25 |
| AGLA013194 | *Agla013194* | V | 32 | 231 | 1.95974E-20 | 367 | 1 | 5 | GEE | - | Abs. | Abs. | A | Abs. | 3 | 264 | 68 |
| AGLA014384 | *Agla014384* | V | 153 | 478 | 1.98889E-53 | 487 | 1 | 9 | Abs. | W | SGIHG | FPAVTVC | E | Abs. | 174 | 287 | - |
| AGLA014385 | *Agla014385* | V | 39 | 396 | 1.3831E-40 | 593 | 2 | 12 | GNQ | W | AGIHG | FPAVTIC | E | Abs. | 49 | 480 | 14 |
| AGLA014386 | *Agla014386* | V | 17 | 80 | 0.00361051 | 394 | 2 | 11 | GFS | - | VKVHG | PNVQRVC | A | Abs. | 31 | 317 | 17 |
| AGLA014387 | *Agla014387* | V | 30 | 196 | 1.99838E-30 | 1182 | 4 | 11 | GFS | - | TGIHG | FPAVTIC | A | GLCFTFN | 749 | 380 | 27 |
| AGLA019103 | *AglaPPK28* | V | 269 | 411 | 3.23224E-14 | 446 | 1 | 5 | GFS | W | Abs. | GAT | T | GVLG | 1 | 352 | 30 |
| AGLA003479 | *AglaPPK13* | I | 1 | 383 | 2.74077E-51 | 418 | 2 | 14 | GAS | - | Abs. | FPSIAIC | V | GRCYGIN | 56 | 285 | - |
| AGLA005467 | *AglaPPK16* | IV | 34 | 361 | 4.68994E-57 | 367 | 1 | 9 | Abs. | W | TALHG | FPAVTVC | - | GFCCSFN | 38 | 329 | 30 |
| AGLA017497 | *AglaPPK3* | II | 8 | 301 | 1.04793E-15 | 320 | 3 | 6 | GIS | W | Abs. | FPAATGC | V | ETAYSDN | 8 | 247 | 14 |
| AGLA016101 | *AglaPPK9* | II | 16 | 352 | 5.9617E-37 | 368 | 3 | 6 | GCS | W | SSVHC | FPSVTLC | V | GICYSFN | 38 | 268 | 11 |

| | | | | | | | | | | | | | | | | | |
|---|---|---|---|---|---|---|---|---|---|---|---|---|---|---|---|---|---|
| GB48330-PA | *Amel48330* | IV | 35 | 500 | 1.94181E-78 | 571 | 2 | 13 | GGS | W | TALHG | FPGITIC | A | GFCCGFN | 56 | 402 | 62 |
| GB48363-PA | *Amel48363* | IV | 67 | 408 | 6.92484E-32 | 437 | 1 | 11 | Abs. | W | TSVHG | FPAVTIC | C | Abs. | 91 | 317 | 3 |
| GB53731-PA | *Amel53731* | I | 14 | 409 | 7.92058E-43 | 421 | 2 | 12 | GAS | W | CSIHG | FPAIAFC | V | GYCLAIN | 51 | 383 | 51 |
| GB55337-PA | *Amel55337* | IV | 29 | 480 | 4.37953E-71 | 536 | 2 | 12 | GFS | W | TKLAG | FPGIAIC | G | GFCCTFN | 87 | 345 | 19 |
| GB53792-PA | *Amel53792* | II | 65 | 478 | 4.56303E-33 | 502 | 2 | 12 | GAS | - | GTLHG | FKIPRWS LFVC | T | GLCLVSR | 36 | 330 | 4 |
| GB45440-PA | *AmelPPK16* | IV | 37 | 509 | 2.00283E-80 | 528 | 2 | 14 | GFS | W | TGLHG | FPAVTIC | A | GICCSFN | 59 | 404 | 14 |
| GB53179-PA | *AmelPPK23* | VI | 23 | 393 | 3.93322E-39 | 446 | 2 | 14 | GFS | W | NSLHG | FPQIFIC | V | Abs. | 45 | 302 | 48 |
| GB46186-PA | *AmelPPK28* | V | 129 | 535 | 4.14598E-72 | 550 | 1 | 14 | Abs. | W | STLHG | FPSVTIC | - | GLCCNFN | 151 | 373 | - |
| ACYPI005555-PA | *Apis005555* | I | 7 | 358 | 2.29168E-40 | 368 | 1 | 14 | Abs. | W | STIHG | FLSLSIC | - | GVCFIIN | 29 | 308 | 5 |
| ACYPI008626-PA | *Apis008626* | I | 5 | 422 | 7.07765E-62 | 448 | 2 | 14 | GAS | W | STIHG | FPAVSVC | V | GICYTLT | 27 | 349 | 21 |
| ACYPI068872-PA | *Apis068872* | V | 6 | 334 | 7.95796E-40 | 356 | 1 | 12 | GCS | - | Abs. | Abs. | G | GLCYSIN | 1 | 312 | 17 |
| ACYPI27784-PA | *Apis27784* | III | 10 | 478 | 9.52277E-79 | 509 | 2 | 14 | GCS | W | GSIHG | FPAVTIC | V | GFCYSFN | 32 | 401 | 25 |
| ACYPI28482-PA | *Apis28482* | I | 46 | 273 | 5.24736E-08 | 340 | 2 | 10 | GIN | W | Abs. | VPAVTVC | - | TPCFCIN | 12 | 285 | 5 |
| ACYPI29894-PA | *Apis29894* | V | 45 | 520 | 9.57637E-55 | 578 | 2 | 13 | GAS | W | TSLHG | FPGITVC | G | GVCYSLN | 67 | 407 | 53 |
| ACYPI30092-PA | *Apis30092* | V | 17 | 321 | 1.75216E-16 | 330 | 1 | 11 | Abs. | W | Abs. | FPAVTIC | - | DVLQTGK | 27 | 277 | - |
| ACYPI33127-PA | *Apis33127* | V | 98 | 322 | 9.97047E-15 | 439 | 1 | 10 | SIS | - | Abs. | Abs. | - | GLCFTVN | 1 | 399 | 29 |
| ACYPI33128-PA | *Apis33128* | V | 16 | 352 | 2.64123E-14 | 374 | 0 | 9 | Abs. | - | Abs. | FPSIMIC | - | GVCYTIN | - | 357 | - |
| ACYPI33129-PA | *Apis33129* | V | 16 | 342 | 1.28862E-18 | 375 | 1 | 7 | Abs. | W | KSINL | FPMITIC | - | GLCFAIN | 27 | 322 | - |
| ACYPI33363-PA | *Apis33363* | III | 25 | 494 | 4.65335E-59 | 509 | 2 | 14 | GFS | W | EVLYS | FPSITIC | V | GFCYSFN | 29 | 420 | 9 |
| ACYPI33364-PA | *Apis33364* | III | 62 | 537 | 1.68839E-73 | 561 | 2 | 14 | GFS | W | CTVHG | FPAVNVC | V | GFCYSFN | 84 | 408 | 18 |
| ACYPI34461-PA | *Apis34461* | III | 23 | 215 | 1.25888E-16 | 218 | 1 | 4 | Abs. | W | ISVNA | FPSVTIC | K | Abs. | 39 | 147 | - |
| ACYPI34462-PA | *Apis34462* | III | 14 | 258 | 1.32704E-23 | 282 | 2 | 10 | GAS | - | Abs. | Abs. | V | Abs. | 1 | 231 | 19 |
| ACYPI34467-PA | *Apis34467* | III | 14 | 463 | 7.6513E-49 | 1086 | 4 | 14 | GGS | W | Abs. | FPMVVIC | V | GLCHSFN | 581 | 402 | 52 |
| ACYPI35976-PA | *Apis35976* | V | 64 | 558 | 4.12746E-72 | 588 | 2 | 13 | GFS | W | TSLHG | FPAVTVC | G | GLCYSFN | 86 | 426 | 25 |
| ACYPI44656-PA | *Apis44656* | III | 43 | 182 | 1.85734E-05 | 242 | 0 | 8 | Abs. | - | Abs. | Abs. | - | Abs. | 1 | 188 | 26 |
| ACYPI008502-PA | *ApisPPK23* | VI | 29 | 449 | 1.57038E-75 | 515 | 3 | 14 | GFS | W | STLHG | FPAITVC | V | GFCYSFN | 51 | 352 | 61 |
| ACYPI50440-PA | *ApisPPK28* | V | 59 | 470 | 2.57545E-71 | 486 | 1 | 14 | Abs. | W | TTLHG | FPAITIC | - | GICCSFN | 81 | 379 | - |
| ACYPI31763-PA | *Apis31763* | II | 112 | 417 | 1.37573E-24 | 445 | 2 | 11 | Abs. | W | TPTHV | FPSSTFC | M | Abs. | 118 | 276 | 7 |
| ACYPI000481-PA | *ApisPPK9* | II | 44 | 481 | 2.52475E-55 | 494 | 2 | 12 | GCS | W | SNIHG | LPAVTLC | V | GLCYSYN | 66 | 369 | 8 |
| PSN30177.1 | *Bger30177* | III | 24 | 321 | 2.75715E-46 | 375 | 2 | 11 | GFS | - | Abs. | Abs. | V | GFCYSFN | 1 | 295 | 49 |
| PSN32397.1 | *Bger32397* | I | 16 | 232 | 1.59123E-11 | 238 | 0 | 9 | Abs. | - | Abs. | HLHC | S | GPCLAIN | 1 | 226 | - |
| PSN33731.1 | *Bger33731* | III | 6 | 273 | 1.35199E-21 | 280 | 0 | 11 | Abs. | - | Abs. | Abs. | - | GFCYTFN | 1 | 257 | - |
| PSN34456.1 | *Bger34456* | V | 28 | 385 | 1.23238E-56 | 393 | 1 | 9 | Abs. | W | TSLHG | FPAVTIC | - | GLCYTSN | 50 | 317 | - |

| | | | | | | | | | | | | | | | | | |
|---|---|---|---|---|---|---|---|---|---|---|---|---|---|---|---|---|---|
| PSN34458.1 | *Bger34458* | V | 1 | 350 | 4.66978E-36 | 366 | 1 | 9 | GFS | - | Abs. | FPSITIC | A | Abs. | 1 | 324 | 11 |
| PSN34460.1 | *Bger34460* | V | 38 | 329 | 4.43405E-13 | 346 | 1 | 6 | GFS | - | SSLHG | Abs. | A | GLCYTAN | 51 | 253 | 12 |
| PSN38565.1 | *Bger38565* | V | 56 | 363 | 3.29313E-33 | 370 | 1 | 8 | Abs. | W | TSLHG | FPAVTIC | - | ADYVYNT | 77 | 267 | - |
| PSN38575.1 | *Bger38575* | V | 124 | 311 | 5.27532E-13 | 327 | 0 | 10 | Abs. | - | KKI | Abs. | - | NERYCCT | 18 | 307 | - |
| PSN38576.1 | *Bger38576* | V | 126 | 380 | 3.26504E-15 | 410 | 1 | 10 | GFS | - | KKI | Abs. | A | NERYCCT | 20 | 338 | 25 |
| PSN54013.1 | *Bger4013* | I | 105 | 168 | 0.000018457 | 227 | 2 | 3 | GAS | - | LDG | Abs. | V | GRSY | 1 | 148 | 52 |
| PSN41206.1 | *Bger41206* | IV | 52 | 203 | 9.02988E-23 | 261 | 2 | 5 | GGS | - | Abs. | Abs. | A | Abs. | 2 | 178 | 52 |
| PSN41668.1 | *Bger41668* | I | 131 | 280 | 5.96391E-16 | 295 | 2 | 9 | GAS | - | Abs. | RNT | V | Abs. | 73 | 164 | 10 |
| PSN41670.1 | *Bger41670* | II | 93 | 182 | 0.000171777 | 438 | 2 | 6 | GKY | V | Abs. | KNTNCSSAALTIC | - | Abs. | 1 | 295 | 89 |
| PSN42712.1 | *Bger42712* | IV | 6 | 262 | 3.45154E-32 | 307 | 0 | 9 | GFS | - | Abs. | Abs. | G | GYCCSFN | 1 | 240 | 40 |
| PSN42805.1 | *Bger42805* | V | 14 | 119 | 0.007056 | 201 | 0 | 5 | SEE | - | Abs. | FPAVTVC | S | Abs. | 1 | 199 | - |
| PSN49921.1 | *Bger49921* | V | 36 | 166 | 4.19869E-17 | 198 | 0 | 4 | Abs. | - | Abs. | Abs. | - | Abs. | 1 | 175 | 2 |
| PSN50549.1 | *Bger50549* | III | 52 | 391 | 2.10817E-47 | 399 | 1 | 8 | KYN | D | Abs. | FPSVTLC | - | PQCQEMF | 74 | 299 | - |
| PSN52920.1 | *Bger52920* | I | 25 | 312 | 1.1455E-09 | 369 | 4 | 3 | GAS | W | SSLHG | Abs. | V | Abs. | 48 | 94 | 50 |
| PSN54949.1 | *Bger54949* | V | 29 | 243 | 2.95473E-36 | 271 | 0 | 10 | GFP | - | Abs. | MT | G | Abs. | 1 | 220 | 7 |
| PSN55271.1 | *Bger55271* | III | 2 | 180 | 1.48254E-06 | 563 | 2 | 7 | GIS | L | Abs. | FPAVTIC | V | GFCYSFN | 172 | 241 | 47 |
| PSN55272.1 | *Bger55272* | III | 42 | 288 | 1.75416E-44 | 326 | 2 | 10 | GVS | - | Abs. | Abs. | V | Abs. | 2 | 292 | 33 |
| PSN55273.1 | *Bger55273* | III | 50 | 205 | 1.35735E-14 | 225 | 1 | 1 | GNS | W | TTLHG | FPAITIC | S | Abs. | 72 | 265 | - |
| PSN55275.1 | *Bger55275* | III | 13 | 427 | 1.66249E-46 | 483 | 3 | 12 | GVS | - | Abs. | FPAITIC | V | GFCYSFN | 5 | 127 | 51 |
| PSN57566.1 | *Bger57566* | III | 116 | 405 | 1.72977E-07 | 406 | 0 | 9 | Abs. | - | Abs. | SPSIVGELKDFC | - | GFCYSFN | 42 | 384 | - |
| PSN57569.1 | *Bger57569* | III | 1 | 272 | 1.1095E-28 | 287 | 1 | 11 | Abs. | - | Abs. | Abs. | G | GFCYSFN | 1 | 339 | 4 |
| PSN57593.1 | *Bger57593* | III | 58 | 177 | 0.00304282 | 293 | 1 | 3 | Abs. | K | Abs. | FPAITIC | - | FFCTNYN | 54 | 278 | - |
| PSN57594.1 | *Bger57594* | III | 5 | 202 | 5.32629E-22 | 391 | 0 | 12 | Abs. | - | Abs. | Abs. | G | GFCYSFN | 1 | 213 | 2 |
| PSN57595.1 | *Bger57595* | III | 1 | 403 | 1.56897E-45 | 404 | 0 | 12 | Abs. | - | Abs. | FPAITIC | G | GFCYSFN | 2 | 387 | 2 |
| PSN57599.1 | *Bger57599* | III | 8 | 205 | 1.216E-23 | 210 | 0 | 8 | Abs. | - | Abs. | Abs. | - | GFCYSFN | 1 | 386 | - |
| PSN57602.1 | *Bger57602* | III | 103 | 195 | 0.000846843 | 276 | 0 | 3 | Abs. | - | Abs. | IMWIRSHKIN | - | GFCYSFN | 23 | 208 | - |
| PSN58038.1 | *Bger58038* | IV | 1 | 349 | 8.33844E-68 | 359 | 1 | 12 | Abs. | W | Abs. | FPAVTIC | - | GFCCSFN | 1 | 226 | - |
| PSN43586.1 | *BgerPPK13a* | I | 1 | 321 | 1.20498E-32 | 368 | 2 | 8 | GAS | - | Abs. | FPSVVLC | V | Abs. | 1 | 334 | 42 |
| PSN55529.1 | *BgerPPK13b* | I | 12 | 478 | 2.60071E-38 | 552 | 2 | 14 | GAS | W | TSFHA | FPSVVVC | I | GTCYAIN | 34 | 287 | 69 |
| PSN58104.1 | *BgerPPK16* | IV | 38 | 311 | 1.02438E-40 | 342 | 2 | 13 | GFS | - | SGLNE | Abs. | A | GYCCSFN | 1 | 398 | 26 |
| PSN44504.1 | *BgerPPK17* | VII | 8 | 202 | 2.1e-05 | 231 | 0 | 4 | Abs. | - | Abs. | FPAVTIC | - | GQCHTVL | 1 | 289 | - |
| PSN38910.1 | *BgerPPK23a* | VI | 63 | 467 | 4.06669E-51 | 572 | 2 | 9 | GFS | W | STLHG | FPTVSLC | V | GYCFSFN | 85 | 212 | 100 |
| PSN38909.1 | *BgerPPK23b* | VI | 78 | 348 | 1.99456E-18 | 435 | 0 | 9 | Abs. | - | TTLHG | FPAITIC | - | Abs. | 99 | 336 | 82 |

| | | | | | | | | | | | | | | | | | |
|---|---|---|---|---|---|---|---|---|---|---|---|---|---|---|---|---|---|
| PSN50939.1 | *BgerPPK28* | V | 105 | 547 | 1.09832E-71 | 609 | 2 | 14 | GFS | W | SSLHG | FPAVTIC | S | GICCSFN | 127 | 235 | 35 |
| PSN31707.1 | *Bger31707* | II | 10 | 479 | 6.00661E-35 | 509 | 3 | 12 | GCS | W | MND | FPAFTIC | V | Abs. | 10 | 396 | 25 |
| PSN51114.1 | *Bger51114* | II | 72 | 208 | 7.97653E-14 | 214 | 1 | 1 | Abs. | W | VSIHG | FPTGTIC | - | Abs. | 94 | 423 | - |
| PSN53854.1 | *BgerPPK9* | II | 1 | 222 | 1.94335E-10 | 243 | 0 | 7 | Abs. | - | Abs. | LPSFYIC | - | GLCYAYN | 1 | 237 | - |
| XP_012239254.1 | *Bimp012239254* | IV | 34 | 497 | 1.01037E-87 | 570 | 2 | 13 | GGS | W | TGLHG | FPGITIC | A | GFCCGFN | 55 | 396 | 68 |
| XP_024222992.1 | *Bimp024222992.1* | IV | 63 | 473 | 3.60545E-48 | 487 | 0 | 14 | Abs. | W | TSVHG | FPAVTIC | - | GLCCSFN | 87 | 351 | - |
| XP_012246752.1 | *Bimp012246752* | I | 15 | 409 | 1.80798E-46 | 421 | 2 | 12 | GAS | W | CTIHG | FPTVAFC | V | GYCIAMN | 37 | 317 | 4 |
| XP_003493499.1 | *Bimp003493499.1* | IV | 7 | 340 | 2.10387E-46 | 383 | 1 | 13 | GIS | - | Abs. | Abs. | G | GFCCIFN | 2 | 358 | 38 |
| XP_033179584.1 | *Bimp033179584.1* | IV | 79 | 446 | 2.19139E-43 | 493 | 1 | 12 | GFS | - | TKLAG | Abs. | G | GFCCTFN | 67 | 344 | 42 |
| XP_012242038.1 | *Bimp012242038* | II | 68 | 480 | 2.8671E-36 | 505 | 4 | 12 | GAS | - | SSVHG | FVTLKPALFIC | T | GACLTTR | 90 | 329 | 20 |
| XP_024224189.1 | *BimpPPK16* | IV | 31 | 500 | 1.41039E-71 | 528 | 2 | 14 | GFS | W | TGLHG | FPAVTVC | A | GLCCSFN | 53 | 408 | 16 |
| XP_012240062.1 | *BimpPPK23* | VI | 23 | 395 | 3.99745E-32 | 439 | 2 | 13 | GYS | W | NTLHG | FPQIFVC | V | Abs. | 45 | 304 | 39 |
| XP_012237817.1 | *BimpPPK28* | V | 147 | 620 | 9.7856E-99 | 658 | 2 | 14 | GFS | W | STLHG | FPSVTIC | S | GMCCNFN | 169 | 405 | 33 |
| XP_004932266.1 | *Bmor004932266.1* | IV | 56 | 451 | 2.12569E-47 | 476 | 1 | 14 | Abs. | W | SSICG | FPAVGIC | - | GYCCTFN | 78 | 354 | - |
| XP_012544414.1 | *Bmor012544414.1* | IV | 13 | 251 | 7.96828E-22 | 268 | 2 | 10 | GFS | - | Abs. | Abs. | S | VFLFSPG | 1 | 372 | 12 |
| XP_012544908.1 | *Bmor012544908.1* | IV | 37 | 227 | 1.07922E-23 | 236 | 1 | 3 | Abs. | W | SSIGG | FPAVAFC | - | Abs. | 59 | 229 | - |
| XP_012545602.1 | *Bmor012545602.1* | IV | 24 | 398 | 2.88218E-36 | 601 | 1 | 12 | GGS | - | MDG | FPALTIC | A | GYCCQFD | 3 | 151 | 198 |
| XP_012545987.1 | *Bmor012545987.1* | IV | 64 | 353 | 1.15797E-07 | 372 | 0 | 13 | TET | Y | TTLHG | LPLTLA | S | GLCCVLR | 29 | 353 | 19 |
| XP_012545990.1 | *Bmor012545990.1* | IV | 12 | 394 | 8.1522E-29 | 439 | 2 | 6 | GVS | W | AQFHG | FPSVWIC | S | GFCCVFN | 24 | 320 | 35 |
| XP_012550654.1 | *Bmor012550654.1* | V | 79 | 574 | 1.47802E-98 | 611 | 2 | 14 | GFS | W | SNLHG | YPAVTLC | A | GLCYTFN | 101 | 324 | 32 |
| XP_012552731.1 | *Bmor012552731.1* | V | 3 | 415 | 6.1755E-50 | 456 | 2 | 14 | GFS | W | Abs. | LQDVSMIC | A | GVCYTMN | 2 | 427 | 15 |
| XP_012553296.1 | *Bmor012553296.1* | I | 20 | 433 | 1.0828E-43 | 440 | 3 | 13 | GAS | W | TSLSG | FPAVTIC | V | GLCYVFN | 43 | 388 | 2 |
| XP_004929670.1 | *Bmor004929670.1* | II | 1 | 354 | 2.39705E-13 | 356 | 0 | 12 | RIS | - | Abs. | YPAIILC | P | GMCHFIN | 1 | 344 | - |
| XP_004932267.1 | *BmorPPK13a* | I | 13 | 434 | 1.01116E-57 | 480 | 2 | 14 | GAS | W | CTFAG | LPAVAIC | V | GTCFIIN | 35 | 353 | 41 |
| XP_012550884.1 | *BmorPPK13b* | I | 24 | 440 | 4.73019E-54 | 509 | 2 | 14 | GAS | W | TSIHG | FPAVAVC | V | GTCYAIN | 43 | 352 | 63 |
| XP_012548978.1 | *BmorPPK16* | IV | 2 | 318 | 8.49348E-46 | 379 | 2 | 11 | GFT | - | Abs. | Abs. | A | NLCCTFN | 1 | 296 | 56 |
| XP_012545187.1 | *BmorPPK17* | VII | 28 | 375 | 2.13569E-07 | 414 | 2 | 8 | KPG | V | PESS | YPSLTFC | S | GRCYTMN | 14 | 325 | 21 |
| XP_012546079.1 | *BmorPPK25* | II | 5 | 328 | 3.42793E-19 | 345 | 0 | 12 | GCS | - | Abs. | LPAVTAC | - | GVCYSTN | 1 | 321 | - |
| XP_012548124.1 | *BmorPPK28* | V | 24 | 475 | 4.27685E-80 | 494 | 2 | 13 | GFS | W | Abs. | FPAITIC | S | GLCCTFN | 27 | 402 | 14 |
| XP_012546099.1 | *Bmor012546099.1* | II | 5 | 253 | 1.92667E-12 | 299 | 1 | 7 | Abs. | - | Abs. | FPAVTAC | - | GMCLTYN | 1 | 353 | 13 |
| XP_012550136.1 | *BmorPPK9* | II | 47 | 468 | 5.01913E-57 | 480 | 2 | 12 | GCS | W | SSIHG | FPCVTIC | V | GLCYAVN | 69 | 259 | 7 |
| XP_025263766.1 | *Cflo025263766* | III | 115 | 529 | 1.1355E-47 | 599 | 2 | 12 | GGS | W | TTLHG | FPSVTLC | V | GFCYSFN | 137 | 346 | 65 |
| XP_025266050.1 | *Cflo025266050* | IV | 59 | 544 | 7.51742E-58 | 559 | 2 | 14 | GAS | W | SSLHG | FPAITVC | A | GLCCSFN | 83 | 415 | 10 |

| | | | | | | | | | | | | | | | | | |
|---|---|---|---|---|---|---|---|---|---|---|---|---|---|---|---|---|---|
| XP_025266757.1 | *Cflo025266757* | IV | 43 | 387 | 3.00142E-46 | 543 | 1 | 9 | Abs. | W | SSLSS | FPAIAIC | - | GFCCTFN | 65 | 436 | 12 |
| XP_025269475.1 | *Cflo025269475* | IV | 8 | 472 | 4.26647E-87 | 556 | 2 | 13 | GGS | W | TGLHG | FPAITIC | A | GFCCAFN | 30 | 396 | 79 |
| XP_025270836.1 | *CfloPPK16*[a] | IV | 28 | 426 | 5.01781E-68 | 483 | 1 | 14 | Abs. | W | TGLHG | FPAITIC | - | GLCCSFN | 50 | 390 | 3 |
| XP_025270842.1 | *CfloPPK16b* | IV | 28 | 442 | 1.54508e-68 | 443 | 1 | 14 | Abs. | W | TGLHG | FPAITIC | - | GLCCSFN | 50 | 367 | - |
| XP_025270843.1 | *CfloPPK16c* | IV | 28 | 475 | 3.88642e-76 | 482 | 1 | 14 | Abs. | W | TGLHG | FPAITIC | - | GLCCSFN | 50 | 403 | 3 |
| XP_011269762.1 | *CfloPPK28* | V | 168 | 622 | 4.75319E-85 | 682 | 2 | 14 | GFS | W | STLHG | FPSITIC | S | GICCNFN | 190 | 408 | 33 |
| CLEC000454-PA | *Clec000454* | I | 3 | 340 | 1.02492E-23 | 635 | 2 | 9 | GAS | - | Abs. | FPTVTIC | V | GVCYSLN | 259 | 291 | 37 |
| CLEC003170-PA | *Clec003170* | I | 11 | 431 | 8.98173E-66 | 485 | 2 | 14 | GAS | W | TSFHG | FPAISVC | V | GTCFSFN | 33 | 352 | 49 |
| CLEC004707-PA | *Clec004707* | III | 9 | 223 | 1.53734E-21 | 225 | 1 | 4 | Abs. | W | STVHC | FPAVTLC | - | GFCYSFN | 52 | 146 | - |
| CLEC004780-PA | *Clec004780* | III | 41 | 489 | 1.40463E-39 | 597 | 2 | 11 | GGS | W | STLHG | FPAVSIC | V | IVCHKIQ | 63 | 400 | 83 |
| CLEC006346-PA | *Clec006346* | V | 1 | 333 | 2.71193E-39 | 369 | 1 | 12 | GVS | - | Abs. | Abs. | A | GMCFSFN | 1 | 311 | 31 |
| CLEC012452-PA | *ClecPPK23* | VI | 63 | 281 | 1.39762E-29 | 289 | 0 | 13 | Abs. | - | Abs. | Abs. | - | GFCYSFN | 8 | 279 | 1 |
| CLEC000413-PA | *ClecPPK28* | V | 26 | 439 | 8.83025E-53 | 497 | 2 | 13 | Abs. | W | STL | FPAITIC | C | GMCCVFN | 26 | 391 | 38 |
| CLEC004429-PA | *Clec004429* | II | 16 | 454 | 4.05095E-53 | 460 | 2 | 12 | GCS | W | TTLHG | FPASTAC | V | GYCYSYS | 38 | 370 | 1 |
| CLEC000460-PA | *ClecPPK9a* | II | 7 | 366 | 6.1699E-26 | 376 | 0 | 10 | NRR | W | Abs. | FPAVSLC | V | GLCYVYN | 1 | 351 | 5 |
| CLEC000458-PA | *ClecPPK9b* | II | 1 | 309 | 1.14007E-32 | 316 | 0 | 12 | Abs. | - | Abs. | FPASTIC | - | GLCYSYN | 2 | 310 | 1 |
| CPIJ002355 | *Cqui002355* | IV | 75 | 543 | 2.81723E-78 | 580 | 2 | 14 | GLS | W | ITTHC | FPAISLC | S | GFCCTFN | 95 | 402 | 32 |
| CPIJ003410 | *Cqui003410* | V | 42 | 503 | 2.9717E-121 | 569 | 2 | 14 | GVS | W | STIHG | FPAVTIC | A | GLCFTFN | 64 | 393 | 61 |
| CPIJ003411 | *Cqui003411* | V | 23 | 438 | 1.86298E-86 | 477 | 2 | 14 | GVS | - | HVI | FPAVTIC | A | GLCYTFN | 21 | 386 | 34 |
| CPIJ005124 | *Cqui005124* | V | 58 | 541 | 1.02737E-97 | 571 | 2 | 14 | GAS | W | TTIHG | FPAITVC | A | GICFTYN | 79 | 416 | 25 |
| CPIJ005125 | *Cqui005125* | V | 12 | 485 | 8.48688E-90 | 518 | 2 | 14 | GIS | W | TTIHG | FPAITVC | A | GICFTYN | 33 | 406 | 28 |
| CPIJ005126 | *Cqui005126* | V | 55 | 533 | 5.62273E-95 | 577 | 2 | 14 | GVS | W | STIHG | FPAITIC | A | GICMTFN | 77 | 422 | 27 |
| CPIJ005127 | *Cqui005127* | V | 52 | 530 | 2.24536E-94 | 568 | 2 | 14 | GVS | W | STIHG | FPAVTIC | A | GICMTFN | 74 | 422 | 21 |
| CPIJ005128 | *Cqui005128* | V | 51 | 503 | 1.06521E-80 | 545 | 2 | 12 | GVS | W | SSIHG | FPAVTIC | A | GICYTLN | 73 | 396 | 25 |
| CPIJ005129 | *Cqui005129* | V | 35 | 518 | 3.28595E-96 | 540 | 2 | 14 | GFS | W | STIHG | FPAVTIC | A | GICYTYN | 56 | 416 | 17 |
| CPIJ005131 | *Cqui005131* | V | 28 | 440 | 5.73784E-86 | 532 | 3 | 11 | GIS | W | STIHG | FPAITVC | A | Abs. | 49 | 345 | 87 |
| CPIJ005430 | *Cqui005430* | IV | 31 | 412 | 2.71836E-54 | 500 | 2 | 7 | GGS | W | TSLHG | FPAVTIC | A | GFCCSFN | 59 | 307 | 83 |
| CPIJ005908 | *Cqui005908* | V | 19 | 482 | 2.98809E-87 | 543 | 2 | 12 | GVS | W | TSIHG | YPAVTIC | A | GICYTFN | 41 | 395 | 56 |
| CPIJ005909 | *Cqui005909* | V | 19 | 489 | 7.69182E-87 | 573 | 2 | 12 | GVS | W | TSIHG | YPALTIC | A | GICYTFN | 41 | 414 | 67 |
| CPIJ007799 | *Cqui007799* | V | 37 | 497 | 7.76399E-67 | 520 | 2 | 12 | GAS | W | NFVTA | FPAITVC | A | GICYTFN | 40 | 411 | 18 |
| CPIJ007800 | *Cqui007800* | V | 1 | 409 | 2.89543E-53 | 469 | 1 | 14 | GAS | - | Abs. | FPAITMC | A | GICYTFN | 1 | 402 | 36 |
| CPIJ007801 | *Cqui007801* | V | 27 | 496 | 1.24068E-72 | 533 | 2 | 14 | GAS | W | NFVYA | FPAITIC | A | GICYTFN | 40 | 410 | 32 |
| CPIJ007802 | *Cqui007802* | V | 28 | 421 | 1.56779E-55 | 924 | 5 | 14 | GAS | W | NFVYA | FPAVTIC | A | GICYTFN | 482 | 358 | 33 |
| CPIJ007803 | *Cqui007803* | V | 18 | 490 | 1.21002E-84 | 515 | 2 | 14 | GAS | W | SSVYA | FPAVTIC | A | GICYTFN | 40 | 404 | 20 |

| | | | | | | | | | | | | | | | | | |
|---|---|---|---|---|---|---|---|---|---|---|---|---|---|---|---|---|---|
| CPIJ007804 | *Cqui007804* | V | 13 | 478 | 1.70791E-66 | 507 | 2 | 12 | GAS | W | GTVYA | FPAVTLC | A | GLCFTFN | 35 | 397 | 24 |
| CPIJ007805 | *Cqui007805* | V | 1 | 399 | 2.44459E-60 | 432 | 1 | 14 | GVS | - | Abs. | FPAITIC | A | GICYTFN | 1 | 386 | 11 |
| CPIJ007806 | *Cqui007806* | V | 34 | 516 | 5.5953E-113 | 565 | 2 | 14 | GVS | W | TSIHG | FPAVTIC | A | GFCYTFN | 55 | 415 | 44 |
| CPIJ007807 | *Cqui007807* | V | 5 | 466 | 1.11616E-87 | 505 | 2 | 14 | GIS | W | Abs. | FPSVTIC | A | GICYTFN | 5 | 415 | 34 |
| CPIJ007808 | *Cqui007808* | V | 20 | 501 | 4.80818E-91 | 541 | 2 | 14 | GIS | W | SSIHG | FPSVTIC | A | GMCYTFN | 42 | 413 | 35 |
| CPIJ007809 | *Cqui007809* | V | 23 | 506 | 2.7636E-111 | 557 | 2 | 14 | GIS | W | SSIHG | FPSVTIC | A | GICYTFN | 45 | 415 | 46 |
| CPIJ007810 | *Cqui007810* | V | 29 | 499 | 8.60292E-89 | 530 | 2 | 14 | GVS | - | TSIHG | FPAITIC | A | GFCFTFN | 51 | 410 | 26 |
| CPIJ007811 | *Cqui007811* | V | 17 | 504 | 1.99823E-74 | 531 | 2 | 14 | GGT | W | TSVHG | FPAVTIC | A | GVCYNFN | 39 | 419 | 22 |
| CPIJ008773 | *Cqui008773* | II | 3 | 309 | 3.97319E-18 | 309 | 1 | 11 | Abs. | W | Abs. | FPSLTVC | - | Abs. | 2 | 281 | - |
| CPIJ010135 | *Cqui010135* | V | 43 | 461 | 8.74483E-47 | 469 | 2 | 11 | GVT | W | TSFFG | FPAVTIC | A | GVCYSFN | 65 | 350 | 3 |
| CPIJ010136 | *Cqui010136* | V | 67 | 538 | 6.40492E-66 | 563 | 2 | 11 | GVS | W | SSIHG | FPAVTIC | G | GICYSFN | 89 | 403 | 20 |
| CPIJ012543 | *Cqui012543* | V | 20 | 511 | 8.061E-106 | 551 | 2 | 14 | GIS | W | SSIHG | FPAVTIC | A | GVCATFN | 42 | 423 | 35 |
| CPIJ012544 | *Cqui012544* | V | 2 | 461 | 1.08322E-94 | 501 | 2 | 14 | GIS | - | Abs. | FPAVTIC | A | GICATFN | 1 | 423 | 35 |
| CPIJ012545 | *Cqui012545* | V | 21 | 513 | 2.5198E-106 | 547 | 2 | 14 | GVS | W | SSVHG | FPAITIC | A | GVCVTFN | 43 | 424 | 29 |
| CPIJ012546 | *Cqui012546* | V | 23 | 497 | 2.70326E-87 | 544 | 2 | 14 | GVS | W | SSVHG | LPAVTIC | A | GVCVTFN | 45 | 418 | 30 |
| CPIJ017961 | *Cqui017961* | V | 51 | 267 | 3.88574E-27 | 272 | 1 | 11 | Abs. | W | SSIHC | FPAVTIC | - | Abs. | 73 | 170 | 3 |
| CPIJ006134 | *CquiPPK10* | II | 37 | 138 | 3.40507E-12 | 414 | 1 | 2 | Abs. | W | GALHG | MPSLTIC | - | GICYTTN | 59 | 329 | - |
| CPIJ019036 | *CquiPPK13* | I | 3 | 306 | 1.05831E-17 | 730 | 2 | 14 | GAS | W | EVLNE | FPAVVVC | V | GLCYAVN | 306 | 354 | 19 |
| CPIJ001728 | *CquiPPK15* | I | 45 | 428 | 8.23795E-50 | 465 | 2 | 14 | GAS | L | Abs. | FPSVGIC | V | GSCYLIN | 33 | 364 | 32 |
| CPIJ012002 | *CquiPPK16* | IV | 52 | 513 | 1.92222E-92 | 529 | 2 | 14 | GFS | W | TALHG | FPAVTIC | A | GLCCSFN | 73 | 394 | 11 |
| CPIJ000851 | *CquiPPK17* | VII | 14 | 366 | 0.250709 | 436 | 2 | 10 | GLS | S | Abs. | YPAITFC | A | GQCFTMN | 14 | 312 | 45 |
| CPIJ017580 | *CquiPPK23a* | VI | 28 | 402 | 2.23132E-59 | 547 | 3 | 13 | GFS | W | STLHG | FPTVLVC | V | Abs. | 50 | 306 | 140 |
| CPIJ017584 | *CquiPPK23b* | VI | 47 | 468 | 1.40501E-76 | 610 | 2 | 14 | GFS | W | STLHG | FPTVMVC | V | GFCYSFN | 69 | 353 | 137 |
| CPIJ000103 | *CquiPPK25* | II | 8 | 277 | 3.90032E-16 | 277 | 1 | 3 | Abs. | W | SSVHG | PPAVTIC | - | GMCFTSS | 29 | 222 | - |
| CPIJ003913 | *CquiPPK3* | II | 184 | 444 | 1.64162E-32 | 461 | 2 | 11 | GKN | - | Abs. | LAVEGLRNV | V | GICYALN | 1 | 412 | 11 |
| CPIJ007315 | *CquiPPK301* | V | 45 | 485 | 2.93192E-85 | 574 | 2 | 14 | GFS | F | Abs. | FPAVTIC | S | GLCCTFN | 44 | 395 | 12 |
| CPIJ015546 | *CquiPPK31* | I | 8 | 393 | 5.72717E-41 | 414 | 2 | 14 | GAS | W | Abs. | FPAVSVC | I | GRCFSVN | 9 | 343 | 84 |
| CPIJ018767 | *CquiPPK6a* | IV | 48 | 334 | 1.40377E-22 | 449 | 2 | 5 | GMS | W | TKVNG | FPGITIC | A | GACCSFN | 70 | 243 | 85 |
| CPIJ005031 | *CquiPPK6b* | IV | 48 | 424 | 3.50942E-39 | 464 | 2 | 11 | GMS | W | TKVNG | FPAITIC | S | GACCSFN | 70 | 308 | 35 |
| CPIJ007762 | *CquiPPK9* | II | 15 | 356 | 1.46128E-31 | 358 | 1 | 11 | Abs. | W | TSAHG | FPSLTVC | - | GSCVSVN | 36 | 295 | - |
| CG3478 | *DmelPPK1* | V | 69 | 574 | 1.65603E-91 | 606 | 2 | 14 | GIS | W | TTIHG | FPTITIC | S | GICYQFN | 91 | 437 | 32 |
| CG34042 | *DmelPPK10* | II | 22 | 471 | 6.00247E-56 | 485 | 4 | 12 | GFS | W | CCIHG | LPSVTVC | V | GLCYLGN | 44 | 381 | 27 |
| CG34058 | *DmelPPK11* | IV | 96 | 479 | 1.12498E-52 | 516 | 2 | 14 | GCS | W | ASIHG | FPAVTIC | G | RLCCSFN | 116 | 317 | 9 |
| CG10972 | *DmelPPK12* | V | 29 | 499 | 3.98379E-92 | 569 | 2 | 14 | GFS | F | TSLHG | FPAITIC | S | GLCCVFN | 51 | 402 | 65 |

| | | | | | | | | | | | | | | | | | |
|---|---|---|---|---|---|---|---|---|---|---|---|---|---|---|---|---|---|
| CG33508 | *DmelPPK13* | I | 10 | 434 | 6.6123E-64 | 460 | 2 | 14 | GAS | W | STLHG | FPAIIVC | V | GICYAIN | 32 | 356 | 21 |
| CG9501 | *DmelPPK14* | III | 45 | 483 | 3.97913E-70 | 504 | 2 | 14 | GCS | W | SHIHG | FPVVIIC | V | GYCLAFN | 67 | 370 | 16 |
| CG14239 | *DmelPPK15* | I | 27 | 466 | 6.25107E-59 | 483 | 2 | 14 | GAS | W | CTLAG | FPAVSVC | V | GRCYMLN | 49 | 371 | 12 |
| CG34059 | *DmelPPK16* | IV | 59 | 518 | 1.12797E-89 | 531 | 2 | 14 | GFS | W | TSLHG | FPAVTIC | A | GQCCTFN | 81 | 391 | 8 |
| CG13278 | *DmelPPK17* | VII | 22 | 358 | 1.21212E-07 | 431 | 2 | 10 | GLS | F | Abs. | MPSVTIC | A | GRCYTLR | 8 | 316 | 44 |
| CG44152 | *DmelPPK18* | IV | 16 | 510 | 5.60081E-35 | 553 | 2 | 12 | GFS | Y | Abs. | FPAVGVC | N | NVCCIFN | 38 | 426 | 38 |
| CG18287 | *DmelPPK19* | III | 37 | 489 | 1.66369E-69 | 511 | 2 | 14 | GAS | W | SNIHG | FPAVAVC | A | GQCLVFN | 57 | 387 | 16 |
| CG1058 | *DmelPPK2* | V | 39 | 546 | 1.8664E-102 | 562 | 2 | 14 | GFS | W | TSIHG | FPAVTVC | A | GICYTFN | 61 | 440 | 32 |
| CG7577 | *DmelPPK20* | III | 34 | 521 | 1.40066E-65 | 587 | 2 | 14 | GAS | W | STIHC | FPAVGIC | A | GFCLVFN | 78 | 397 | 10 |
| CG12048 | *DmelPPK21* | III | 51 | 513 | 1.06259E-80 | 550 | 2 | 14 | GAS | W | SSVHG | FPSIGLC | A | GLCFVFN | 75 | 392 | 61 |
| CG31105 | *DmelPPK22* | IV | 46 | 527 | 9.41524E-74 | 561 | 2 | 14 | GFS | W | ISLQG | FPAVSIC | S | GPCCTFN | 66 | 415 | 29 |
| CG8527 | *DmelPPK23* | VI | 34 | 476 | 6.48182E-67 | 502 | 4 | 14 | GFS | W | STLHG | FPTTVVC | V | GFCYAFN | 56 | 374 | 21 |
| CG15555 | *DmelPPK24* | IV | 65 | 551 | 8.26743E-73 | 595 | 2 | 14 | GFS | W | LSFHC | FPAISVC | S | GFCCTFN | 86 | 422 | 39 |
| CG33349 | *DmelPPK25* | II | 18 | 432 | 3.2717E-42 | 451 | 2 | 12 | GAS | W | SSIHG | FPFVGITLC | M | GLCQTSS | 39 | 347 | 14 |
| CG8546 | *DmelPPK26* | V | 74 | 570 | 2.4458E-109 | 597 | 2 | 14 | GVS | W | TTIHG | FPAVTVC | A | GVCFSFN | 96 | 428 | 22 |
| CG10858 | *DmelPPK27* | IV | 15 | 407 | 1.21699E-40 | 422 | 2 | 14 | GCS | W | TSLNG | FPELKIC | S | GNCCVLR | 37 | 324 | 10 |
| CG4805 | *DmelPPK28* | V | 43 | 535 | 5.50756E-83 | 632 | 2 | 14 | GFS | F | STLHG | FPAITIC | S | GLCCNFN | 90 | 398 | 92 |
| CG13568 | *DmelPPK29* | I | 8 | 414 | 8.22697E-31 | 425 | 2 | 11 | GAS | W | Abs. | FPSIGIC | V | GYCFLAN | 22 | 347 | 6 |
| CG30181 | *DmelPPK3* | II | 59 | 464 | 9.96815E-57 | 535 | 4 | 12 | GVS | W | TGLLA | LPAVTLC | V | GACYAIN | 68 | 352 | 9 |
| CG18110 | *DmelPPK30* | III | 47 | 446 | 2.96293E-41 | 457 | 2 | 12 | GIS | - | Abs. | FPAVTII | V | GFAFEFN | 38 | 361 | 64 |
| CG31065 | *DmelPPK31* | I | 40 | 387 | 2.50847E-34 | 421 | 3 | 14 | NTS | L | Abs. | FPAITVC | I | GLCFSFN | 3 | 342 | 10 |
| CG8178 | *DmelPPK4* | IV | 30 | 468 | 2.42416E-82 | 516 | 2 | 14 | GMS | W | SSIHG | FPPVTIC | S | GHCCSFN | 52 | 370 | 43 |
| CG33289 | *DmelPPK5* | V | 4 | 405 | 1.09789E-54 | 405 | 1 | 14 | Abs. | F | MEM | FPTITVC | - | GLCCIFN | 18 | 361 | - |
| CG11209 | *DmelPPK6* | IV | 85 | 476 | 5.04492E-47 | 498 | 2 | 14 | GLS | W | TKVSG | FPGVTIC | S | GPCCSFN | 63 | 367 | 17 |
| CG9499 | *DmelPPK7* | III | 38 | 493 | 1.57375E-80 | 573 | 2 | 14 | GCS | W | TTIHG | FPVITIC | V | GLCWAFN | 60 | 387 | 75 |
| CG32792 | *DmelPPK8* | V | 42 | 508 | 9.46536E-93 | 569 | 2 | 14 | GFS | F | TTIHG | FPAVTIC | S | GLCCVFN | 64 | 398 | 56 |
| CG34369 | *DmelPPK9* | II | 154 | 587 | 1.14405E-55 | 599 | 3 | 12 | GCS | W | SSIHG | FPSLTVC | V | GICHSVN | 176 | 365 | 7 |
| DPOGS204183 | *Dplex204183* | IV | 15 | 405 | 1.18732E-44 | 434 | 1 | 10 | Abs. | W | GVVEI | Abs. | - | IFIFESS | 15 | 392 | - |
| DPOGS204735 | *Dplex204735* | IV | 142 | 221 | 0.00296715 | 241 | 0 | 6 | Abs. | - | AQFHG | Abs. | - | Abs. | 20 | 212 | 5 |
| DPOGS204741 | *Dplex204741* | IV | 13 | 86 | 0.000023284 | 188 | 1 | 2 | Abs. | W | TTLHG | LPTVAVC | - | Abs. | 33 | 129 | - |
| DPOGS209994 | *Dplex209994* | IV | 9 | 281 | 1.08669E-22 | 297 | 0 | 11 | Abs. | - | Abs. | Abs. | S | GFCCVFN | 1 | 279 | 16 |
| DPOGS210747 | *Dplex210747* | I | 59 | 219 | 2.37762E-15 | 638 | 3 | 12 | GAS | W | SNLHG | FPGVTVC | V | GSCYVFN | 261 | 325 | 1 |
| DPOGS214762 | *Dplex214762* | IV | 36 | 349 | 3.49155E-09 | 438 | 1 | 11 | GRQ | Y | SSIKG | FPAVGLC | S | GLQVVIK | 58 | 194 | 47 |
| DPOGS215856 | *Dplex215856* | I | 26 | 246 | 7.34511E-26 | 262 | 2 | 10 | GG | - | Abs. | Abs. | V | Abs. | 1 | 286 | 11 |

| | | | | | | | | | | | | | | | | | |
|---|---|---|---|---|---|---|---|---|---|---|---|---|---|---|---|---|---|
| | | | | | | | | | S | | | | | | | | |
| DPOGS216040 | *Dplex216040* | V | 110 | 201 | 3.53199E-18 | 310 | 0 | 6 | Abs. | - | SNLHG | PAQSA | - | Abs. | 68 | 224 | 13 |
| DPOGS216041 | *Dplex216041* | V | 43 | 515 | 1.81236E-68 | 1034 | 2 | 13 | Abs. | W | SNLHG | YPAVTIC | - | GVCYTFN | 64 | 207 | 514 |
| DPOGS203427 | *DplexPPK13a* | I | 10 | 241 | 3.95954E-10 | 286 | 1 | 6 | Abs. | W | CSFGG | LPTVAIC | - | Abs. | 32 | 428 | - |
| DPOGS202727 | *DplexPPK13b* | I | 44 | 282 | 2.49064E-15 | 728 | 1 | 12 | Abs. | W | SSICG | FPSIAVC | - | GPCFIIN | 46 | 217 | 3 |
| DPOGS200906 | *DplexPPK16* | IV | 153 | 551 | 8.74084E-55 | 613 | 2 | 12 | GFT | R | RNTLHR | FPAVTIC | A | Abs. | 36 | 288 | 57 |
| DPOGS215595 | *DplexPPK23* | VI | 52 | 457 | 5.18464E-62 | 507 | 2 | 14 | Abs. | W | STLHG | FPAVTIC | - | GFCYAFN | 74 | 455 | 44 |
| DPOGS207910 | *DplexPPK25* | II | 8 | 347 | 1.89794E-37 | 376 | 2 | 11 | GAS | W | TSFHG | MPAVTAC | V | Abs. | 30 | 363 | 24 |
| DPOGS206892 | *DplexPPK28* | V | 46 | 440 | 3.64686E-68 | 502 | 2 | 14 | GPV | W | STLHG | FPAITIC | L | GLCCTFN | 68 | 271 | 9 |
| DPOGS211506 | *Dplex211506* | II | 37 | 215 | 3.87061E-16 | 279 | 1 | 2 | Abs. | W | TTLHG | FPAVTAC | - | GLCTTFN | 59 | 374 | - |
| DPOGS214642 | *DplexPPK6* | IV | 2 | 283 | 7.27264E-22 | 297 | 0 | 14 | Abs. | - | Abs. | YPAITIC | - | GMCCIFN | - | 296 | - |
| DPOGS210227 | *DplexPPK9* | II | 65 | 264 | 1.06889E-34 | 276 | 2 | 11 | GCS | - | MVVHG | VPLSVLTDC | V | Abs. | 19 | 210 | 7 |
| ENN71305 | *Dpon71305* | V | 41 | 443 | 4.47085E-61 | 484 | 2 | 9 | GFS | W | TGIHG | FPAVTIC | A | GLCFTFN | 62 | 335 | 36 |
| ENN76926 | *Dpon76926* | V | 41 | 431 | 1.99598E-71 | 464 | 2 | 11 | GFS | - | TGIHG | Abs. | A | GLCYTFN | 78 | 332 | 28 |
| ENN81350 | *Dpon81350* | II | 124 | 275 | 0.000870684 | 319 | 0 | 9 | Abs. | W | Abs. | ISGVSYIC | T | Abs. | 6 | 272 | 16 |
| ENN81697 | *Dpon81697* | V | 27 | 426 | 1.13574E-66 | 448 | 2 | 11 | GVS | W | SSIHG | FPAVTIC | A | Abs. | 48 | 332 | 17 |
| ENN81698 | *Dpon81698* | V | 18 | 463 | 5.03735E-55 | 497 | 2 | 12 | GFS | W | SFILG | FPAVTIC | G | GICATFN | 18 | 399 | 29 |
| ENN81481 | *DponPPK13* | I | 27 | 170 | 0.000069552 | 181 | 1 | 6 | Abs. | W | MG | FPAVFVC | - | Abs. | 26 | 128 | 1 |
| ENN77435 | *DponPPK16a* | IV | 1 | 273 | 7.73628E-21 | 273 | 1 | 7 | Abs. | W | Abs. | FPAVTIC | S | Abs. | 1 | 233 | 12 |
| ENN77436 | *DponPPK16b* | IV | 89 | 227 | 3.26519E-10 | 473 | 2 | 12 | Abs. | K | MSG | FPAATIC | S | GFCCSYN | 38 | 386 | 9 |
| ENN77437 | *DponPPK16c* | IV | 48 | 196 | 1.97127E-17 | 208 | 0 | 5 | Abs. | - | Abs. | Abs. | S | Abs. | 39 | 168 | - |
| ENN71156 | *DponPPK17* | VII | 23 | 348 | 1.32312E-07 | 453 | 1 | 10 | GLS | - | Abs. | YPSVTIC | A | GKCFSFT | 1 | 328 | 75 |
| ENN81927 | *DponPPK23* | VI | 55 | 381 | 4.111E-49 | 394 | 1 | 14 | Abs. | W | QTL | FPAVAVC | - | GLCYA | 55 | 301 | 12 |
| ENN71763 | *DponPPK28* | V | 84 | 393 | 1.47043E-54 | 393 | 1 | 5 | Abs. | W | TTLHG | FPAITIC | - | GLCCSFN | 106 | 261 | - |
| ENN73452 | *DponPPK3* | II | 118 | 201 | 0.0358152 | 213 | 0 | 6 | Abs. | - | Abs. | SGF | - | QFYHIH | 1 | 208 | - |
| ENN75495 | *DponPPK9* | II | 33 | 235 | 1.23287E-35 | 246 | 1 | 10 | GCS | - | Abs. | Abs. | V | Abs. | 1 | 210 | 6 |
| GMOY007470 | *GmorPPK10a* | II | 36 | 500 | 5.50692E-56 | 514 | 4 | 12 | GFS | W | CSIHG | LPSVTIC | V | GLCYLTN | 58 | 396 | 9 |
| GMOY004892 | *GmorPPK10b* | II | 46 | 522 | 9.33696E-49 | 534 | 2 | 12 | GFN | W | STIHG | MPSVTIC | V | GLCYMTN | 68 | 408 | 7 |
| GMOY011877 | *GmorPPK15* | I | 66 | 411 | 5.57964E-33 | 427 | 3 | 11 | GAS | W | CP | FPTISVC | V | Abs. | 70 | 295 | 11 |
| GMOY005158 | *GmorPPK17* | VII | 40 | 384 | 0.134472 | 455 | 2 | 9 | GLS | F | Abs. | MPAITIC | A | GRCFTMR | 26 | 324 | 40 |
| GMOY010902 | *GmorPPK1a* | V | 65 | 200 | 7.97528E-15 | 200 | 1 | 1 | Abs. | W | TPIHG | FPTITIC | - | Abs. | 87 | 85 | 2 |
| GMOY004489 | *GmorPPK1b* | V | 65 | 200 | 3.78286E-16 | 200 | 1 | 1 | Abs. | W | TSIHG | FPTITIC | - | Abs. | 87 | 85 | 2 |
| GMOY006517 | *GmorPPK1c* | V | 1 | 417 | 3.73056E-53 | 440 | 1 | 12 | GIS | - | Abs. | Abs. | A | GVCYQFN | 8 | 388 | 18 |

| | | | | | | | | | | | | | | | | | |
|---|---|---|---|---|---|---|---|---|---|---|---|---|---|---|---|---|---|
| GMOY002353 | *GmorPPK2* | V | 15 | 193 | 9.84149E-40 | 777 | 4 | 14 | GFS | W | TSIHG | FPAVTIC | A | GVCFTFN | 284 | 436 | 6 |
| GMOY008422 | *GmorPPK22* | IV | 44 | 460 | 9.79213E-53 | 460 | 1 | 14 | Abs. | W | ISLHG | FPAISIC | - | GPCCIFN | 65 | 369 | - |
| GMOY007016 | *GmorPPK23* | VI | 1 | 127 | 9.36091E-25 | 248 | 2 | 8 | GFS | - | Abs. | Abs. | V | Abs. | - | 106 | 116 |
| GMOY004243 | *GmorPPK26* | V | 73 | 556 | 3.3127E-107 | 583 | 2 | 14 | GVS | W | STIHG | FPTITIC | A | GMCYSFN | 95 | 415 | 22 |
| GMOY005504 | *GmorPPK28* | V | 18 | 473 | 4.05287E-82 | 516 | 2 | 14 | GFS | F | TTLHG | FPAITLC | S | GLCCTFN | 40 | 387 | 38 |
| GMOY001021 | *GmorPPK29* | I | 18 | 287 | 3.47045E-07 | 326 | 1 | 4 | Abs. | W | Abs. | FPSVTFC | - | GYCFVAN | 26 | 274 | - |
| GMOY009113 | *GmorPPK6* | IV | 23 | 387 | 5.2469E-43 | 397 | 1 | 14 | Abs. | W | TKVSG | FPGVTVC | - | GPCCSFN | 45 | 326 | - |
| GMOY001228 | *GmorPPK7* | III | 36 | 467 | 1.68857E-50 | 467 | 1 | 14 | Abs. | W | STIPG | FPVVSIC | - | GYCWSFN | 58 | 382 | 1 |
| LDEC008039 | *Ldec008039* | IV | 57 | 205 | 5.47669E-10 | 205 | 0 | 10 | Abs. | - | MLS | DPAV | - | Abs. | 1 | 203 | - |
| LDEC011575 | *Ldec011575* | V | 3 | 315 | 1.48701E-29 | 316 | 1 | 5 | Abs. | W | KTC | FPAVTIC | - | GVCYSFN | 16 | 274 | - |
| LDEC017033 | *Ldec017033* | V | 40 | 427 | 7.07871E-57 | 448 | 1 | 14 | GKH | W | TSIHG | FPAVTFC | G | SVCYTYN | 61 | 361 | - |
| LDEC017036 | *Ldec017036* | V | 89 | 190 | 2.85039E-20 | 221 | 2 | 5 | GFS | - | LTHC | Abs. | A | Abs. | 3 | 166 | 26 |
| LDEC017037 | *Ldec017037* | V | 49 | 167 | 0.000047 | 178 | 0 | 4 | Abs. | Y | TKNALSS | FSDSSLIC | - | GMFMEGE | 1 | 161 | - |
| LDEC019895 | *Ldec019895* | V | 15 | 457 | 5.12991E-73 | 481 | 2 | 12 | GFS | W | Abs. | FPAVTIC | A | GLCMTFN | 16 | 395 | 19 |
| LDEC019897 | *Ldec019897* | V | 29 | 429 | 1.59465E-73 | 450 | 2 | 11 | GFS | W | TGIHG | FPAVTIC | S | GICYSYN | 50 | 333 | 16 |
| LDEC024423 | *Ldec024423* | V | 6 | 409 | 2.88717E-40 | 410 | 1 | 12 | GFS | - | Abs. | MAMVC | A | GICYSFN | 29 | 348 | - |
| LDEC019444 | *LdecPPK3* | II | 23 | 113 | 0.000292006 | 174 | 0 | 5 | GIS | - | Abs. | Abs. | V | Abs. | 5 | 105 | 29 |
| LDEC010412 | *LdecPPK9* | II | 1 | 360 | 3.92894E-33 | 382 | 2 | 11 | GCS | W | Abs. | Abs. | V | GQCYSFN | 14 | 300 | 17 |
| JAMg_model_10489.2 | *Lmig10489.2* | IV | 22 | 465 | 3.8345E-93 | 528 | 2 | 14 | GCS | W | TTLHG | YPAVTVC | A | GFCCSFN | 44 | 390 | 43 |
| JAMg_model_11504.1 | *Lmig11504.1* | I | 1 | 147 | 6.60498E-12 | 180 | 1 | 6 | Abs. | W | Abs. | FPAVSIC | - | RPSACKR | 1 | 154 | - |
| JAMg_model_12479.1 | *Lmig12479.1* | III | 56 | 301 | 3.64077E-27 | 351 | 2 | 10 | GVS | - | Abs. | Abs. | V | Abs. | 1 | 276 | 45 |
| JAMg_model_12972.1 | *Lmig12972.1* | IV | 25 | 191 | 4.9574E-17 | 193 | 0 | 11 | Abs. | - | KLF | Abs. | N | Abs. | 2 | 188 | 2 |
| JAMg_model_14370.1 | *Lmig14370* | I | 2 | 358 | 3.41012E-38 | 407 | 2 | 13 | GAS | - | Abs. | Abs. | V | GVCYAIN | 8 | 320 | 44 |
| JAMg_model_14820.1 | *Lmig14820.1* | V | 45 | 387 | 5.21275E-64 | 465 | 2 | 12 | GFS | - | Abs. | VC | A | GLCFTFN | 25 | 341 | 73 |
| JAMg_model_1583.1 | *Lmig1583.1* | III | 4 | 132 | 4.03543E-11 | 345 | 0 | 5 | Abs. | - | Abs. | FPAVTVC | S | GFCYSFN | 1 | 319 | 14 |
| JAMg_model_17317.1 | *Lmig17317.1* | V | 47 | 513 | 3.02253E-76 | 523 | 2 | 14 | GFS | W | TSLHG | FPAVTLC | A | GICYSFN | 69 | 398 | 6 |
| JAMg_model_18227.1 | *Lmig18227.1* | IV | 46 | 173 | 1.6019E-16 | 233 | 2 | 5 | GFS | - | Abs. | Abs. | S | Abs. | 1 | 151 | 55 |
| JAMg_model_18332.1 | *Lmig18332* | IV | 39 | 281 | 6.21847E-30 | 311 | 1 | 4 | Abs. | W | TAQHG | FPAITIC | - | GRCCAFN | 61 | 216 | 8 |
| JAMg_model_2248.1 | *Lmig2248.1* | V | 2 | 396 | 1.03841E-45 | 602 | 2 | 8 | GFS | - | Abs. | FPAVTIC | A | GLCYSFN | 1 | 355 | 200 |
| JAMg_model_8491.1 | *Lmig8491.1* | V | 65 | 246 | 8.74798E-30 | 258 | 1 | 3 | Abs. | W | TSIHG | FPAVTIC | - | GVCFTFN | 87 | 145 | - |
| JAMg_model_9685.1 | *Lmig9685.1* | V | 89 | 345 | 1.36881E-58 | 367 | 1 | 10 | GFS | - | Abs. | FN | A | Abs. | 11 | 313 | 17 |
| JAMg_model_9564.3 | *LmigPPK13* | I | 4 | 366 | 1.25548E-34 | 383 | 2 | 14 | GAS | - | Abs. | FPTIVLC | V | GLCFAIN | 1 | 345 | 11 |
| JAMg_model_7461.1 | *LmigPPK16* | IV | 12 | 374 | 5.72831E-47 | 404 | 2 | 13 | GFS | - | Abs. | AQSIL | G | GYCCSFN | 7 | 347 | 24 |
| JAMg_model_492.1 | *LmigPPK17* | V | 11 | 336 | 5.10821E-10 | 383 | 1 | 10 | GLS | - | Abs. | YPAVTIC | A | GRCYTLK | 1 | 309 | 36 |

| | | | | | | | | | | | | | | | | | |
|---|---|---|---|---|---|---|---|---|---|---|---|---|---|---|---|---|---|
| | | II | | | | | | | | | | | | | | | |
| JAMg_model_1998.1 | *LmigPPK28a* | V | 1 | 200 | 6.7853E-30 | 202 | 0 | 8 | Abs. | - | Abs. | Abs. | - | GICCSFN | 1 | 199 | 1 |
| JAMg_model_20185.1 | *LmigPPK28b* | V | 21 | 122 | 6.35747E-17 | 165 | 1 | 3 | GFS | - | Abs. | Abs. | S | Abs. | 2 | 99 | 38 |
| JAMg_model_1169.1 | *Lmig1169.1* | II | 19 | 421 | 8.8226E-39 | 422 | 1 | 12 | Abs. | W | TSVHG | FPAVTIC | I | GLCDTLA | 39 | 357 | - |
| XP_019891600.1 | *Mdom019891600.1* | II | 111 | 328 | 9.24147E-15 | 347 | 2 | 11 | GIS | W | NPMHG | YPSVTLC | V | Abs. | 42 | 254 | 14 |
| XP_019894840.1 | *Mdom019894840.1* | III | 3 | 137 | 0.0440384 | 182 | 2 | 3 | Abs. | - | ALR | ILC | - | Abs. | 1 | 113 | 39 |
| XP_005174914.1 | *MdomPPK1* | V | 65 | 572 | 1.97061E-85 | 601 | 2 | 14 | GIS | W | TSVHG | FPTITIC | A | GICYQFN | 87 | 439 | 53 |
| XP_005175357.1 | *MdomPPK10a* | II | 12 | 361 | 4.38104E-30 | 375 | 1 | 11 | GFS | - | Abs. | Abs. | V | GLCYLTN | 6 | 334 | 24 |
| XP_005187879.1 | *MdomPPK10b* | II | 19 | 473 | 1.73218E-58 | 490 | 2 | 12 | GFS | W | SNIHG | MPSVTIC | V | GLCYMTN | 41 | 386 | 9 |
| XP_019895075.1 | *MdomPPK11* | IV | 155 | 615 | 2.00837E-67 | 673 | 4 | 14 | GYS | W | ASIHG | FPAVTIC | G | GICCAFN | 175 | 394 | 12 |
| XP_019890716.1 | *MdomPPK12a* | V | 27 | 492 | 1.99104E-78 | 574 | 2 | 14 | GFS | F | TTLHG | LPALTLC | S | GLCCSFN | 49 | 397 | 77 |
| XP_019890715.1 | *MdomPPK12b* | V | 30 | 504 | 4.07919E-79 | 572 | 2 | 14 | GFS | F | TTLHG | LPALTLC | S | GLCCAFN | 52 | 406 | 63 |
| XP_005184244.2 | *MdomPPK13* | I | 11 | 432 | 3.78572E-68 | 454 | 2 | 14 | GAS | W | STLHG | FPAIVVC | V | GLCYAIN | 33 | 353 | 17 |
| XP_019891003.1 | *MdomPPK14a* | III | 15 | 168 | 1.40478E-23 | 215 | 1 | 9 | GCS | - | Abs. | Abs. | A | Abs. | - | 147 | 42 |
| XP_005190919.1 | *MdomPPK14b* | III | 36 | 322 | 1.21541E-25 | 323 | 1 | 5 | Abs. | L | TTVHG | FPEVTVC | - | GACLAFN | 54 | 243 | - |
| XP_019894349.1 | *MdomPPK15* | I | 38 | 251 | 2.61452E-31 | 269 | 2 | 10 | GAS | - | Abs. | Abs. | V | Abs. | 36 | 194 | 13 |
| XP_011292215.1 | *Mdom011292215.1* | I | 1 | 296 | 4.9093E-11 | 314 | 0 | 9 | Abs. | - | G | YPSVAVC | A | GKCFILN | 3 | 285 | 4 |
| XP_019895076.1 | *MdomPPK16* | IV | 91 | 555 | 3.12097E-75 | 569 | 2 | 14 | GFS | W | TSLHG | FPAVTLC | A | GECCTFN | 113 | 396 | 9 |
| XP_005176331.1 | *MdomPPK17* | VII | 22 | 363 | 1.09448E-07 | 432 | 2 | 10 | GLS | F | P | MPAVTIC | A | GRCHTIR | 8 | 321 | 40 |
| XP_019895070.1 | *MdomPPK18* | IV | 3 | 382 | 9.28623E-31 | 447 | 1 | 11 | GFS | - | Abs. | Abs. | N | GVCCMFN | 1 | 360 | 60 |
| XP_019894439.1 | *MdomPPK19a* | III | 10 | 58 | 0.872187 | 284 | 0 | 5 | Abs. | Y | Abs. | LAVC | - | GICLVFN | 27 | 237 | - |
| XP_019893823.1 | *MdomPPK19b* | III | 11 | 249 | 1.32874E-21 | 250 | 0 | 5 | Abs. | - | Abs. | FPALALC | - | GVCLVFN | - | 240 | - |
| XP_019893824.1 | *MdomPPK19c* | III | 79 | 505 | 8.58112E-49 | 537 | 2 | 14 | GAS | W | ECIAG | FPALALC | G | GVCLVFN | 70 | 390 | 26 |
| XP_011294238.1 | *MdomPPK19d* | III | 51 | 505 | 1.23527E-63 | 544 | 2 | 14 | GAS | W | NSING | FPAVAVC | S | GICLVFN | 72 | 388 | 33 |
| XP_005187753.2 | *MdomPPK19e* | III | 48 | 459 | 1.55572E-54 | 499 | 3 | 14 | GAS | W | NTIHG | FPAVAVC | S | GVCLVFN | 69 | 345 | 34 |
| XP_019893825.1 | *MdomPPK19f* | III | 52 | 298 | 3.58928E-24 | 310 | 1 | 5 | Abs. | W | HTIHG | FPAVAVC | S | GECMVFN | 73 | 207 | 4 |
| XP_019894440.1 | *MdomPPK20* | III | 51 | 512 | 3.59e-80 | 577 | 2 | 14 | GAS | W | STIHG | FPAVGVC | A | GYCFVFN | 73 | 393 | 60 |
| XP_019894437.1 | *MdomPPK21a* | III | 13 | 311 | 3.21116E-24 | 314 | 1 | 8 | Abs. | W | Abs. | FPSIGLC | - | GYCYTFN | - | 287 | 1 |
| XP_019894438.1 | *MdomPPK21b* | III | 1 | 18 | 0.00506451 | 528 | 2 | 14 | GAS | W | SAVEL | FPSIGLC | A | GFCFIFN | 38 | 395 | 44 |
| XP_019893922.1 | *MdomPPK22* | IV | 39 | 490 | 1.40375E-58 | 493 | 1 | 14 | Abs. | W | ISLHA | FPAVSIC | R | GPCCMFN | 60 | 407 | - |
| XP_005180321.1 | *MdomPPK23* | VI | 30 | 447 | 2.01839E-72 | 563 | 2 | 14 | GFS | W | STLHG | FPTTVVC | V | GYCFAFN | 52 | 349 | 111 |
| XP_019893923.1 | *MdomPPK24* | IV | 27 | 268 | 1.39214E-21 | 277 | 0 | 8 | Abs. | - | Abs. | FPAVSIC | - | DELFFNT | 1 | 265 | - |
| XP_019891859.1 | *MdomPPK25* | II | 14 | 436 | 6.25557E-37 | 484 | 3 | 12 | GAS | W | SSIHG | FPFVGITLC | M | GLCRSTS | 32 | 359 | 42 |
| XP_011291290.1 | *MdomPPK26* | V | 66 | 559 | 5.7241E-113 | 589 | 2 | 14 | GVS | W | STIHG | FPAITIC | A | GPCYTFN | 88 | 425 | 25 |

| | | | | | | | | | | | | | | | | | |
|---|---|---|---|---|---|---|---|---|---|---|---|---|---|---|---|---|---|
| XP_005178987.1 | *MdomPPK27a* | IV | 26 | 492 | 8.55505E-64 | 509 | 2 | 14 | GCS | W | TSLNG | FPSIAVC | S | GYCCVFN | 48 | 398 | 12 |
| XP_019890758.1 | *MdomPPK27b* | IV | 12 | 463 | 2.88212E-58 | 480 | 2 | 14 | GCS | W | Abs. | FPSIAVC | S | GYCCVFN | 12 | 405 | 12 |
| XP_005189738.2 | *MdomPPK28a* | V | 106 | 364 | 6.53134E-54 | 446 | 1 | 10 | GFS | - | TTLHG | Abs. | S | Abs. | 75 | 268 | 77 |
| XP_005189737.1 | *MdomPPK28b* | V | 36 | 294 | 5.56888E-54 | 376 | 1 | 10 | GFS | - | Abs. | Abs. | S | Abs. | 5 | 268 | 77 |
| XP_005181951.1 | *MdomPPK29* | I | 13 | 436 | 1.42884E-27 | 443 | 2 | 11 | GAS | W | Abs. | FPAVSFC | V | GYCYLAN | 21 | 370 | 1 |
| XP_019890548.1 | *MdomPPK2a* | V | 51 | 553 | 3.0119E-104 | 567 | 2 | 14 | GFS | W | TSIHG | FPAVTIC | A | GICYTFN | 72 | 436 | 8 |
| XP_011292906.2 | *MdomPPK2b* | V | 16 | 214 | 1.69891E-42 | 214 | 0 | 11 | Abs. | - | Abs. | Abs. | A | Abs. | 9 | 190 | - |
| XP_019890549.1 | *MdomPPK2c* | V | 66 | 560 | 2.9027E-90 | 574 | 2 | 14 | GVS | W | TSIHG | FPAITLC | A | GLCFTFN | 87 | 428 | 8 |
| XP_019890552.1 | *MdomPPK2d* | V | 31 | 516 | 1.27261E-85 | 531 | 2 | 14 | GVS | W | TSIHG | FPAVTIC | A | GICYTFN | 52 | 419 | 9 |
| XP_005178343.1 | *MdomPPK2e* | V | 46 | 533 | 1.65524E-92 | 547 | 2 | 14 | GFS | W | TSVHG | FPAITIC | A | GLCYTFN | 67 | 421 | 8 |
| XP_011294524.1 | *MdomPPK2f* | V | 47 | 534 | 2.38846E-93 | 561 | 2 | 14 | GFS | W | TSLHG | FPAITIC | A | GLCYTFN | 68 | 421 | 21 |
| XP_011290422.1 | *MdomPPK2g* | V | 45 | 531 | 5.75704E-71 | 566 | 2 | 12 | GFS | W | TALHG | FPAVTIC | S | GVCYTFN | 66 | 440 | 9 |
| XP_005176880.1 | *MdomPPK2h* | V | 37 | 536 | 2.12587E-95 | 553 | 2 | 14 | GVS | W | SSIHG | FPAVTIC | A | GICFTFN | 58 | 433 | 11 |
| XP_005176877.1 | *MdomPPK2i* | V | 46 | 537 | 2.19416E-96 | 554 | 2 | 14 | GIS | W | TSIHG | FPAITIC | A | GICYTFN | 67 | 425 | 11 |
| XP_005176878.1 | *MdomPPK2j* | V | 47 | 543 | 2.13496E-84 | 561 | 2 | 14 | GVS | W | TSIHG | FPRLTIC | A | GICYAFN | 68 | 430 | 12 |
| XP_005176879.1 | *MdomPPK2k* | V | 46 | 543 | 7.75734E-86 | 561 | 2 | 14 | GVS | W | TSIHG | FPRLTIC | A | GICYAFN | 67 | 431 | 12 |
| XP_005179130.2 | *MdomPPK3* | II | 62 | 467 | 2.04683E-48 | 568 | 4 | 12 | GIS | W | TGVLA | MPAITLC | V | GSCYVIN | 71 | 351 | 9 |
| XP_019894441.1 | *MdomPPK30* | III | 3 | 456 | 2.7793E-49 | 496 | 2 | 14 | GFS | - | Abs. | FPAVTIC | V | GICYSFN | 4 | 413 | 95 |
| XP_005178919.2 | *MdomPPK31* | I | 14 | 405 | 1.67437E-39 | 419 | 2 | 14 | NVS | Y | Abs. | FPAVTIC | I | GLCFAFN | 12 | 346 | 33 |
| XP_005178822.1 | *MdomPPK4* | IV | 29 | 468 | 8.8532E-83 | 507 | 2 | 14 | GMS | W | TSIHG | FPATTIC | S | GFCCAFN | 51 | 371 | 34 |
| XP_005179825.2 | *MdomPPK5a* | V | 7 | 411 | 4.9781E-72 | 446 | 1 | 14 | GFS | - | Abs. | FPSVTIC | S | GICCSFN | 1 | 389 | 30 |
| XP_019891014.1 | *MdomPPK5b* | V | 33 | 501 | 2.34631E-76 | 575 | 2 | 14 | GFS | Y | TALNG | FPTVTIC | S | GLCCVFN | 55 | 400 | 69 |
| XP_011290988.2 | *MdomPPK5c* | V | 1 | 332 | 1.27312E-51 | 401 | 1 | 13 | GFS | - | Abs. | Abs. | S | GMCCVFN | - | 311 | 64 |
| XP_005175804.1 | *MdomPPK6* | IV | 30 | 465 | 2.8692E-54 | 483 | 2 | 14 | GLS | W | TKVSG | FPGVTVC | S | GPCCSFN | 52 | 367 | 13 |
| XP_005190920.2 | *MdomPPK7* | III | 55 | 482 | 1.07663E-58 | 487 | 1 | 14 | Abs. | W | TTVHG | FPTITIC | - | GICWSFN | 77 | 380 | 3 |
| XP_019894556.1 | *MdomPPK8a* | V | 60 | 103 | 0.00109814 | 695 | 3 | 14 | GFS | L | TTLHG | FPAVTIC | S | GLCCSFN | 86 | 392 | 80 |
| XP_019894557.1 | *MdomPPK8b* | V | 38 | 507 | 4.42435E-86 | 601 | 2 | 14 | GFS | F | TTLHG | FPALTIC | S | GHCCAFN | 60 | 401 | 89 |
| XP_019892519.1 | *MdomPPK8c* | V | 42 | 506 | 1.67214E-91 | 604 | 2 | 14 | GFS | F | STLHG | FPAITFC | S | GLCCVFN | 64 | 396 | 93 |
| XP_011293257.1 | *MdomPPK9* | II | 213 | 638 | 5.61125E-53 | 649 | 3 | 12 | GCS | W | SSIHG | FPSLTVC | V | GTCHSVN | 235 | 357 | 6 |
| MYZPE13164_G006_v1.0_000000820.1_pep | *Mper000000820.1* | V | 10 | 461 | 8.36279E-26 | 1762 | 4 | 9 | SCA | W | KLTNI | FPSIMIC | A | GLCYTIN | 472 | 1062 | 180 |
| MYZPE13164_G006_v1.0_000036460.1_pep | *Mper000036460.1* | V | 3 | 445 | 5.27747E-55 | 472 | 1 | 13 | GCS | - | Abs. | FPAITFC | G | GLCYSIN | 1 | 405 | 22 |
| MYZPE13164_G006_v1.0_000070390.1_pep | *Mper000070390.1* | I | 7 | 430 | 1.3409E-61 | 466 | 2 | 14 | GVS | W | STIHG | VPAVTVC | V | GVCFIVN | 29 | 355 | 31 |
| MYZPE13164_G006_v1.0_000084580.1_pep | *Mper000084580.1* | I | 46 | 273 | 8.08912E-07 | 340 | 2 | 10 | GMN | W | Abs. | FPAVTIC | - | NPCFCIN | 12 | 285 | 5 |
| MYZPE13164_G006_v1.0_000086800.1_pep | *Mper000086800.1* | III | 10 | 478 | 1.51931E-76 | 502 | 2 | 14 | GCS | W | GSIHG | FPGITIC | V | GFCYSFN | 32 | 401 | 18 |
| MYZPE13164_G006_v | *Mper000095070.1* | V | 39 | 514 | 1.69977E-51 | 541 | 2 | 13 | GAS | W | TSLHG | FPMVVIC | G | GVCYSLN | 61 | 407 | 22 |

| | | | | | | | | | | | | | | | | | |
|---|---|---|---|---|---|---|---|---|---|---|---|---|---|---|---|---|---|
| 1.0_000095070.1_pep | | | | | | | | | | | | | | | | | |
| MYZPE13164_G006_v 1.0_000138700.1_pep | *Mper000138700.1* | III | 43 | 512 | 1.45727E-51 | 1135 | 4 | 14 | GGS | W | CSLHG | FPSVTIC | V | GLCHSFN | 630 | 402 | 52 |
| MYZPE13164_G006_v 1.0_000138710.1_pep | *Mper000138710.1* | III | 23 | 436 | 1.56613E-36 | 460 | 2 | 11 | GAS | W | ISVNA | FPAVNVC | V | Abs. | 39 | 351 | 19 |
| MYZPE13164_G006_v 1.0_000154690.1_pep | *Mper000154690.1* | III | 39 | 514 | 1.39145E-72 | 539 | 2 | 14 | GFS | W | CTVHG | FPSITIC | V | GFCYSFN | 61 | 408 | 19 |
| MYZPE13164_G006_v 1.0_000154700.1_pep | *Mper000154700.1* | III | 49 | 520 | 2.56305E-71 | 535 | 2 | 14 | GFS | W | STVHG | FPAVSVC | V | GFCYSFN | 69 | 406 | 9 |
| MYZPE13164_G006_v 1.0_000163900.1_pep | *Mper000163900.1* | I | 5 | 422 | 3.78315E-62 | 448 | 2 | 14 | GAS | W | STIHG | FPAVTIC | V | GICYTLT | 27 | 349 | 21 |
| MYZPE13164_G006_v 1.0_000164680.1_pep | *Mper000164680.1* | III | 10 | 478 | 6.08347E-72 | 1020 | 4 | 14 | GCS | W | GSIHG | FPAITIC | V | GFCYSFN | 32 | 401 | 536 |
| MYZPE13164_G006_v 1.0_000165800.1_pep | *Mper000165800.1* | V | 19 | 481 | 2.61582E-33 | 489 | 3 | 12 | GVS | W | Abs. | FPAVTVC | - | DVLQTGK | 36 | 399 | 3 |
| MYZPE13164_G006_v 1.0_000191120.1_pep | *Mper000191120.1* | V | 61 | 515 | 8.01704E-76 | 545 | 2 | 13 | GFS | W | TSLHG | FLSLSIC | G | GLCYSFN | 83 | 386 | 25 |
| MYZPE13164_G006_v 1.0_000066440.1_pep | *MperPPK23a* | VI | 30 | 216 | 1.86368E-33 | 268 | 1 | 5 | Abs. | W | STLHG | FPAITVC | - | GFCYTFN | 52 | 187 | 3 |
| MYZPE13164_G006_v 1.0_000035270.1_pep | *MperPPK23b* | VI | 30 | 450 | 2.85727E-79 | 516 | 2 | 14 | GFS | W | STLHG | FPAITVC | V | GFCYTFN | 52 | 352 | 61 |
| MYZPE13164_G006_v 1.0_000060760.1_pep | *MperPPK28* | V | 59 | 470 | 1.34367E-70 | 486 | 1 | 14 | Abs. | W | TTLHG | FPAITIC | - | GICCSFN | 81 | 379 | - |
| MYZPE13164_G006_v 1.0_000070370.1_pep | *MperPPK9a* | II | 44 | 467 | 2.5682E-57 | 480 | 2 | 12 | GCS | W | SNIHG | LPAVTLC | V | GLCYSYN | 66 | 355 | 8 |
| MYZPE13164_G006_v 1.0_000070380.1_pep | *MperPPK9b* | II | 44 | 449 | 5.33765E-48 | 462 | 2 | 10 | GCS | W | SNIHG | LPAFTLC | V | GLCYSYN | 66 | 337 | 8 |
| PHUM527890-PA | *Phumppk17* | VII | 27 | 376 | 2.19432E-14 | 389 | 2 | 9 | GLS | G | R | YPAVTIC | G | GMCHTIK | 17 | 306 | 12 |
| g18724.t1 | *Pxyl18724.t1* | IV | 67 | 123 | 0.0946322 | 250 | 0 | 8 | Abs. | - | Abs. | Abs. | P | Abs. | 1 | 223 | 5 |
| g16424.t1 | *PxylPPK13a* | I | 115 | 694 | 5.1846E-36 | 718 | 3 | 14 | GAS | W | CSFTG | MPAVALC | V | GTCYIFN | 137 | 511 | 19 |
| g16431.t1 | *PxylPPK13b* | I | 23 | 438 | 4.17794E-59 | 496 | 2 | 14 | GAS | W | TSIHG | FPSVAVC | V | GTCYAIN | 43 | 350 | 52 |
| g28744.t1 | *PxylPPK16a* | IV | 131 | 458 | 5.55464E-47 | 510 | 1 | 11 | GFT | R | Abs. | Abs. | A | GLCCTFN | 28 | 387 | 47 |
| g28745.t1 | *PxylPPK16b* | IV | 30 | 121 | 3.59473E-15 | 220 | 1 | 2 | Abs. | W | TDLHG | FPAVTIC | - | Abs. | 52 | 142 | - |
| g893.t1 | *PxylPPK17* | VII | 29 | 413 | 1.05742E-08 | 473 | 2 | 8 | GLS | L | PLT | YPSLTFC | S | GRCYTLE | 16 | 344 | 59 |
| g18344.t1 | *PxylPPK23a* | VI | 40 | 392 | 3.26114E-49 | 410 | 1 | 10 | Abs. | W | STLHG | FPAVTIC | - | GFCYAFN | 62 | 314 | - |
| g28101.t1 | *PxylPPK25* | II | 18 | 272 | 9.78191E-22 | 874 | 3 | 12 | GAS | W | TSMHG | FPAITAC | V | GACYVTN | 387 | 373 | 63 |
| g35214.t1 | *PxylPPK28a* | V | 6 | 251 | 2.94237E-56 | 272 | 1 | 10 | GFS | - | Abs. | Abs. | S | Abs. | 1 | 229 | 16 |
| g35217.t1 | *PxylPPK28b* | V | 29 | 198 | 5.14905E-33 | 252 | 1 | 4 | Abs. | W | STLHG | FPAVTIC | - | GLCCTFN | 51 | 174 | 1 |
| g15752.t1 | *Pxyl15752.t1* | II | 101 | 272 | 4.10814E-09 | 273 | 0 | 8 | Abs. | - | DIH | PSISL | - | GICMTFN | 1 | 268 | - |
| g30278.t1 | *Pxyl30278.t1* | II | 58 | 385 | 1.00455E-33 | 413 | 1 | 12 | Abs. | W | TSIHG | FPAVTAC | R | GICMTFN | 80 | 292 | - |
| g26486.t1 | *PxylPPK6a* | IV | 10 | 260 | 3.6298E-21 | 283 | 1 | 10 | GCS | - | S | FPT | A | GTLLN | 1 | 233 | 17 |
| g26485.t1 | *PxylPPK6b* | IV | 10 | 197 | 1.36666E-24 | 200 | 1 | 4 | Abs. | W | TTIIG | WPAITIC | - | GACCIFN | 31 | 143 | - |
| g18897.t1 | *PxylPPK9a* | II | 234 | 531 | 1.80885E-37 | 543 | 4 | 7 | GCS | W | ASFHG | FPAATVC | V | GLCYTIN | 77 | 408 | 7 |
| g31367.t1 | *PxylPPK9b* | II | 55 | 223 | 1.87665E-14 | 232 | 1 | 2 | Abs. | W | ASFHG | FPAATVC | - | GLCYTIN | 77 | 129 | - |
| g10292.t1 | *Pxyl10292.t1* | V | 21 | 210 | 9.96835E-22 | 244 | 1 | 8 | GFS | - | Abs. | Abs. | A | Abs. | 1 | 203 | 14 |

| | | | | | | | | | | | | | | | | | |
|---|---|---|---|---|---|---|---|---|---|---|---|---|---|---|---|---|---|
| g10294.t1 | *Pxyl10294.t1* | V | 42 | 222 | 2.09801E-19 | 263 | 1 | 1 | GLP | W | TTLHG | FPSITIC | K | FTLD | 62 | 175 | - |
| g10539.t1 | *Pxyl10539.t1* | IV | 40 | 403 | 2.53522E-23 | 420 | 2 | 11 | GFS | W | CTLAG | FPAVAIC | S | DDKFNIS | 62 | 296 | 12 |
| g11832.t1 | *Pxyl11832.t1* | I | 30 | 421 | 1.23295E-35 | 585 | 2 | 14 | Abs. | Y | LTVHG | LPSTTVC | V | GICFSSN | 51 | 344 | 80 |
| g12604.t1 | *Pxyl12604.t1* | V | 79 | 572 | 2.82453E-37 | 596 | 4 | 8 | GFS | W | GSLHG | FPSVVLC | - | GLCYAFN | 101 | 425 | 19 |
| g15324.t1 | *Pxyl15324.t1* | I | 26 | 283 | 9.78988E-24 | 295 | 0 | 13 | Abs. | - | Abs. | Abs. | - | GICFSSN | - | 291 | - |
| g16919.t1 | *Pxyl16919.t1* | IV | 13 | 407 | 3.92024E-30 | 442 | 2 | 13 | GVG | W | TTLHG | MPDVGVC | S | GVCCIVR | 33 | 328 | 30 |
| g18725.t1 | *Pxyl18725.t1* | IV | 1 | 237 | 5.11179E-17 | 285 | 1 | 5 | Abs. | W | Abs. | FPAVAFC | - | GFCCIFN | 1 | 248 | 6 |
| g20100.t1 | *Pxyl20100.t1* | I | 24 | 407 | 2.151E-35 | 424 | 1 | 14 | Abs. | W | SCLHG | FPAVTVC | - | GTCFAFN | 47 | 338 | 1 |
| g21610.t1 | *Pxyl21610.t1* | I | 24 | 410 | 1.63095E-32 | 419 | 2 | 14 | GKS | W | SCLHG | FPAVTVC | - | GTCFAFN | 47 | 333 | 1 |
| g26144.t2 | *Pxyl26144.t2* | IV | 13 | 231 | 2.81421E-12 | 247 | 1 | 3 | GVT | W | AQFHG | Abs. | - | GICCIFN | 27 | 194 | - |
| g26358.t1 | *Pxyl26358.t1* | V | 93 | 344 | 2.8055E-25 | 368 | 2 | 8 | GFS | - | Abs. | Abs. | - | Abs. | 1 | 322 | 19 |
| g34278.t1 | *Pxyl34278.t1* | IV | 1 | 412 | 8.76773E-30 | 540 | 2 | 10 | GGS | W | MHG | FPGLVLC | A | GYCCQFH | 16 | 362 | 111 |
| g34279.t1 | *Pxyl34279.t1* | IV | 10 | 234 | 5.10322E-27 | 284 | 0 | 8 | Abs. | - | Abs. | Abs. | T | GYCCAFN | 1 | 281 | 1 |
| g36719.t1 | *Pxyl36719.t1* | V | 50 | 231 | 5.68843E-31 | 243 | 1 | 4 | Abs. | W | SNLHG | YPAVTVC | - | GQCYTFN | 72 | 145 | - |
| g36720.t1 | *Pxyl36720.t1* | V | 6 | 355 | 5.42354E-70 | 382 | 2 | 13 | GFS | - | Abs. | Abs. | A | GQCYTFN | 6 | 328 | 22 |
| g36750.t1 | *Pxyl36750.t1* | I | 5 | 265 | 6.19844E-38 | 273 | 1 | 12 | GGS | - | Abs. | Abs. | V | GTCYTFN | 4 | 240 | 3 |
| g5537.t1 | *Pxyl5537.t1* | V | 5 | 217 | 7.95955E-38 | 238 | 0 | 10 | Abs. | - | Abs. | Abs. | A | Abs. | 1 | 214 | 5 |
| g5538.t1 | *Pxyl5538.t1* | V | 1 | 157 | 1.22831E-11 | 176 | 2 | 8 | GFS | - | Abs. | Abs. | A | Abs. | 1 | 135 | 14 |
| g25324.t1 | *PxylPPK23b* | VI | 40 | 622 | 4.362E-60 | 704 | 2 | 14 | GFS | W | STLHG | FPAVTIC | V | GFCYAFN | 62 | 514 | 77 |
| RPRC000048 | *Rpro000048* | V | 24 | 499 | 1.6041E-111 | 543 | 2 | 14 | GVS | W | TSIHG | FPAVTLC | A | GVCFSFN | 46 | 407 | 39 |
| RPRC000341 | *Rpro000341* | I | 18 | 439 | 1.36436E-70 | 501 | 2 | 14 | GAS | W | TSFHG | FPSISVC | V | GPCYMFS | 40 | 353 | 57 |
| RPRC013510 | *Rpro013510* | V | 55 | 528 | 2.15296E-47 | 540 | 2 | 13 | GFS | W | TTLHG | FPAITLC | A | GACLTFN | 77 | 405 | 7 |
| RPRC014276 | *Rpro014276* | III | 19 | 496 | 9.81857E-52 | 523 | 2 | 14 | GGS | W | STFHC | FPAVTIC | V | GFCYSFN | 41 | 410 | 21 |
| RPRC000099 | *RproPPK23* | VI | 58 | 479 | 4.13969E-82 | 552 | 2 | 14 | GFS | W | STLHG | FPGVTVC | V | GFCYSFN | 80 | 353 | 68 |
| RPRC000471 | *RproPPK28* | V | 38 | 497 | 9.69331E-85 | 563 | 2 | 14 | GFS | W | SSMHG | FPAITVC | S | GICCVFN | 60 | 413 | 39 |
| Abs. in VectorBase | *RproPPKlike7* | II | 19 | 444 | 3.86206E-61 | 459 | 2 | 12 | GCS | W | TTLHG | FPAATAC | V | GICYTFA | 41 | 357 | 10 |
| Abs. in VectorBase | *RproPPK9* | II | 28 | 425 | 2.72465E-40 | 434 | 1 | 12 | GME | W | SSIHG | LPAVSIC | - | GICYSYN | 50 | 349 | 5 |
| RproPPKlike5 | *RproPPKlike5* | I | 1 | 272 | 2.37947E-46 | 337 | 1 | 12 | GAS | - | Abs. | Abs. | V | GVCYSLN | 1 | 250 | 60 |
| RproPPKlike6 | *RproPPKlike6* | I | 14 | 301 | 1.07971E-15 | 389 | 2 | 12 | KQS | V | Abs. | LPALSVC | - | TKDGVCY | 16 | 315 | 25 |
| SFRICE002083 | *Sfru002083* | IV | 41 | 433 | 2.47687E-45 | 542 | 1 | 14 | Abs. | W | SSICG | FPAVGIC | V | GYCCTFN | 63 | 370 | 16 |
| SFRICE009385 | *Sfru009385* | I | 27 | 258 | 1.07276E-05 | 272 | 1 | 7 | Abs. | W | SSGVHG | FPGVTVC | - | Abs. | 49 | 196 | - |
| SFRICE017730 | *Sfru017730* | IV | 119 | 464 | 9.59459E-26 | 478 | 1 | 9 | Abs. | W | Abs. | FPSVIVC | - | GFCCVFN | 117 | 335 | - |
| SFRICE019222 | *Sfru019222* | IV | 1 | 235 | 5.11487E-22 | 345 | 0 | 10 | Abs. | W | Abs. | Abs. | - | ILVFNSY | 1 | 343 | - |

| | | | | | | | | | | | | | | | | | |
|---|---|---|---|---|---|---|---|---|---|---|---|---|---|---|---|---|---|
| SFRICE020430 | *Sfru020430* | IV | 313 | 548 | 0.000357397 | 682 | 2 | 9 | NKN | W | VNTHK | FPAVAIC | - | Abs. | 67 | 498 | 28 |
| SFRICE023811 | *Sfru023811* | V | 41 | 234 | 4.22123E-30 | 437 | 1 | 5 | Abs. | W | TTLHG | FPSVTIC | S | GICYTMN | 63 | 320 | 25 |
| SFRICE024850 | *Sfru024850* | V | 68 | 510 | 1.00998E-70 | 510 | 1 | 14 | Abs. | W | SNLHG | YPAVTVC | - | GLCYTFN | 90 | 305 | 1 |
| SFRICE030732 | *Sfru030732* | I | 24 | 440 | 9.1599E-57 | 449 | 3 | 14 | GGS | Y | LTVHG | FASVTVC | V | GTCYSIN | 45 | 393 | 4 |
| SFRICE032362 | *Sfru032362* | IV | 10 | 307 | 1.09726E-32 | 307 | 0 | 13 | Abs. | - | Abs. | Abs. | - | GHCCTFN | 1 | 349 | 1 |
| SFRICE034560 | *Sfru034560* | V | 1 | 138 | 4.05867E-11 | 185 | 2 | 8 | GFS | - | Abs. | Abs. | S | Abs. | 1 | 265 | 27 |
| SFRICE035107 | *Sfru035107* | IV | 1 | 287 | 6.66296E-08 | 292 | 1 | 10 | Abs. | W | Abs. | VPTVAVC | - | GVCCVMR | 1 | 296 | - |
| SFRICE023370 | *Sfru023370* | II | 11 | 332 | 7.23195E-12 | 332 | 0 | 12 | RLS | - | P | YPTIVLC | P | GMCHIIN | 9 | 131 | - |
| SFRICE002084 | *SfruPPK13a* | I | 163 | 465 | 4.74466E-36 | 468 | 1 | 9 | Abs. | W | CTFAG | MPAVAIC | - | GPCFILN | 185 | 266 | - |
| SFRICE008547 | *SfruPPK13b* | I | 24 | 133 | 2.27318E-16 | 202 | 2 | 5 | GAS | - | Abs. | Abs. | V | Abs. | 1 | 257 | 64 |
| SFRICE008543 | *SfruPPK13c* | I | 23 | 438 | 4.9981E-60 | 496 | 2 | 14 | GAS | W | TSIHG | FPTVVVC | V | GTCYAIN | 43 | 111 | 52 |
| SFRICE026402 | *SfruPPK17* | VII | 64 | 425 | 6.31618E-09 | 427 | 1 | 8 | Abs. | L | PRWS | YPSITLC | S | GRCYTMS | 50 | 350 | - |
| SFRICE002679 | *SfruPPK23* | VI | 40 | 469 | 2.13863E-73 | 543 | 2 | 14 | GFS | W | STLHG | FPAVTIC | V | GFCYAFN | 62 | 339 | 69 |
| SFRICE018167 | *SfruPPK25a* | II | 35 | 314 | 1.66873E-13 | 349 | 1 | 3 | Abs. | W | TSMHG | LPALTLC | - | GVCHTTN | 57 | 361 | 3 |
| SFRICE030048 | *SfruPPK25b* | II | 65 | 237 | 2.88336E-12 | 369 | 1 | 3 | Abs. | W | TSMHG | LPALTLC | - | GVCHTTN | 87 | 263 | - |
| SFRICE001686 | *SfruPPK28* | V | 49 | 490 | 3.36871E-84 | 510 | 2 | 14 | GFS | W | STLHG | FPALTVC | S | GLCCTFN | 71 | 256 | 15 |
| SFRICE031715 | *Sfru031715* | II | 3 | 284 | 3.08233E-17 | 288 | 0 | 7 | Abs. | - | TTLHG | FPAVTAC | - | GMCLTFN | 15 | 373 | - |
| SFRICE012692 | *SfruPPK6a* | IV | 11 | 458 | 3.97386E-20 | 989 | 2 | 12 | GFY | W | STIVG | YPAITLC | T | GYCCTFN | 32 | 592 | 67 |
| SFRICE020475 | *SfruPPK6b* | IV | 4 | 417 | 2.05714E-38 | 444 | 2 | 12 | GFS | W | Abs. | YPAITLC | S | GYCCTFN | 8 | 363 | 22 |
| SFRICE016175 | *SfruPPK6c* | IV | 30 | 434 | 1.45273E-40 | 435 | 1 | 14 | Abs. | W | SMG | YPAITVC | - | GYCCSFN | 35 | 374 | - |
| SFRICE025969 | *SfruPPK9* | II | 44 | 511 | 1.41673E-57 | 524 | 4 | 12 | GCS | W | SSIHG | FPCVTVC | V | GLCYAVN | 66 | 399 | 8 |
| TC001349 | *Tcas001349* | I | 45 | 344 | 4.58819E-30 | 387 | 2 | 13 | GAS | - | Abs. | Abs. | V | GPCFGFN | 1 | 314 | 38 |
| TC002554 | *Tcas002554* | V | 24 | 412 | 6.72673E-83 | 626 | 3 | 11 | GFS | W | CNIHG | FPAVTVC | S | Abs. | 45 | 321 | 209 |
| TC002630 | *Tcas002630* | V | 23 | 421 | 2.24517E-75 | 442 | 2 | 11 | GFS | W | TSIHG | FPAVTIC | A | Abs. | 46 | 329 | 16 |
| TC002631 | *Tcas002631* | V | 16 | 355 | 2.79372E-55 | 356 | 1 | 11 | Abs. | W | TSVHG | FPAVTIC | - | Abs. | 37 | 293 | - |
| TC002632 | *Tcas002632* | V | 32 | 137 | 1.9728E-10 | 188 | 2 | 1 | GFS | W | FSI | FPAVTIC | A | Abs. | 32 | 89 | 16 |
| TC002633 | *Tcas002633* | V | 24 | 402 | 9.16246E-51 | 409 | 0 | 14 | Abs. | - | TGIHG | FPSVTIC | - | GICYSFN | 36 | 372 | - |
| TC003563 | *Tcas003563* | V | 40 | 472 | 5.12028E-69 | 472 | 1 | 14 | Abs. | W | GTIHG | FPAVTIC | - | GICYTFN | 61 | 385 | - |
| TC005542 | *Tcas005542* | V | 27 | 501 | 1.51114E-77 | 520 | 2 | 14 | GFS | W | TGIHG | FPAVTIC | S | GICYSFN | 48 | 407 | 14 |
| TC006569 | *Tcas006569* | V | 22 | 445 | 1.72001E-60 | 465 | 2 | 9 | GFS | W | TSIHG | FPSVTIC | A | GVCYTFN | 43 | 356 | 15 |
| TC006570 | *Tcas006570* | V | 22 | 445 | 1.10653E-61 | 465 | 2 | 9 | GFS | W | TSIHG | FPSVTIC | A | GVCYTFN | 43 | 356 | 15 |
| TC006571 | *Tcas006571* | V | 22 | 509 | 5.89238E-93 | 529 | 2 | 14 | GFS | W | TSIHG | FPSVTIC | A | GVCYTFN | 43 | 420 | 15 |
| TC006572 | *Tcas006572* | V | 21 | 506 | 1.79716E-92 | 526 | 2 | 14 | GFS | W | TSIHG | FPAVTVC | A | GICYTFN | 42 | 418 | 15 |

| | | | | | | | | | | | | | | | | | |
|---|---|---|---|---|---|---|---|---|---|---|---|---|---|---|---|---|---|
| TC006573 | *Tcas006573* | V | 32 | 521 | 1.44702E-94 | 544 | 2 | 14 | GFT | W | TGIHG | FPAVTIC | A | GLCYSFN | 54 | 421 | 18 |
| TC006574 | *Tcas006574* | V | 42 | 523 | 3.3521E-101 | 561 | 2 | 14 | GFS | W | TGIHG | FPAITIC | A | GMCFTFN | 64 | 413 | 33 |
| TC006575 | *Tcas006575* | V | 46 | 531 | 6.06139E-98 | 553 | 2 | 14 | GFS | W | GTIHG | FPAVTIC | A | GICYVFN | 67 | 418 | 17 |
| TC006581 | *Tcas006581* | V | 18 | 485 | 4.09396E-87 | 682 | 2 | 14 | GFS | W | TSIHG | FPAVTIC | A | GVCYSFN | 39 | 409 | 183 |
| TC009302 | *Tcas009302* | IV | 22 | 357 | 9.21663E-62 | 431 | 1 | 9 | GQL | W | TSIHG | FPAVTIC | A | GNCCSFN | 44 | 288 | 66 |
| TC012955 | *Tcas012955* | V | 19 | 497 | 7.26062E-97 | 517 | 2 | 14 | GFS | W | TSIHG | FPAVTIC | A | GICYTFN | 40 | 411 | 15 |
| TC015167 | *TcasPPK27b* | IV | 23 | 264 | 5.16264E-31 | 266 | 0 | 10 | Abs. | - | Abs. | Abs. | - | GFCCVFN | 8 | 255 | - |
| TC015541 | *TcasPPK27a* | IV | 3 | 270 | 1.4099E-30 | 274 | 0 | 10 | Abs. | - | Abs. | Abs. | - | GFCCTFN | 13 | 256 | - |
| TC006095 | *TcasPPK13* | I | 11 | 429 | 3.45182E-68 | 464 | 2 | 14 | GAS | W | TSFHG | FPSISVC | V | GVCYAIN | 33 | 350 | 30 |
| TC013324 | *TcasPPK16* | IV | 19 | 492 | 4.02996E-84 | 548 | 2 | 14 | GFS | W | TALHG | FPAVTIC | A | GYCCAFN | 41 | 405 | 51 |
| TC005481 | *TcasPPK17* | VII | 3 | 360 | 9.25187E-11 | 436 | 2 | 10 | GLS | V | P | YPAITVC | A | GLCHTMI | 6 | 319 | 46 |
| TC012362 | *TcasPPK23* | VI | 34 | 432 | 1.21846E-59 | 522 | 2 | 14 | Abs. | W | STLHG | FPAITVC | - | GFCYTFN | 56 | 344 | 85 |
| TC002095 | *TcasPPK28* | V | 130 | 592 | 1.44132E-86 | 627 | 2 | 14 | GFS | W | STLHG | FPSITIC | S | GLCCSFN | 152 | 394 | 30 |
| TC004432 | *TcasPPK3* | II | 2 | 312 | 2.76943E-25 | 341 | 2 | 11 | GIS | - | Abs. | FPAVTGC | V | Abs. | - | 274 | - |
| TC002260 | *TcasPPK9* | II | 18 | 382 | 2.33817E-41 | 402 | 2 | 12 | GCS | W | SSIHG | FPAATIC | - | GICYSFN | 40 | 303 | 15 |

**Table S3. Calmodulin Binding Motif (CBM) analysis.** SF: Subfamily; TM: transmembrane domains. Length is expressed in number of amino acids.

| PPK | SF | Length | Start 1st TM | Finish 2nd TM | CBM location | Start | End | CBS Length | CBM sequence |
|---|---|---|---|---|---|---|---|---|---|
| *AaegPPK302* | I | 481 | 41 | 456 | C-terminal | 586 | 598 | 12 | AGWMDGAFRLRKK |
| *AalbPPK15a* | I | 482 | 41 | 456 | N-terminal | 27 | 43 | 16 | LGYIASRKYHYTERLFW |
| *AalbPPK15b* | I | 499 | 41 | 473 | N-terminal | 27 | 43 | 16 | LGYIASRKYHYTERLFW |
| *AalbPPK31* | I | 332 | 2 | 325 | C-terminal | 585 | 596 | 11 | ITAWMDGAFRLR |
| *AgamPPK15a* | I | 489 | 40 | 456 | N-terminal | 25 | 40 | 15 | GVGYISNRKYHWTERL |
| *Amel53731* | I | 421 | 52 | 370 | N-terminal | 8 | 20 | 12 | WKIIKQYLTNCSI |
| *Bger41668* | I | 295 | 74 | 285 | N-terminal | 19 | 28 | 9 | ILSWWRKKSL |
| *Bimp012246752* | I | 421 | 38 | 417 | N-terminal | 10 | 20 | 10 | KIIKQYLTNCT |
| *BmorPPK13b* | I | 509 | 44 | 446 | N-terminal | 37 | 48 | 11 | KRHWVERIFWLV |
| *Clec000454* | I | 635 | 260 | 598 | N-terminal | 25 | 35 | 10 | RKRLYGVSSSK |
| *CquiPPK13* | I | 730 | 307 | 711 | N-terminal | 71 | 82 | 11 | KRNVVRGRLDLV |
| *CquiPPK31* | I | 414 | 10 | 330 | C-terminal | 395 | 410 | 15 | VAFLGFHRFRTGAANK |
| *DmelPPK15* | I | 483 | 50 | 471 | N-terminal | 11 | 22 | 11 | VLRKKRGFGFVT |
| *Dplex215856* | I | 262 | 2 | 251 | C-terminal | 243 | 260 | 17 | VFYYICFKQWKYIRERTT |
| *AaegPPK201* | II | 438 | 37 | 424 | N-terminal | 5 | 21 | 16 | FYISKFFRRVISKSSLH |
| *Agam007084* | II | 481 | 37 | 440 | N-terminal | 28 | 42 | 14 | HKRSTYVEKVIWLGI |
| *AgamPPK25* | II | 431 | 29 | 414 | C-terminal | 414 | 421 | 7 | ITQKVYRR |
| *AgamPPK9* | II | 459 | 44 | 457 | N-terminal | 10 | 23 | 13 | LVKRSKRFVVNLLT |
| *ApisPPK9* | II | 494 | 67 | 486 | N-terminal | 27 | 41 | 14 | DRIIVKLKTNVFNFV |
| *Bger41670* | II | 438 | 2 | 349 | C-terminal | 418 | 429 | 11 | IRLRLKTGKLDV |
| *BmorPPK9* | II | 480 | 70 | 473 | N-terminal | 37 | 46 | 9 | TKKAKKYLVL |
| *DmelPPK25* | II | 451 | 40 | 437 | N-terminal | 31 | 46 | 15 | RRDLHWAERLFWTFII |
| *DplexPPK25* | II | 376 | 31 | 352 | N-terminal | 24 | 39 | 15 | RRHWSERLLWVCFIVA |
| *MperPPK9a* | II | 480 | 67 | 472 | N-terminal | 26 | 42 | 16 | KDRIFIRLKKGVFNFVS |
| *MperPPK9b* | II | 462 | 67 | 454 | N-terminal | 26 | 42 | 16 | QDRIFIKLKKGVFNFAW |
| *TcasPPK9* | II | 402 | 41 | 387 | C-terminal | 382 | 397 | 15 | FFTLRLYWFVIKYGKE |
| *Apis34462* | III | 282 | 2 | 263 | C-terminal | 259 | 272 | 13 | ITFRLYDYWLNRRH |
| *Bger30177* | III | 375 | 2 | 326 | C-terminal | 327 | 336 | 9 | FGVWRSEKRV |
| *DmelPPK21* | III | 550 | 76 | 489 | C-terminal | 525 | 549 | 24 | SIGLYLYIHGKRKLR |
| *DmelPPK7* | III | 573 | 61 | 498 | N-terminal | 46 | 55 | 9 | GLDRLLSAKA |
| *Mper000138710.1* | III | 460 | 40 | 441 | N-terminal | 31 | 41 | 10 | IRAKNVGERRF |
| *Mper000154690.1* | III | 539 | 62 | 520 | N-terminal | 21 | 36 | 15 | ERLAIAKSKRLGNIF |
| *Acep17448* | IV | 466 | 10 | 428 | C-terminal | 421 | 436 | 15 | VYLLARQLFRRQKKML |
| *Acep19650* | IV | 513 | 107 | 501 | N-terminal | 95 | 112 | 17 | FFKKEKKKKTKKIIMWVI |
| *Agam006703* | IV | 529 | 51 | 450 | C-terminal | 451 | 466 | 15 | FTVYSALRNRNGKAVG |
| *Agam006704* | IV | 565 | 52 | 482 | C-terminal | 525 | 538 | 16 | FSRRNVLARERSAG |
| *AgamPPK16* | IV | 533 | 67 | 513 | C-terminal | 513 | 525 | 12 | LLFDAIAKKSDKR |
| *AgamPPK6* | IV | 557 | 99 | 530 | C-terminal | 524 | 539 | 15 | FFSIFRLRKMYHKLKQ |

| | | | | | | | | | |
|---|---|---|---|---|---|---|---|---|---|
| *Amel48330* | IV | 571 | 57 | 509 | N-terminal | 16 | 24 | 8 | LRKGCLSKF |
| *BimpPPK16* | IV | 528 | 54 | 512 | C-terminal | 507 | 521 | 14 | FFIIRVLTDACVKRN |
| *Bmor012545990.1* | IV | 439 | 25 | 404 | N-terminal | 16 | 23 | 7 | KLRFENRF |
| *Cqui005430* | IV | 500 | 60 | 417 | C-terminal | 433 | 448 | 15 | NVMDVRGRSRVGQGF |
| *CquiPPK6a* | IV | 449 | 71 | 364 | N-terminal | 61 | 71 | 10 | RRNVTTGFARF |
| *CquiPPK6b* | IV | 464 | 71 | 429 | N-terminal | 61 | 71 | 10 | RRNVTTGFARF |
| *DmelPPK11* | IV | 516 | 117 | 507 | N-terminal | 58 | 65 | 7 | WYNRVSKR |
| *DmelPPK16* | IV | 531 | 82 | 523 | N-terminal | 73 | 86 | 13 | RQDISRHERWFWLV |
| *DmelPPK22* | IV | 561 | 67 | 532 | C-terminal | 525 | 540 | 15 | LFFVTKYIYKGCNRMV |
| *DmelPPK24* | IV | 595 | 87 | 556 | N-terminal | 80 | 90 | 10 | GRRIQERFFWF |
| *DmelPPK27* | IV | 422 | 38 | 412 | N-terminal | 20 | 32 | 12 | SLNGFGLLYFIRK |
| *DmelPPK6* | IV | 498 | 64 | 481 | N-terminal | 55 | 64 | 9 | RRNRTYGLSRF |
| *MdomPPK16* | IV | 569 | 114 | 560 | N-terminal | 104 | 119 | 14 | TRMDLSKNERIFWLII |
| *MdomPPK4* | IV | 507 | 52 | 473 | C-terminal | 471 | 485 | 14 | VILKKFYRQEKNARE |
| *MdomPPK6* | IV | 483 | 53 | 470 | N-terminal | 15 | 23 | 8 | WKRFRLAL |
| *Pxyl34278.t1* | IV | 540 | 17 | 429 | C-terminal | 437 | 444 | 7 | FMTKIYRK |
| *SfruPPK6b* | IV | 444 | 9 | 422 | C-terminal | 421 | 434 | 13 | GLKNYVKNKLAKVF |
| *AaegPPK313* | V | 599 | 43 | 561 | N-terminal | 26 | 36 | 10 | HLANRRLTLFE |
| *AaegPPK317* | V | 544 | 43 | 514 | N-terminal | 34 | 48 | 14 | CRERSICEKFWWVVV |
| *Aalb019525011.2* | V | 568 | 57 | 529 | N-terminal | 48 | 62 | 14 | CRERSLCEKFWWVVV |
| *Aalb029712101.1* | V | 617 | 66 | 581 | N-terminal | 27 | 40 | 13 | YNFAKEVKQIRKRG |
| *Aalb029732095.1* | V | 577 | 77 | 552 | N-terminal | 68 | 78 | 10 | SRRRSILERIW |
| *Aalb029732188.1* | V | 448 | 58 | 445 | N-terminal | 45 | 61 | 16 | KRALNKELSWLERLFWL |
| *Agam011610* | V | 575 | 65 | 530 | N-terminal | 56 | 67 | 11 | GNKRTTIERIWW |
| *Agam011611* | V | 568 | 67 | 529 | N-terminal | 57 | 66 | 9 | VGNGRTRVEK |
| *Apis29894* | V | 578 | 68 | 525 | C-terminal | 524 | 535 | 11 | RLYVDRWNKNKK |
| *Apis35976* | V | 588 | 87 | 563 | C-terminal | 563 | 579 | 16 | PLVNFCNAKRKLKKVDP |
| *BimpPPK28* | V | 658 | 170 | 625 | N-terminal | 72 | 88 | 16 | PLRLEGYATADIFKVRP |
| *Bmor012552731.1* | V | 456 | 3 | 441 | C-terminal | 437 | 450 | 13 | ITLRIWCALAREKR |
| *Cqui003410* | V | 569 | 65 | 508 | N-terminal | 57 | 65 | 8 | KRRTLYEKI |
| *Cqui005125* | V | 518 | 34 | 490 | C-terminal | 495 | 510 | 15 | KEGTCHNGKVVIVIVK |
| *Cqui005126* | V | 577 | 78 | 550 | N-terminal | 70 | 80 | 10 | RQRSLFEKFWW |
| *Cqui005127* | V | 568 | 75 | 547 | N-terminal | 65 | 79 | 14 | GSRHRTLFEKFWWVA |
| *Cqui005128* | V | 545 | 74 | 520 | N-terminal | 68 | 76 | 8 | RSRWERLWW |
| *Cqui005908* | V | 543 | 42 | 487 | C-terminal | 521 | 528 | 7 | LDAGIRRR |
| *Cqui007806* | V | 565 | 56 | 521 | N-terminal | 38 | 55 | 17 | SIHGVKYFVGSNRALIEK |
| *Cqui007807* | V | 505 | 6 | 471 | C-terminal | 470 | 487 | 17 | KPLLMWWRGDIGVKQVGP |
| *Cqui007808* | V | 541 | 43 | 506 | C-terminal | 509 | 522 | 13 | LWWKELNKVETVKQ |
| *Cqui007809* | V | 557 | 46 | 511 | C-terminal | 538 | 552 | 14 | ASKILHAGYKSDVVG |
| *Cqui007810* | V | 530 | 52 | 504 | N-terminal | 11 | 28 | 17 | SPLRRKVLLKLSKQAG |

| | | | | | | | | | |
|---|---|---|---|---|---|---|---|---|---|
| | | | | | | | | | YE |
| *Cqui010135* | V | 469 | 66 | 466 | N-terminal | 44 | 54 | 10 | LRNTSFFGIKY |
| *Cqui010136* | V | 563 | 90 | 543 | N-terminal | 31 | 50 | 19 | IKSPVGVHNFGNGLIKRLGN |
| *Cqui012545* | V | 547 | 44 | 518 | N-terminal | 34 | 45 | 11 | RGSERTWYEKLV |
| *Cqui012546* | V | 544 | 46 | 514 | N-terminal | 38 | 50 | 12 | RERSGCEKFWWFA |
| *CquiPPK301* | V | 574 | 45 | 562 | N-terminal | 25 | 42 | 17 | RQRTESYDWAGKGFAILL |
| *DmelPPK2* | V | 562 | 62 | 530 | N-terminal | 13 | 29 | 16 | GIDLTFRRRRKAGSVAC |
| *DmelPPK26* | V | 597 | 97 | 575 | C-terminal | 575 | 585 | 10 | LISNLRMRRKT |
| *DmelPPK28* | V | 632 | 91 | 540 | N-terminal | 28 | 46 | 18 | NTICSRAAIKRSVVYYLKN |
| *DmelPPK8* | V | 569 | 65 | 513 | N-terminal | 56 | 68 | 12 | NSKLRSSDRLFFG |
| *Dplex216041* | V | 1034 | 65 | 520 | C-terminal | 792 | 805 | 13 | TGFGAKSGLTFLLK |
| *Dpon71305* | V | 484 | 63 | 448 | N-terminal | 53 | 64 | 11 | IGERRSLCEKVS |
| *GmorPPK28* | V | 516 | 41 | 478 | C-terminal | 497 | 505 | 8 | KARYRIYLT |
| *Lmig14820.1* | V | 465 | 26 | 392 | C-terminal | 389 | 402 | 13 | ARLGFSQLLKRSR |
| *MdomPPK2c* | V | 574 | 88 | 566 | N-terminal | 70 | 90 | 20 | ERRPTLDRFIWFALILISCF |
| *MdomPPK2i* | V | 554 | 68 | 543 | N-terminal | 59 | 72 | 13 | GQRRQRKEVGFWII |
| *MdomPPK8b* | V | 601 | 61 | 512 | N-terminal | 14 | 29 | 15 | VIIGDKKKSRWNSIKV |
| *Pxyl36720.t1* | V | 382 | 7 | 360 | C-terminal | 358 | 371 | 13 | LRLCCILWRRRRSK |
| *Rpro000048* | V | 543 | 47 | 504 | C-terminal | 505 | 513 | 8 | GNMKATKK |
| *SfruPPK28* | V | 510 | 72 | 495 | N-terminal | 62 | 71 | 9 | GERKLTWFER |
| *Tcas002632* | V | 188 | 33 | 172 | C-terminal | 230 | 239 | 9 | LAKKSEQFS |
| *Tcas006569* | V | 465 | 44 | 450 | N-terminal | 33 | 45 | 12 | FGEKRTIFERIW |
| *Tcas006570* | V | 465 | 44 | 450 | N-terminal | 34 | 45 | 11 | FGEKRTIFERIW |
| *Tcas006571* | V | 529 | 44 | 514 | N-terminal | 35 | 45 | 10 | GEKRTIFERVW |
| *Tcas006572* | V | 526 | 43 | 511 | N-terminal | 33 | 45 | 12 | FGEKRTIFERIWW |
| *Tcas006573* | V | 544 | 55 | 526 | N-terminal | 40 | 57 | 17 | MKYLGEQNRTIAEKIFW |
| *Tcas006581* | V | 682 | 40 | 499 | N-terminal | 32 | 44 | 12 | EKRSKIEKTIWSL |
| *Tcas012955* | V | 517 | 41 | 502 | N-terminal | 29 | 42 | 13 | RYFGEKRTIFERIW |
| *AaegPPK101* | VI | 618 | 74 | 478 | N-terminal | 33 | 49 | 16 | PFGTVRLLKGSLLYQTK |
| *AaegPPK102* | VI | 576 | 44 | 436 | C-terminal | 524 | 539 | 15 | AKRYISRYERIASMAK |
| *AalbPPK23b* | VI | 623 | 76 | 480 | N-terminal | 35 | 51 | 16 | PFGTIRLLKGSLLYQTK |
| *AgamPPK23* | VI | 656 | 87 | 507 | C-terminal | 618 | 631 | 13 | QLGHANYAGSKYAE |
| *CquiPPK23b* | VI | 610 | 70 | 473 | N-terminal | 29 | 42 | 13 | LFNGFRLFRSSLLY |
| *PxylPPK23b* | VI | 704 | 63 | 627 | N-terminal | 15 | 30 | 15 | RKVSDKTQAGGLSLL |
| *RproPPK23* | VI | 552 | 81 | 484 | N-terminal | 20 | 47 | 27 | VAVQTGINFFSGMHRALR |
| *AalbPPK17b* | VII | 443 | 16 | 392 | C-terminal | 487 | 499 | 12 | LFIRNLIAEKRNR |
| *BmorPPK17* | VII | 414 | 15 | 393 | C-terminal | 388 | 401 | 13 | LLKKLGISFVLDRK |
| *CquiPPK17* | VII | 436 | 15 | 391 | C-terminal | 382 | 400 | 18 | IVELLFIRRLIAEKVSKRK |
| *GmorPPK17* | VII | 455 | 27 | 415 | C-terminal | 413 | 430 | 17 | GASRNGLVKRVKHVNENR |

**Table S4. Genomic location of PPK genes and cluster characteristics.** PPKs located in clusters are depicted in red. Exp: Expansion.

| Annotation | Original code | Chromosome | Start | Stop | Gene length (bp) | PPK distance (bp) | SF | Cluster N° | Genes per Cluster | Cluster Length (bp) | Exp. |
|---|---|---|---|---|---|---|---|---|---|---|---|
| AaegPPK101 | AAEL014009-PA | 1 | 57568844 | 57583811 | 14967 | 28398 | VI | 1 | 2 | 57712 | |
| AaegPPK102 | AAEL014010-PB | 1 | 57612209 | 57626556 | 14347 | 170819739 | VI | | | | |
| AaegPPK103 | AAEL002575-PB | 1 | 228446295 | 228482750 | 36455 | | IV | | | | |
| AaegPPK201 | AAEL022385-PA | 2 | 95598504 | 95600146 | 1642 | 79419515 | II | | | | |
| AaegPPK202 | AAEL019946-PB | 2 | 175019661 | 175035492 | 15831 | 22581693 | II | | | | |
| AaegPPK203 | AAEL003714-PB | 2 | 197617185 | 197630564 | 13379 | 107435353 | IV | | | | |
| AaegPPK204 | AAEL010779-PA | 2 | 305065917 | 305077405 | 11488 | 9346025 | IV | | | | |
| AaegPPK205 | AAEL022016-PA | 2 | 314423430 | 314462552 | 39122 | 148565072 | II | | | | |
| AaegPPK206 | AAEL006613-PA | 2 | 463027624 | 463039234 | 11610 | | I | | | | |
| AaegPPK301 | AAEL000582-PC | 3 | 11869920 | 11871999 | 2079 | 50787579 | V | | | | |
| AaegPPK302 | AAEL006258-PA | 3 | 62659578 | 62662148 | 2570 | 70352170 | I | | | | |
| AaegPPK303 | AAEL008809-PB | 3 | 133014318 | 133016180 | 1862 | 73373836 | II | | | | |
| AaegPPK29 | AAEL023544-PA | 3 | 206390016 | 206401577 | 11561 | 14931360 | I | | | | |
| AaegPPK305 | AAEL003470-PB | 3 | 221332937 | 221337955 | 5018 | 66133947 | IV | | | | |
| AaegPPK306 | AAEL014228-PA | 3 | 287471902 | 287473808 | 1906 | 31048 | V | 2 | 7 | 176202 | Culicidae Exp. |
| AaegPPK307 | AAEL026145-PA | 3 | 287504856 | 287505518 | 662 | 1122 | V | | | | Culicidae Exp. |
| AaegPPK308 | AAEL014230-PA | 3 | 287506640 | 287508475 | 1835 | 17646 | V | | | | Culicidae Exp. |
| AaegPPK309 | AAEL011005-PA | 3 | 287526121 | 287527835 | 1714 | 14774 | V | | | | Culicidae Exp. |
| AaegPPK310 | AAEL011002-PB | 3 | 287542609 | 287543377 | 768 | 7230 | V | | | | Culicidae Exp. |
| AaegPPK311 | AAEL026983-PA | 3 | 287550607 | 287552334 | 1727 | 93901 | V | | | | Culicidae Exp. |
| AaegPPK312 | AAEL010995-PB | 3 | 287646235 | 287648104 | 1869 | 9228364 | V | | | | Culicidae Exp. |
| AaegPPK313 | AAEL019676-PA | 3 | 296876468 | 296890541 | 14073 | 4776 | V | 3 | 2 | 20328 | Culicidae Exp. |
| AaegPPK314 | AAEL019675-PA | 3 | 296895317 | 296896796 | 1479 | 36965676 | V | | | | Culicidae Exp. |
| AaegPPK315 | AAEL000863-PA | 3 | 333862472 | 333863955 | 1483 | 827824 | V | | | | Culicidae Exp. |
| AaegPPK316 | AAEL000926-PB | 3 | 334691779 | 334693491 | 1712 | 26840 | V | 4 | 2 | 30305 | Culicidae Exp. |
| AaegPPK317 | AAEL000873-PB | 3 | 334720331 | 334722084 | 1753 | 24944461 | V | | | | Culicidae Exp. |

| | | | | | | | | | | | |
|---|---|---|---|---|---|---|---|---|---|---|---|
| AaegPPK318 | AAEL008053-PA | 3 | 359666545 | 359668314 | 1769 | 25107522 | V | | | | Culicidae Exp. |
| AaegPPK319 | AAEL000534-PB | 3 | 384775836 | 384777088 | 1252 | 8461 | V | 5 | 3 | 59970 | Culicidae Exp. |
| AaegPPK320 | AAEL000552-PB | 3 | 384785549 | 384787254 | 1705 | 46665 | V | | | | Culicidae Exp. |
| AaegPPK321 | AAEL000547-PA | 3 | 384833919 | 384835806 | 1887 | 17228343 | V | | | | Culicidae Exp. |
| AaegPPK322 | AAEL004091-PB | 3 | 402064149 | 402075834 | 11685 | | V | | | | Culicidae Exp. |
| AaegPPK323 (304) | AAEL027684-PA | NIGP01001847 | 29788 | 34801 | 5013 | | IV | | | | |

| Annotation | Original code | Scaffold | Start | Stop | Gene length (bp) | PPK distance (bp) | SF | Cluster N° | Genes per Cluster | Cluster Length (bp) | Exp. |
|---|---|---|---|---|---|---|---|---|---|---|---|
| AalbPPK15a | XP_019558124.2 | NW_021837045.1 | 73424556 | 73426188 | 1632 | 1804421 | I | | | | |
| AalbPPK15b | XP_029726981.1 | NW_021837045.1 | 71618503 | 71620135 | 1632 | 109347188 | I | | | | |
| AalbPPK9 | XP_029735319.1 | NW_021837045.1 | 182773376 | 182774572 | 1196 | | II | | | | |
| AalbPPK301A | XP_019551763.2 | NW_021837257.1 | 9765 | 11861 | 2096 | | V | | | | |
| AalbPPK13a | XP_029720145.1 | NW_021837378.1 | 30373505 | 30391433 | 17928 | | I | | | | |
| Aalb029721586.1 | XP_029721586.1 | NW_021837489.1 | 47765963 | 47767639 | 1676 | 18732 | V | 1 | 2 | 783674 | Culicid Exp. |
| Aalb029720868.1 | XP_029720868.1 | NW_021837489.1 | 47786371 | 47788250 | 1879 | 532898 | V | | | | Culicid Exp. |
| Aalb029721587.1 | XP_029721587.1 | NW_021837489.1 | 48321148 | 48338567 | 17419 | 231 | V | 2 | 8 | 228489 | Culicid Exp. |
| Aalb029721588.1 | XP_029721588.1 | NW_021837489.1 | 48338798 | 48339399 | 601 | 17938 | V | | | | Culicid Exp. |
| Aalb029720869.1 | XP_029720869.1 | NW_021837489.1 | 48357337 | 48359179 | 1842 | 6983 | V | | | | Culicid Exp. |
| Aalb029721589.1 | XP_029721589.1 | NW_021837489.1 | 48366162 | 48367879 | 1717 | 11566 | V | | | | Culicid Exp. |
| Aalb029721590.1 | XP_029721590.1 | NW_021837489.1 | 48379445 | 48380670 | 1225 | 5224 | V | | | | Culicid Exp. |
| Aalb019556948.2 | XP_019556948.2 | NW_021837489.1 | 48385894 | 48387615 | 1721 | 10773 | V | | | | Culicid Exp. |
| Aalb019556935.2 | XP_019556935.2 | NW_021837489.1 | 48398388 | 48400170 | 1782 | 59037 | V | | | | Culicid Exp. |
| Aalb029721592.1 | XP_029721592.1 | NW_021837489.1 | 48459207 | 48549637 | 90430 | | V | | | | Culicid Exp. |
| Aalb019557794.2 | XP_019557794.2 | NW_021838153.1 | 5986060 | 5987684 | 1624 | | II | | | | |
| AalbPPK25 | XP_019540188.2 | NW_021838343.1 | 8822904 | 8825703 | 2799 | | II | | | | |
| AalbPPK13b | XP_029729201.1 | NW_021838387.1 | 734852 | 765626 | 30774 | | I | | | | |
| AalbPPK17a | XP_019555869.2 | NW_021838399.1 | 13667348 | 13676355 | 9007 | 257695 | VII | | | | |
| AalbPPK17b | XP_029729547.1 | NW_021838399.1 | 13934050 | 13949369 | 15319 | | VII | | | | |
| AalbPPK3 | XP_029730550.1 | NW_021838454.1 | 2937882 | 2955109 | 17227 | | II | | | | |
| Aalb029731012.1 | XP_029731012.1 | NW_021838465.1 | 33832180 | 33834260 | 2080 | 19602 | V | 3 | 3 | 887974 | Culicid Exp. |
| Aalb019525011.2 | XP_019525011.2 | NW_021838465.1 | 33853862 | 33855635 | 1773 | 18732 | V | | | | Culicid Exp. |
| Aalb029731018.1 | XP_029731018.1 | NW_021838465.1 | 34718652 | 34720154 | 1502 | | V | | | | Culicid Exp. |
| Aalb029732188.1 | XP_029732188.1 | NW_021838465.1 | 49420555 | 49422021 | 1466 | 18441727 | V | | | | Culicid Exp. |
| Aalb019537219.2 | XP_019537219.2 | NW_021838465.1 | 67863748 | 67865514 | 1766 | 856399 | V | | | | Culicid Exp. |
| Aalb029731325.1 | XP_029731325.1 | NW_021838465.1 | 68721913 | 68723676 | 1763 | 31313225 | V | | | | Culicid Exp. |
| Aalb029731663.1 | XP_029731663.1 | NW_021838465.1 | 100036901 | 100038154 | 1253 | 35828 | V | 4 | 2 | 38815 | Culicid Exp. |
| Aalb029731661.1 | XP_029731661.1 | NW_021838465.1 | 100073982 | 100075716 | 1734 | 3709343 | V | | | | Culicid Exp. |
| Aalb029732095.1 | XP_029732095.1 | NW_021838465.1 | 103785059 | 103786916 | 1857 | 5549 | V | 5 | 4 | 52662 | Culicid Exp. |
| Aalb029732096.1 | XP_029732096.1 | NW_021838465.1 | 103792465 | 103794737 | 2272 | 21034 | V | | | | Culicid Exp. |

| | | | | | | | | | | | |
|---|---|---|---|---|---|---|---|---|---|---|---|
| Aalb029731672.1 | XP_029731672.1 | NW_021838465.1 | 103815771 | 103817505 | 1734 | 18322 | V | | | | Culicid Exp. |
| Aalb019539770.2 | XP_019539770.2 | NW_021838465.1 | 103835827 | 103837721 | 1894 | | V | | | | Culicid Exp. |
| Aalb019536370.2 | XP_019536370.2 | NW_021838465.1 | 133279436 | 133293104 | 13668 | | V | | | | Culicid Exp. |
| AalbPPK29b | XP_029733934.1 | NW_021838554.1 | 4845254 | 4857047 | 11793 | 1876340 | I | | | | |
| AalbPPK29c | XP_029733926.1 | NW_021838554.1 | 6733387 | 6738657 | 5270 | | I | | | | |
| Aalb029734094.1 | XP_029734094.1 | NW_021838557.1 | 12381 | 14117 | 1736 | 15856 | V | 6 | 3 | 32661 | Culicid Exp. |
| Aalb029734091.1 | XP_029734091.1 | NW_021838557.1 | 29973 | 35014 | 5041 | 8520 | V | | | | Culicid Exp. |
| Aalb029734092.1 | XP_029734092.1 | NW_021838557.1 | 43534 | 45042 | 1508 | | V | | | | Culicid Exp. |
| Aalb029734497.1 | XP_029734497.1 | NW_021838576.1 | 49511189 | 49531703 | 20514 | 773274 | IV | | | | |
| Aalb029735277.1 | XP_029735277.1 | NW_021838576.1 | 50304977 | 50325921 | 20944 | | IV | | | | |
| AalbPPK301b | XP_029707777.1 | NW_021838621.1 | 737127 | 739222 | 2095 | | V | | | | |
| AalbPPK29a | XP_029709904.1 | NW_021838719.1 | 20955 | 25394 | 4439 | | I | | | | |
| AalbPPK23a | XP_029711542.1 | NW_021838798.1 | 41023749 | 41043474 | 19725 | 13877 | VI | 7 | 2 | 39371 | |
| AalbPPK23b | XP_019530003.2 | NW_021838798.1 | 41057351 | 41063120 | 5769 | | VI | | | | |
| Aalb029712101.1 | XP_029712101.1 | NW_021838821.1 | 3637597 | 3639568 | 1971 | | V | | | | Culicid Exp. |
| Aalb029713859.1 | XP_029713859.1 | NW_021838914.1 | 200692 | 202734 | 2042 | | II | | | | |
| AalbPPK10 | XP_029715164.1 | NW_021839020.1 | 21387413 | 21419469 | 32056 | | II | | | | |
| AalbPPK31 | XP_029715424.1 | NW_021839026.1 | 100629 | 112120 | 11491 | | I | | | | |

| **Annotation** | **Original code** | **Chromosome** | **Start** | **Stop** | **Gene length (bp)** | **PPK distance (bp)** | **SF** | **Cluster N°** | **Genes per Cluster** | **Cluster Length (bp)** | **Exp.** |
|---|---|---|---|---|---|---|---|---|---|---|---|
| AgamPPK13 | AGAP007945 | 3R | 3250860 | 3252530 | 1670 | 6866383 | I | | | | |
| AgamPPK15a | AGAP008378 | 3R | 10118913 | 10120576 | 1663 | 11043 | I | 1 | 2 | 15647 | |
| AgamPPK15b | AGAP008380 | 3R | 10131619 | 10134560 | 2941 | 26584037 | I | | | | |
| AgamPPK16 | AGAP009590 | 3R | 36718597 | 36720400 | 1803 | 6576938 | IV | | | | |
| AgamPPK10 | AGAP009789 | 3R | 43297338 | 43299700 | 2362 | 6135139 | II | | | | |
| Agamppk17 | AGAP010146 | 3R | 49434839 | 49436537 | 1698 | | VII | | | | |
| AgamPPK6 | AGAP010430 | 3L | 3193890 | 3196118 | 2228 | 10773901 | IV | | | | |
| Agam028699 | AGAP028699 | 3L | 13970019 | 13971752 | 1733 | 4788 | V | 2 | 5 | 17743 | Culicid Exp. |
| Agam028700 | AGAP028700 | 3L | 13976540 | 13978272 | 1732 | 681 | V | | | | Culicid Exp. |
| Agam028701 | AGAP028701 | 3L | 13978953 | 13980751 | 1798 | 2430 | V | | | | Culicid Exp. |
| Agam028702 | AGAP028702 | 3L | 13983181 | 13984969 | 1788 | 1275 | V | | | | Culicid Exp. |
| Agam028703 | AGAP028703 | 3L | 13986244 | 13987762 | 1518 | 2966836 | V | | | | Culicid Exp. |
| Agam011103 | AGAP011103 | 3L | 16954598 | 16956643 | 2045 | 7632010 | V | | | | Culicid Exp. |
| Agam011433 | AGAP011433 | 3L | 24588653 | 24590133 | 1480 | 5104270 | V | | | | Culicid Exp. |
| Agam011610 | AGAP011610 | 3L | 29694403 | 29696388 | 1985 | 3010 | V | 3 | 2 | 6943 | Culicid Exp. |
| Agam011611 | AGAP011611 | 3L | 29699398 | 29701346 | 1948 | 10022200 | V | | | | Culicid Exp. |
| Agam012279 | AGAP012279 | 3L | 39723546 | 39725365 | 1819 | | V | | | | Culicid Exp. |
| AgamPPK9 | AGAP004474 | 2R | 56741249 | 56744267 | 3018 | 49702423 | II | | | | |
| AgamPPK301 | AGAP001602 | 2R | 6641935 | 6644418 | 2483 | 389885 | V | | | | |
| Agam001631 | AGAP001631 | 2R | 7034303 | 7038826 | 4523 | | IV | | | | |
| AgamPPK25 | AGAP005516 | 2L | 16554936 | 16558482 | 3546 | 20531320 | II | | | | |
| Agam006703 | AGAP006703 | 2L | 37089802 | 37091803 | 2001 | 651 | IV | 4 | 3 (MIXED) | 76527 | |
| Agam006704 | AGAP006704 | 2L | 37092454 | 37094562 | 2108 | 67486 | IV | | | | |
| AgamPPK3 | AGAP006720 | 2L | 37162048 | 37166329 | 4281 | 5114075 | II | | | | |

| | | | | | | | |
|---|---|---|---|---|---|---|---|
| Agam007084 | AGAP007084 | 2L | 42280404 | 42282090 | 1686 | | II |
| AgamPPK31 | AGAP000657 | X | 11718049 | 11720785 | 2736 | 3816210 | I |
| AgamPPK23 | AGAP000840 | X | 15536995 | 15539148 | 2153 | | VI |

| Annotation | Original code | Scaffold | Start | Stop | Gene length (bp) | PPK distance (bp) | SF | Cluster N° | Genes per Cluster | Cluster Length (bp) | Exp. |
|---|---|---|---|---|---|---|---|---|---|---|---|
| Agla006813 | AGLA006813 | KB933596 | 686098 | 728445 | 42347 | 232640 | IV | | | | Coleopteran Exp. |
| AglaPPK27 | AGLA006011 | KB933606 | 961085 | 964108 | 3023 | | IV | | | | |
| Agla013191 | AGLA013191 | KB933815 | 307003 | 320703 | 13700 | 5646 | V | 1 | 3 | 50202 | |
| Agla013193 | AGLA013193 | KB933815 | 326349 | 337065 | 10716 | 6356 | V | | | | |
| Agla013194 | AGLA013194 | KB933815 | 343421 | 357205 | 13784 | | V | | | | Coleopteran Exp. |
| Agla012530 | AGLA012530 | KB933877 | 199257 | 205613 | 6356 | 57596 | V | 2 | 3 | 79.382 | Coleopteran Exp. |
| Agla012532 | AGLA012532 | KB933877 | 263209 | 278639 | 15430 | 86963 | V | | | | Coleopteran Exp. |
| Agla012536 | AGLA012536 | KB933877 | 365602 | 388420 | 22818 | | V | | | | |
| Agla014384 | AGLA014384 | KB934189 | 171778 | 218837 | 47059 | 4791 | V | 3 | 4 | 130459 | Coleopteran Exp. |
| Agla014385 | AGLA014385 | KB934189 | 223628 | 247258 | 23630 | 2 | V | | | | Coleopteran Exp. |
| Agla014386 | AGLA014386 | KB934189 | 247260 | 254274 | 7014 | 13752 | V | | | | Coleopteran Exp. |
| Agla014387 | AGLA014387 | KB934189 | 268026 | 302237 | 34211 | | V | | | | |
| AglaPPK28 | AGLA09103 | KB934660 | 16077 | 51768 | 35691 | | V | | | | |
| AglaPPK13 | AGLA003479 | KB933651 | 11233 | 117231 | 105998 | | I | | | | |
| AglaPPK16 | AGLA005467 | KB933540 | 213957 | 222424 | 8467 | | IV | | | | |
| AglaPPK3 | AGLA017497 | KB934892 | 110087 | 115447 | 5360 | | II | | | | |
| AglaPPK9 | AGLA016101 | KB934338 | 16009 | 19490 | 3481 | | II | | | | |

| Annotation | Original code | Scaffold | Start | Stop | Gene length (bp) | PPK distance (bp) | SF | Cluster N° | Genes per Cluster | Cluster Length (bp) | Exp. |
|---|---|---|---|---|---|---|---|---|---|---|---|
| Amel55337 | GB55337-PA | LG1 | 17944914 | 17952616 | 7702 | | IV | | | | |
| AmelPPK23 | GB53179-PA | LG1 | 4727922 | 4730264 | 2342 | | VI | | | | |
| Amel48330 | GB48330-PA | LG10 | 5620602 | 5627975 | 7373 | 734412 | IV | | | | |
| Amel48363 | GB48363-PA | LG10 | 6362387 | 6364884 | 2497 | | IV | | | | |
| AmelPPK28 | GB46186-PA | LG15 | 1811999 | 1816744 | 4745 | | V | | | | |
| Amel53731 | GB53731-PA | LG3 | 8374093 | 8378042 | 3949 | | I | | | | |
| AmelPPK16 | GB45440-PA | LG6 | 16364186 | 16368457 | 4271 | | IV | | | | |
| Amel53792 | GB53792-PA | LG7 | 2981805 | 2984338 | 2533 | | II | | | | |

| Annotation | Original code | Scaffold | Start | Stop | Gene length (bp) | PPK distance (bp) | SF | Cluster N° | Genes per Cluster | Cluster Length (bp) | Exp. |
|---|---|---|---|---|---|---|---|---|---|---|---|
| ApisPPK28 | ACYPI50440 | GL349621 | 1733937 | 1745320 | 11383 | | V | | | | |
| Apis008626 | ACYPI008626 | GL349628 | 1388475 | 1393860 | 5385 | | I | | | | |
| Apis44656 | ACYPI44656 | GL349661 | 54249 | 60197 | 5948 | | III | | | | |
| Apis34467 | ACYPI34467 | GL349670 | 949.642 | 972.702 | 23.060 | 9036 | III | 1 | 3 | 48653 | |
| Apis34462 | ACYPI34462 | GL349670 | 981.738 | 993.696 | 11.958 | 812 | III | | | | |
| Apis34461 | ACYPI34461 | GL349670 | 994.508 | 998.295 | 3.787 | | III | | | | |
| ApisPPK9 | ACYPI000481 | GL349673 | 586808 | 601946 | 15138 | | II | | | | |
| Apis28482 | ACYPI28482 | GL349690 | 1012848 | 1024902 | 12054 | | I | | | | |

| | | | | | | | | | | | |
|---|---|---|---|---|---|---|---|---|---|---|---|
| Apis33127 | ACYPI33127 | GL349713 | 587886 | 603850 | 15964 | 11366 | V | 2 | 3 | 59667 | Hemipteran Exp. |
| Apis33128 | ACYPI33128 | GL349713 | 615216 | 624114 | 8898 | 13581 | V | | | | Hemipteran Exp. |
| Apis33129 | ACYPI33129 | GL349713 | 637695 | 647553 | 9858 | | V | | | | Hemipteran Exp. |
| Apis31763 | ACYPI31763 | GL349774 | 816369 | 839074 | 22705 | | II | | | | |
| Apis068872 | ACYPI068872 | GL349793 | 74776 | 81537 | 6761 | | V | | | | Hemipteran Exp. |
| Apis33363 | ACYPI33363 | GL349876 | 142215 | 146405 | 4190 | 107 | III | 3 | 2 | 10762 | |
| Apis33364 | ACYPI33364 | GL349876 | 135643 | 142108 | 6465 | | III | | | | |
| Apis27784 | ACYPI27784 | GL349914 | 328827 | 354587 | 25760 | | III | | | | |
| ApisPPK23 | ACYPI008502 | GL349936 | 59372 | 67955 | 8583 | | VI | | | | |
| Apis005555 | ACYPI005555 | GL350004 | 726615 | 731990 | 5375 | | I | | | | |
| Apis35976 | ACYPI35976 | GL350203 | 284957 | 304563 | 19606 | | V | | | | Hemipteran Exp. |
| Apis30092 | ACYPI30092 | GL350222 | 81096 | 96886 | 15790 | | V | | | | Hemipteran Exp. |
| Apis29894 | ACYPI29894 | GL350224 | 160376 | 173912 | 13536 | | V | | | | Hemipteran Exp. |

| **Annotation** | **Original code** | **Scaffold** | **Start** | **Stop** | **Gene length (bp)** | **PPK distance (bp)** | **SF** | **Cluster N°** | **Genes per Cluster** | **Cluster Length (bp)** | **Exp.** |
|---|---|---|---|---|---|---|---|---|---|---|---|
| BgerPPK16 | PSN58104.1 | PYGN01000009.1 | 1998271 | 2016087 | 17816 | | IV | | | | |
| Bger58038 | PSN58038.1 | PYGN01000010.1 | 1741667 | 1757531 | 15864 | | IV | | | | |
| Bger57595 | PSN57595.1 | PYGN01000018.1 | 576714 | 592143 | 15429 | 25510 | III | 1 | 7 | 256112 | Blattella Exp. |
| Bger57602 | PSN57602.1 | PYGN01000018.1 | 617653 | 642388 | 24735 | 5281 | III | | | | Blattella Exp. |
| Bger57599 | PSN57599.1 | PYGN01000018.1 | 647669 | 650117 | 2448 | 5737 | III | | | | Blattella Exp. |
| Bger57593 | PSN57593.1 | PYGN01000018.1 | 655854 | 669085 | 13231 | 11155 | III | | | | Blattella Exp. |
| Bger57594 | PSN57594.1 | PYGN01000018.1 | 680240 | 695330 | 15090 | 23667 | III | | | | Blattella Exp. |
| Bger57566 | PSN57566.1 | PYGN01000018.1 | 718997 | 734401 | 15404 | 89653 | III | | | | Blattella Exp. |
| Bger57569 | PSN57569.1 | PYGN01000018.1 | 824054 | 832826 | 8772 | | III | | | | Blattella Exp. |
| BgerPPK13b | PSN55529.1 | PYGN01000067.1 | 1787630 | 1806193 | 18563 | | I | | | | |
| Bger55271 | PSN55271.1 | PYGN01000073.1 | 12279 | 45406 | 33127 | 12646 | III | 2 | 4 | 105163 | Blattella Exp. |
| Bger55275 | PSN55275.1 | PYGN01000073.1 | 58052 | 71109 | 13057 | 20335 | III | | | | Blattella Exp. |
| Bger55272 | PSN55272.1 | PYGN01000073.1 | 91444 | 99863 | 8419 | 8123 | III | | | | Blattella Exp. |
| Bger55273 | PSN55273.1 | PYGN01000073.1 | 107986 | 117442 | 9456 | | III | | | | Blattella Exp. |
| Bger54949 | PSN54949.1 | PYGN01000079.1 | 1039494 | 1054239 | 14745 | | V | | | | |
| Bger4013 | PSN54013.1 | PYGN01000110.1 | 955848 | 984739 | 28891 | | I | | | | |
| BgerPPK9 | PSN53854.1 | PYGN01000115.1 | 426312 | 433565 | 7253 | | II | | | | |
| Bger52920 | PSN52920.1 | PYGN01000144.1 | 1611158 | 1631777 | 20619 | | I | | | | |
| Bger51114 | PSN51114.1 | PYGN01000211.1 | 1073470 | 1079092 | 5622 | | II | | | | |
| BgerPPK28 | PSN50939.1 | PYGN01000218.1 | 130355 | 168892 | 38537 | | V | | | | |
| Bger50549 | PSN50549.1 | PYGN01000235.1 | 107610 | 133991 | 26381 | | III | | | | |
| Bger49921 | PSN49921.1 | PYGN01000257.1 | 636129 | 642614 | 6485 | | V | | | | |
| BgerPPK17 | PSN44504.1 | PYGN01000545.1 | 591949 | 600773 | 8824 | | VII | | | | |
| BgerPPK13a | PSN43586.1 | PYGN01000599.1 | 828341 | 841959 | 13618 | | I | | | | |
| Bger42805 | PSN42805.1 | PYGN01000652.1 | 606798 | 612089 | 5291 | | V | | | | |
| Bger42712 | PSN42712.1 | PYGN01000657.1 | 458769 | 469517 | 10748 | | IV | | | | |

| | | | | | | | | | | |
|---|---|---|---|---|---|---|---|---|---|---|
| Bger41668 | PSN41668.1 | PYGN01000723.1 | 384615 | 404308 | 19693 | 5218 | I | 3 | 2 (MIXED) | 66637 |
| Bger41670 | PSN41670.1 | PYGN01000723.1 | 409526 | 451252 | 41726 | | II | | | |
| Bger41206 | PSN41206.1 | PYGN01000758.1 | 526269 | 550139 | 23870 | | IV | | | |
| BgerPPK23a | PSN38910.1 | PYGN01000958.1 | 137483 | 153216 | 15733 | 31162 | VI | 4 | 2 | 69736 |
| BgerPPK23b | PSN38909.1 | PYGN01000958.1 | 184378 | 207219 | 22841 | | VI | | | |
| Bger38565 | PSN38565.1 | PYGN01001003.1 | 325122 | 359557 | 34435 | 34527 | V | 5 | 3 | 151151 |
| Bger38575 | PSN38575.1 | PYGN01001003.1 | 270367 | 290595 | 20228 | 31214 | V | | | |
| Bger38576 | PSN38576.1 | PYGN01001003.1 | 208406 | 239153 | 30747 | | V | | | |
| Bger34458 | PSN34458.1 | PYGN01001495.1 | 118207 | 137147 | 18940 | 10109 | V | 6 | 3 | 95645 |
| Bger34456 | PSN34456.1 | PYGN01001495.1 | 147256 | 153442 | 6186 | 45264 | V | | | |
| Bger34460 | PSN34460.1 | PYGN01001495.1 | 198706 | 213852 | 15146 | | V | | | |
| Bger33731 | PSN33731.1 | PYGN01001616.1 | 115786 | 144787 | 29001 | | III | | | |
| Bger32397 | PSN32397.1 | PYGN01001870.1 | 46959 | 51748 | 4789 | | I | | | |
| Bger31707 | PSN31707.1 | PYGN01002044.1 | 66878 | 103601 | 36723 | | II | | | |
| Bger30177 | PSN30177.1 | PYGN01002624.1 | 110648 | 146438 | 35790 | | III | | | |

| **Annotation** | **Original code** | **Scaffold** | **Start** | **Stop** | **Gene length (bp)** | **PPK distance (bp)** | **SF** | **Cluster N°** | **Genes per Cluster** | **Cluster Length (bp)** | **Exp.** |
|---|---|---|---|---|---|---|---|---|---|---|---|
| Bimp003493499 | XP_003493499.1 | NT_177527.1 | 33749 | 43166 | 9417 | | IV | | | | |
| Bimp033179584 | XP_033179584.1 | NT_177527.1 | 11131 | 21300 | 10169 | 12449 | IV | 1 | 2 | 32035 | |
| BimpPPK16 | XP_024224189.1 | NT_176861.1 | 1309322 | 1312516 | 3194 | | IV | | | | |
| BimpPPK23 | XP_012240062.1 | NT_176683.1 | 857853 | 859593 | 1740 | | VI | | | | |
| Bimp024222992 | XP_024222992.1 | NT_176736.1 | 2723995 | 2728077 | 4082 | | IV | | | | |
| Bimp012242038 | XP_012242038.1 | NT_176825.1 | 766576 | 768694 | 2118 | | II | | | | |
| Bimp012246752 | XP_012246752.1 | NT_176468.1 | 942207 | 945953 | 3746 | | I | | | | |
| BimpPPK28 | XP_012237817.1 | NT_176570.1 | 733740 | 738925 | 5185 | | V | | | | |
| Bimp012239254 | XP_012239254.1 | NT_176644.1 | 1159992 | 1174269 | 14277 | | IV | | | | |

| **Annotation** | **Original code** | **Scaffold** | **Start** | **Stop** | **Gene length (bp)** | **PPK distance (bp)** | **SF** | **Cluster N°** | **Genes per Cluster** | **Cluster Length (bp)** | **Exp.** |
|---|---|---|---|---|---|---|---|---|---|---|---|
| Bmor012544414.1 | XP_012544414.1 | NW_004581681.1 | 595890 | 601417 | 5527 | 120 | IV | 1 | 2 | 23628 | |
| Bmor012544908.1 | XP_012544908.1 | NW_004581681.1 | 601537 | 619518 | 17981 | | IV | | | | |
| Bmor012552731.1 | XP_012552731.1 | NW_004581690.1 | 2271152 | 2290777 | 19625 | | V | | | | |
| Bmor012553296.1 | XP_012553296.1 | NW_004581698.1 | 1265877 | 1284309 | 18432 | | I | | | | |
| BmorPPK17 | XP_012545187.1 | NW_004581729.1 | 1510553 | 1519793 | 9240 | | VII | | | | |
| Bmor012545602.1 | XP_012545602.1 | NW_004581742.1 | 740117 | 811237 | 71120 | | IV | | | | |
| Bmor012545987.1 | XP_012545987.1 | NW_004581753.1 | 346169 | 356083 | 9914 | 131920 | IV | | | | |
| Bmor012545990.1 | XP_012545990.1 | NW_004581753.1 | 488003 | 497344 | 9341 | | IV | | | | |
| BmorPPK25 | XP_012546079.1 | NW_004581756.1 | 1141812 | 1151989 | 10177 | 387450 | II | | | | |
| Bmor012546099.1 | XP_012546099.1 | NW_004581756.1 | 739997 | 754362 | 14365 | | II | | | | |
| BmorPPK28 | XP_012548124.1 | NW_004582011.1 | 3573272 | 3581231 | 7959 | | V | | | | |
| Bmor004929670.1 | XP_004929670.1 | NW_004582014.1 | 4360833 | 4368331 | 7498 | | II | | | | |
| BmorPPK16 | XP_012548978.1 | NW_004582015.1 | 3825347 | 3846390 | 21043 | | IV | | | | |
| BmorPPK9 | XP_012550136.1 | NW_004582021.1 | 5958914 | 5965670 | 6756 | | II | | | | |
| Bmor012550654.1 | XP_012550654.1 | NW_004582024.1 | 4482765 | 4493679 | 10914 | | V | | | | |
| BmorPPK13b | XP_012550884.1 | NW_004582026.1 | 1032748 | 1037512 | 4764 | 17664 | I | 2 | 3 (MIXED) | 43437 | |
| Bmor004932266.1 | XP_004932266.1 | NW_004582026.1 | 1055176 | 1066058 | 10882 | 5240 | IV | | | | |
| BmorPPK13a | XP_004932267.1 | NW_004582026.1 | 1071298 | 1076185 | 4887 | | I | | | | |

| **Annotation** | **Original code** | **Scaffold** | **Start** | **Stop** | **Gene length** | **PPK distance** | **SF** | **Cluster** | **Genes per** | **Cluster** | **Exp.** |
|---|---|---|---|---|---|---|---|---|---|---|---|

| | | | | | (bp) | (bp) | | N° | Cluster | Length (bp) | |
|---|---|---|---|---|---|---|---|---|---|---|---|
| CfloPPK28 | XP_011269762.1 | NW_020229681.1 | 544757 | 549374 | 4617 | | V | | | | |
| CfloPPK16c | XP_025270843.1 | NW_020229867.1 | 23418 | 29717 | 6299 | 6072 | IV | 1 | 3 | 43862 | |
| CfloPPK16b | XP_025270842.1 | NW_020229867.1 | 41767 | 47104 | 5337 | 12050 | IV | | | | |
| CfloPPK16a | XP_025270836.1 | NW_020229867.1 | 3242 | 17346 | 14104 | | IV | | | | |
| Cflo025269475 | XP_025269475.1 | NW_020229838.1 | 3077241 | 3080606 | 3365 | | IV | | | | |
| Cflo025266757 | XP_025266757.1 | NW_020229506.1 | 33274 | 53899 | 20625 | | IV | | | | |
| Cflo025266050 | XP_025266050.1 | NW_020229437.1 | 765232 | 767782 | 2550 | | IV | | | | |
| Cflo025263766 | XP_025263766.1 | NW_020229325.1 | 2520400 | 2531915 | 11515 | | III | | | | |
| **Annotation** | **Original code** | **Scaffold** | **Start** | **Stop** | **Gene length (bp)** | **PPK distance (bp)** | **SF** | **Cluster N°** | **Genes per Cluster** | **Cluster Length (bp)** | **Exp.** |
| Clec003170 | CLEC003170 | KK244455 | 10322651 | 10327891 | 5240 | 13373672 | I | | | | |
| ClecPPK28 | CLEC000413 | KK244445 | 23701563 | 23704963 | 3400 | 3360945 | V | | | | |
| Clec000454 | CLEC000454 | KK244445 | 27065908 | 27072599 | 6691 | 25134 | I | 1 | 3(MIXED) | 43311 | |
| ClecPPK9b | CLEC000458 | KK244445 | 27097733 | 27100265 | 2532 | 6586 | II | | | | |
| ClecPPK9a | CLEC000460 | KK244445 | 27106851 | 27109219 | 2368 | | II | | | | |
| Clec004429 | CLEC004429 | KK244457 | 2939963 | 2942413 | 2450 | | II | | | | |
| Clec004707 | CLEC004707 | KK244468 | 3573470 | 3574277 | 807 | 4515454 | III | | | | |
| Clec004780 | CLEC004780 | KK244468 | 8089731 | 8104841 | 15110 | | III | | | | |
| Clec006346 | CLEC006346 | KK244482 | 4154388 | 4157361 | 2973 | | V | | | | |
| ClecPPK23 | CLEC012452 | KK244534 | 757342 | 758543 | 1201 | | VI | | | | |
| **Annotation** | **Original code** | **Scaffold** | **Start** | **Stop** | **Gene length (bp)** | **PPK distance (bp)** | **SF** | **Cluster N°** | **Genes per Cluster** | **Cluster Length (bp)** | **Exp.** |
| CquiPPK25 | CPIJ000103 | supercont3.1 | 2259438 | 2260999 | 1561 | | II | | | | |
| Cqui017961 | CPIJ017961 | supercont3.1053 | 50795 | 52966 | 2171 | | V | | | | Culicidae Exp. |
| Cqui005908 | CPIJ005908 | supercont3.111 | 841784 | 843914 | 2130 | 1530 | V | 1 | 2 | 5513 | Culicidae Exp. |
| Cqui005909 | CPIJ005909 | supercont3.111 | 845444 | 847297 | 1853 | | V | | | | Culicidae Exp. |
| CquiPPK10 | CPIJ006134 | supercont3.117 | 887180 | 906813 | 19633 | | II | | | | |
| CquiPPK301 | CPIJ007315 | supercont3.142 | 440666 | 448594 | 7928 | | V | | | | |
| CquiPPK6a | CPIJ018767 | supercont3.1463 | 82863 | 89802 | 6939 | | IV | | | | |
| CquiPPK13 | CPIJ019036 | supercont3.1484 | 64489 | 71578 | 7089 | | I | | | | |
| Cqui007799 | CPIJ007799 | supercont3.160 | 173477 | 175257 | 1780 | 122 | V | **2** | 13 | 27168 | Culicidae Exp. |
| Cqui007800 | CPIJ007800 | supercont3.160 | 175379 | 176918 | 1539 | 461 | V | | | | Culicidae Exp. |
| Cqui007801 | CPIJ007801 | supercont3.160 | 177379 | 179147 | 1768 | 343 | V | | | | Culicidae Exp. |
| Cqui007802 | CPIJ007802 | supercont3.160 | 179490 | 183124 | 3634 | 161 | V | | | | Culicidae Exp. |
| Cqui007803 | CPIJ007803 | supercont3.160 | 183285 | 185049 | 1764 | 134 | V | | | | Culicidae Exp. |
| Cqui007804 | CPIJ007804 | supercont3.160 | 185183 | 186789 | 1606 | 496 | V | | | | Culicidae Exp. |
| Cqui007805 | CPIJ007805 | supercont3.160 | 187285 | 188688 | 1403 | 158 | V | | | | Culicidae Exp. |
| Cqui007806 | CPIJ007806 | supercont3.160 | 188846 | 190702 | 1856 | 205 | V | | | | Culicidae |

| | | | | | | | | | | | |
|---|---|---|---|---|---|---|---|---|---|---|---|
| | | | | | | | | | | | Exp. |
| Cqui007807 | CPIJ007807 | supercont3.160 | 190907 | 192670 | 1763 | 290 | V | | | | Culicidae Exp. |
| Cqui007808 | CPIJ007808 | supercont3.160 | 192960 | 194697 | 1737 | 243 | V | | | | Culicidae Exp. |
| Cqui007809 | CPIJ007809 | supercont3.160 | 194940 | 196769 | 1829 | 189 | V | | | | Culicidae Exp. |
| Cqui007810 | CPIJ007810 | supercont3.160 | 196958 | 198678 | 1720 | 162 | V | | | | Culicidae Exp. |
| Cqui007811 | CPIJ007811 | supercont3.160 | 198840 | 200645 | 1805 | | V | | | | Culicidae Exp. |
| CquiPPK9 | CPIJ007762 | supercont3.162 | 360057 | 361190 | 1133 | | II | | | | |
| Cqui008773 | CPIJ008773 | supercont3.205 | 365171 | 366604 | 1433 | | II | | | | |
| CquiPPK15 | CPIJ001728 | supercont3.25 | 678991 | 680613 | 1622 | | I | | | | |
| Cqui010135 | CPIJ010135 | supercont3.251 | 372882 | 374400 | 1518 | 3824 | V | 3 | 2 | 7149 | Culicidae Exp. |
| Cqui010136 | CPIJ010136 | supercont3.251 | 378224 | 380031 | 1807 | | V | | | | Culicidae Exp. |
| Cqui002355 | CPIJ002355 | supercont3.33 | 187365 | 207137 | 19772 | | IV | | | | |
| CquiPPK16 | CPIJ012002 | supercont3.346 | 218799 | 222850 | 4051 | | IV | | | | |
| Cqui012543 | CPIJ012543 | supercont3.432 | 321861 | 323633 | 1772 | 1762 | V | 4 | 5 | 14829 | Culicidae Exp. |
| Cqui012544 | CPIJ012544 | supercont3.432 | 325395 | 327014 | 1619 | 820 | V | | | | Culicidae Exp. |
| Cqui012545 | CPIJ012545 | supercont3.432 | 327834 | 334390 | 6556 | 609 | V | | | | Culicidae Exp. |
| Cqui012546 | CPIJ012546 | supercont3.432 | 334999 | 336690 | 1691 | | V | | | | Culicidae Exp. |
| Cqui003410 | CPIJ003410 | supercont3.44 | 1107139 | 1109249 | 2110 | 423 | V | 5 | 2 | 4447 | Culicidae Exp. |
| Cqui003411 | CPIJ003411 | supercont3.44 | 1109672 | 1111586 | 1914 | | V | | | | Culicidae Exp. |
| CquiPPK3 | CPIJ003913 | supercont3.61 | 108564 | 124123 | 15559 | | II | | | | |
| CquiPPK31 | CPIJ015546 | supercont3.650 | 15232 | 21113 | 5881 | | I | | | | |
| Cqui005124 | CPIJ005124 | supercont3.84 | 314058 | 315896 | 1838 | 1215 | V | 6 | 7 | 41008 | Culicidae Exp. |
| Cqui005125 | CPIJ005125 | supercont3.84 | 317111 | 318724 | 1613 | 16659 | V | | | | Culicidae Exp. |
| Cqui005126 | CPIJ005126 | supercont3.84 | 335383 | 337167 | 1784 | 132 | V | | | | Culicidae Exp. |
| Cqui005127 | CPIJ005127 | supercont3.84 | 337299 | 339485 | 2186 | 559 | V | | | | Culicidae Exp. |
| Cqui005128 | CPIJ005128 | supercont3.84 | 340044 | 343186 | 3142 | 588 | V | | | | Culicidae Exp. |
| Cqui005129 | CPIJ005129 | supercont3.84 | 343774 | 345451 | 1677 | 1141 | V | | | | Culicidae Exp. |
| Cqui005131 | CPIJ005131 | supercont3.84 | 346592 | 355066 | 8474 | | V | | | | Culicidae Exp. |
| CquiPPK6b | CPIJ005031 | supercont3.89 | 342530 | 348000 | 5470 | | IV | | | | |
| CquiPPK17 | CPIJ000851 | supercont3.9 | 267433 | 270786 | 3353 | | VII | | | | |
| CquiPPK23a | CPIJ017580 | supercont3.959 | 113250 | 115112 | 1862 | 9823 | VI | 7 | 2 | 13662 | |
| CquiPPK23b | CPIJ017584 | supercont3.959 | 124935 | 126912 | 1977 | | VI | | | | |
| Cqui005430 | CPIJ005430 | supercont3.97 | 85324 | 101224 | 15900 | | IV | | | | |

| Annotation | Original code | Scaffold | Start | Stop | Gene length (bp) | PPK distance (bp) | SF | Cluster N° | Genes per Cluster | Cluster Length (bp) | Exp. |
|---|---|---|---|---|---|---|---|---|---|---|---|
| DmelPPK7 | CG9499 | 2L | 6352324 | 6354331 | 2007 | 625 | III | 1 | 2 | 4348 | |
| DmelPPK14 | CG9501 | 2L | 6354956 | 6356672 | 1716 | 3408685 | III | | | | |
| DmelPPK18 | CG44152 | 2L | 9765357 | 9770996 | 5639 | 159 | IV | 2 | 3 | 9673 | |
| DmelPPK11 | CG34058 | 2L | 9771155 | 9773042 | 1887 | | IV | | | | |
| DmelPPK16 | CG34059 | 2L | 9771155 | 9775030 | 3875 | 675104 | IV | | | | |
| DmelPPK10 | CG34042 | 2L | 10450134 | 10452201 | 2067 | 3926620 | II | | | | |
| DmelPPK1 | CG3478 | 2L | 14378821 | 14381134 | 2313 | 2377216 | V | | | | |
| DmelPPK17 | CG13278 | 2L | 16758350 | 16760113 | 1763 | 4326787 | VII | | | | |
| DmelPPK13 | CG33508 | 2L | 21086900 | 21088730 | 1830 | | I | | | | |
| DmelPPK25 | CG33349 | 2R | 6950391 | 6952158 | 1767 | 9701047 | II | | | | |
| DmelPPK4 | CG8178 | 2R | 16653205 | 16655046 | 1841 | 3590725 | IV | | | | |
| DmelPPK6 | CG11209 | 2R | 20245771 | 20247507 | 1736 | 1716666 | IV | | | | |
| DmelPPK9 | CG34369 | 2R | 21964173 | 21970664 | 6491 | 456815 | II | | | | |
| DmelPPK12 | CG10972 | 2R | 22427479 | 22429584 | 2105 | 1109047 | V | | | | |
| DmelPPK3 | CG30181 | 2R | 23538631 | 23540773 | 2142 | 512256 | II | | | | |
| DmelPPK29 | CG13568 | 2R | 24053029 | 24054873 | 1844 | | I | | | | |
| DmelPPK26 | CG8546 | 3L | 7456189 | 7458353 | 2164 | 3701100 | V | | | | |
| DmelPPK27 | CG10858 | 3L | 3753693 | 3755089 | 1396 | 14076307 | IV | | | | |
| DmelPPK5 | CG33289 | 3L | 21534660 | 21536861 | 2201 | | V | | | | |
| DmelPPK2 | CG1058 | 3R | 4646064 | 4648138 | 2074 | 20211102 | V | | | | |
| DmelPPK22 | CG31105 | 3R | 24859240 | 24859711 | 471 | | IV | | | | |
| DmelPPK15 | CG14239 | 3R | 26401909 | 26403656 | 1747 | 759468 | I | | | | |
| DmelPPK31 | CG31065 | 3R | 27163124 | 27165131 | 2007 | 2499648 | I | | | | |
| DmelPPK21 | CG12048 | 3R | 29664779 | 29666783 | 2004 | 31 | III | 3 | 4 | 9444 | |
| DmelPPK20 | CG7577 | 3R | 29666814 | 29668877 | 2063 | 281 | III | | | | |
| DmelPPK30 | CG18110 | 3R | 29669158 | 29670872 | 1714 | 1584 | III | | | | |
| DmelPPK19 | CG18287 | 3R | 29672456 | 29674223 | 1767 | 1591507 | III | | | | |
| DmelPPK24 | CG15555 | 3R | 31265730 | 31268511 | 2781 | | IV | | | | |
| DmelPPK8 | CG32792 | X | 3569504 | 3571800 | 2296 | 13234507 | V | | | | |
| DmelPPK28 | CG4805 | X | 16806307 | 16808385 | 2078 | 758860 | V | | | | |
| DmelPPK23 | CG8527 | X | 17567245 | 17569135 | 1890 | | VI | | | | |

| Annotation | Original code | Scaffold | Start | Stop | Gene length (bp) | PPK distance (bp) | SF | Cluster N° | Genes per Cluster | Cluster Length (bp) | Exp. |
|---|---|---|---|---|---|---|---|---|---|---|---|
| DplexPPK28 | DPOGS206892 | DPSCF300001 | 1828834 | 1836130 | 7296 | | V | | | | |
| Dplex210747 | DPOGS210747 | DPSCF300013 | 549109 | 556515 | 7406 | | I | | | | |
| Dplex214762 | DPOGS214762 | DPSCF300022 | 1335996 | 1339533 | 3537 | | IV | | | | |
| Dplex215856 | DPOGS215856 | DPSCF300029 | 1147465 | 1148253 | 788 | | I | | | | |
| Dplex204183 | DPOGS204183 | DPSCF300034 | 218638 | 224272 | 5634 | | IV] | | | | |
| DplexPPK6 | DPOGS214642 | DPSCF300050 | 664207 | 667491 | 3284 | | IV | | | | |
| DplexPPK16 | DPOGS200906 | DPSCF300066 | 133826 | 155197 | 21371 | | IV | | | | |
| DplexPPK23 | DPOGS215595 | DPSCF300097 | 228774 | 232058 | 3284 | | VI | | | | |
| Dplex216041 | DPOGS216041 | DPSCF300067 | 298137 | 318504 | 20367 | 1139 | V | 1 | 2 | 25686 | |
| Dplex216040 | DPOGS216040 | DPSCF300067 | 319643 | 323823 | 4180 | | V | | | | |
| DplexPPK9 | DPOGS210227 | DPSCF300196 | 449782 | 452935 | 3153 | | II | | | | |

| | | | | | | | | | | | |
|---|---|---|---|---|---|---|---|---|---|---|---|
| Dplex204735 | DPOGS204735 | DPSCF300231 | 648273 | 649850 | 1577 | 74492 | IV | 2 | 2 | 78878 | |
| Dplex204741 | DPOGS204741 | DPSCF300231 | 570972 | 573781 | 2809 | | IV | | | | |
| Dplex209994 | DPOGS209994 | DPSCF300247 | 9596 | 15256 | 5660 | | IV | | | | |
| DplexPPK13b | DPOGS202727 | DPSCF300284 | 199743 | 208011 | 8268 | | I | | | | |
| Dplex211506 | DPOGS211506 | DPSCF300354 | 149982 | 152329 | 2347 | | II | | | | |
| DplexPPK25 | DPOGS207910 | DPSCF300478 | 2913 | 5705 | 2792 | | II | | | | |
| DplexPPK13a | DPOGS203427 | DPSCF300548 | 18946 | 21302 | 2356 | | I | | | | |
| **Annotation** | **Original code** | **Scaffold** | **Start** | **Stop** | **Gene length (bp)** | **PPK distance (bp)** | **SF** | **Cluster N°** | **Genes per Cluster** | **Cluster Length (bp)** | **Exp.** |
| DponPPK23 | ENN81927 | Seq_1101744 | 95576 | 104098 | 8522 | | VI | | | | |
| Dpon81697 | ENN81697 | Seq_1101817 | 496555 | 501212 | 4657 | 3092 | V | 1 | 2 | 11942 | Coleopteran Exp. |
| Dpon81698 | ENN81698 | Seq_1101817 | 504304 | 508497 | 4193 | | V | | | | Coleopteran Exp. |
| DponPPK13 | ENN81481 | Seq_1101838 | 941284 | 943431 | 2147 | | I | | | | |
| Dpon81350 | ENN81350 | Seq_1101853 | 67517 | 72722 | 5205 | | II | | | | |
| DponPPK16a | ENN77435 | Seq_1102694 | 3323863 | 3325196 | 1333 | 409 | IV | 2 | 3 | 20825 | |
| DponPPK16b | ENN77436 | Seq_1102694 | 3325605 | 3333384 | 7779 | 8395 | IV | | | | |
| DponPPK16c | ENN77437 | Seq_1102694 | 3341779 | 3344688 | 2909 | | IV | | | | |
| Dpon76926 | ENN76926 | Seq_1102712 | 682254 | 690526 | 8272 | | V | | | | Coleopteran Exp. |
| DponPPK9 | ENN75495 | Seq_1102760 | 2344197 | 2345325 | 1128 | | II | | | | |
| DponPPK3 | ENN73452 | Seq_1102910 | 364497 | 370039 | 5542 | | II | | | | |
| DponPPK28 | ENN71763 | Seq_1103002 | 171845 | 175886 | 4041 | | V | | | | |
| Dpon71305 | ENN71305 | Seq_1103023 | 3994015 | 3998392 | 4377 | 1929269 | V | | | | Coleopteran Exp. |
| DponPPK17 | ENN71156 | Seq_1103023 | 2059370 | 2064746 | 5376 | | VII | | | | |
| **Annotation** | **Original code** | **Scaffold** | **Start** | **Stop** | **Gene length (bp)** | **PPK distance (bp)** | **SF** | **Cluster N°** | **Genes per Cluster** | **Cluster Length (bp)** | **Exp.** |
| GmorPPK29 | GMOY001021 | scf7180000640787 | 313761 | 321285 | 7524 | | I | | | | |
| GmorPPK7 | GMOY001228 | scf7180000641081 | 7897 | 10906 | 3009 | | III | | | | |
| GmorPPK2 | GMOY002353 | scf7180000643466 | 26875 | 31160 | 4285 | | V | | | | |
| GmorPPK26 | GMOY004243 | scf7180000648009 | 59928 | 62572 | 2644 | | V | | | | |
| GmorPPK1b | GMOY004489 | scf7180000648117 | 10731 | 11392 | 661 | | V | | | | |
| GmorPPK10b | GMOY004892 | scf7180000648322 | 11141 | 16527 | 5386 | | II | | | | |
| GmorPPK17 | GMOY005158 | scf7180000649450 | 62216 | 66034 | 3818 | | VII | | | | |
| GmorPPK28 | GMOY005504 | scf7180000648623 | 48678 | 51784 | 3106 | | V | | | | |
| GmorPPK1c | GMOY006517 | scf7180000649080 | 73775 | 77259 | 3484 | | V | | | | |
| GmorPPK23 | GMOY007016 | scf7180000649907 | 40475 | 43304 | 2829 | | VI | | | | |
| GmorPPK10a | GMOY007470 | scf718000065041 | 195682 | 199624 | 3942 | | II | | | | |

| | | | | | | | |
|---|---|---|---|---|---|---|---|
| | | 1 | | | | | |
| GmorPPK22 | GMOY008422 | scf7180000651278 | 7467 | 14150 | 6683 | | IV |
| GmorPPK6 | GMOY009113 | scf7180000651771 | 568398 | 570756 | 2358 | | IV |
| GmorPPK1a | GMOY010902 | scf7180000652158 | 176418 | 177079 | 661 | | V |
| GmorPPK15 | GMOY011877 | scf7180000652170 | 22568058 | 22579117 | 11059 | | I |

| Annotation | Original code | Scaffold | Start | Stop | Gene length (bp) | PPK distance (bp) | SF | Cluster N° | Genes per Cluster | Cluster Length (bp) | Exp. |
|---|---|---|---|---|---|---|---|---|---|---|---|
| Ldec008039 | LDEC008039 | NW_019289547.1 | 774211 | 778332 | 4121 | | IV | | | | |
| Ldec011575 | LDEC011575 | NW_019291145.1 | 51259 | 68157 | 16898 | | V | | | | Coleopteran Exp. |
| Ldec017033 | LDEC017033 | NW_019292422.1 | 2619 | 22174 | 19555 | | V | | | | Coleopteran Exp. |
| Ldec017036 | LDEC017036 | NW_019291856.1 | 4342 | 10310 | 5968 | 5734 | V | 1 | 2 | 18765 | Coleopteran Exp. |
| Ldec017037 | Ldec017037 | NW_019291856.1 | 16044 | 23107 | 7063 | | V | | | | Coleopteran Exp. |
| Ldec019895 | LDEC019895 | NW_019291092.1 | 43513 | 52761 | 9248 | 16034 | V | 2 | 3 | 40420 | Coleopteran Exp. |
| Ldec024423 | LDEC024423 | NW_019291092.1 | 68795 | 72870 | 4075 | 150 | V | | | | Coleopteran Exp. |
| Ldec019897 | LDEC019897 | NW_019291092.1 | 73020 | 83933 | 10913 | | V | | | | Coleopteran Exp. |
| LdecPPK3 | LDEC019444 | NW_019292019.1 | 47075 | 59488 | 12413 | | II | | | | |
| LdecPPK9 | LDEC010412 | NW_019289516.1 | 207584 | 214521 | 6937 | | II | | | | |

| Annotation | Original code | Scaffold | Start | Stop | Gene length (bp) | PPK distance (bp) | SF | Cluster N° | Genes per Cluster | Cluster Length (bp) | Exp. |
|---|---|---|---|---|---|---|---|---|---|---|---|
| Lmig10489.2 | JAMg_model_10489.2 | scaffold185071 | 292992 | 382317 | 89325 | | IV | | | | |
| Lmig11504.1 | JAMg_model_11504.1 | scaffold145746 | 71787 | 102526 | 30739 | | I | | | | |
| Lmig1169.1 | JAMg_model_1169.1 | scaffold145746 | 166286 | 227161 | 60785 | | II | | | | |
| Lmig12479.1 | JAMg_model_12479.1 | scaffold74895 | 158969 | 212556 | 53587 | | III | | | | |
| Lmig12972.1 | JAMg_model_12972.1 | scaffold13984 | 2204122 | 2261231 | 57109 | | IV | | | | |
| Lmig14370 | JAMg_model_14370.1 | scaffold158451 | 16981 | 140971 | 123990 | | I | | | | |
| Lmig14820.1 | JAMg_model_14820.1 | scaffold249954 | 10440 | 134369 | 123929 | | V | | | | |
| Lmig1583.1 | JAMg_model_1583.1 | scaffold155404 | 315823 | 353517 | 37694 | | III | | | | |
| Lmig17317.1 | JAMg_model_17317.1 | scaffold4844 | 577920 | 827567 | 249647 | | V | | | | |
| Lmig18227.1 | JAMg_model_18227.1 | scaffold166805 | 180332 | 217868 | 37536 | | IV | | | | |
| Lmig18332 | JAMg_model_18332.1 | scaffold26869 | 57498 | 143419 | 85921 | | IV | | | | |
| Lmig2248.1 | JAMg_model_2248.1 | scaffold145426 | 323111 | 533760 | 210649 | | V | | | | |
| Lmig8491.1 | JAMg_model_8491.1 | scaffold21927 | 471755 | 583429 | 111674 | | V | | | | |
| Lmig9685.1 | JAMg_model_9685.1 | scaffold200846 | 130453 | 264976 | 134523 | | V | | | | |
| LmigPPK13 | JAMg_model_9564.3 | scaffold242481 | 281203 | 370092 | 88889 | | I | | | | |
| LmigPPK16 | JAMg_model_7461.1 | scaffold141449 | 50793 | 169237 | 118444 | | IV | | | | |
| LmigPPK17 | JAMg_model_492.1 | scaffold66553 | 710503 | 872962 | 162459 | | V | | | | |

| | | | | | | | | | | | |
|---|---|---|---|---|---|---|---|---|---|---|---|
| LmigPPK28a | JAMg_model_1998.1 | scaffold95074 | 21457 | 63822 | 42362 | | V | | | | |
| LmigPPK28b | JAMg_model_20185.1 | scaffold127719 | 2178818 | 2189814 | 10996 | | V | | | | |
| **Annotation** | **Original code** | **Scaffold** | **Start** | **Stop** | **Gene length (bp)** | **PPK distance (bp)** | **SF** | **Cluster N°** | **Genes per Cluster** | **Cluster Length (bp)** | **Exp.** |
| MdomPPK1 | XP_005174914.1 | NW_004754940.1 | 2111288 | 2115906 | 4618 | | V | | | | |
| MdomPPK14a | XP_019891003.1 | NW_004757230.1 | 548 | 1314 | 766 | | III | | | | |
| MdomPPK10a | XP_005175357.1 | NW_004758386.1 | 340 | 5481 | 5141 | | II | | | | |
| MdomPPK6 | XP_005175804.1 | NW_004763385.1 | 17218 | 19296 | 2078 | | IV | | | | |
| MdomPPK17 | XP_005176331.1 | NW_004764473.1 | 9076 | 22658 | 13582 | | VII | | | | |
| MdomPPK2i | XP_005176877.1 | NW_004764503.1 | 215617 | 217347 | 1730 | 857 | V | 1 | 4 | 21485 | |
| MdomPPK2j | XP_005176878.1 | NW_004764503.1 | 218204 | 223864 | 5660 | 3290 | V | | | | |
| MdomPPK2k | XP_005176879.1 | NW_004764503.1 | 227154 | 230857 | 3703 | 4526 | V | | | | |
| MdomPPK2h | XP_005176880.1 | NW_004764503.1 | 235383 | 237102 | 1719 | | V | | | | |
| MdomPPK2a | XP_019890548.1 | NW_004764576.1 | 328835 | 330594 | 1759 | 7940 | V | 2 | 5 | 43245 | |
| MdomPPK2e | XP_005178343.1 | NW_004764576.1 | 338534 | 340252 | 1718 | 5793 | V | | | | |
| MdomPPK2g | XP_011290422.1 | NW_004764576.1 | 346045 | 349952 | 3907 | 9357 | V | | | | |
| MdomPPK2c | XP_019890549.1 | NW_004764576.1 | 359309 | 365234 | 5925 | 3115 | V | | | | |
| MdomPPK2d | XP_019890552.1 | NW_004764576.1 | 368349 | 372080 | 3731 | | V | | | | |
| MdomPPK4 | XP_005178822.1 | NW_004764606.1 | 435103 | 438499 | 3396 | | IV | | | | |
| MdomPPK12a | XP_019890716.1 | NW_004764610.1 | 88096 | 94341 | 6245 | 11869 | V | 3 | 2 | 11869 | |
| MdomPPK12b | XP_019890715.1 | NW_004764610.1 | 82472 | 86974 | 4502 | | V | | | | |
| MdomPPK31 | XP_005178919.2 | NW_004764613.1 | 367058 | 372551 | 5493 | | I | | | | |
| MdomPPK27a | XP_005178987.1 | NW_004764616.1 | 378876 | 384421 | 5545 | 1719 | IV | 4 | 2 | 56166 | |
| MdomPPK27b | XP_019890758.1 | NW_004764616.1 | 382702 | 384421 | 1719 | | IV | | | | |
| MdomPPK3 | XP_005179130.2 | NW_004764620.1 | 1270833 | 1278674 | 7841 | | II | | | | |
| MdomPPK5b | XP_019891014.1 | NW_004764670.1 | 419559 | 426023 | 6464 | 6144 | V | 5 | 3 | 582337 | |
| MdomPPK5c | XP_011290988.2 | NW_004764670.1 | 432167 | 435042 | 2875 | 8113 | V | | | 15396 | |
| MdomPPK5a | XP_005179825.2 | NW_004764670.1 | 443155 | 447563 | 4408 | | V | | | | |
| MdomPPK23 | XP_005180321.1 | NW_004764723.1 | 329695 | 337055 | 7360 | | VI | | | | |
| MdomPPK26 | XP_011291290.1 | NW_004764744.1 | 992678 | 1001896 | 9218 | | V | | | | |
| Mdom019891600.1 | XP_019891600.1 | NW_004764826.1 | 88668 | 93780 | 5112 | | II | | | | |
| MdomPPK29 | XP_005181951.1 | NW_004764881.1 | 91656 | 97116 | 5460 | | I | | | | |
| MdomPPK25 | XP_019891859.1 | NW_004764906.1 | 950167 | 954135 | 3968 | | II | | | | |
| Mdom011292215.1 | XP_011292215.1 | NW_004764967.1 | 101719 | 110063 | 8344 | | I | | | | |
| MdomPPK8c | XP_019892519.1 | NW_004765155.1 | 2323 | 13660 | 11337 | | V | | | | |
| MdomPPK13 | XP_005184244.2 | NW_004765167.1 | 72549 | 77789 | 5240 | | I | | | | |
| MdomPPK2b | XP_011292906.2 | NW_004765198.1 | 120581 | 120656 | 75 | | V | | | | |
| MdomPPK9 | XP_011293257.1 | NW_004765349.1 | 719137 | 722926 | 3789 | | II | | | | |
| MdomPPK19b | XP_019893823.1 | NW_004765905.1 | 342 | 3776 | 3434 | 3887 | III | 6 | 5 | 3887 | |
| MdomPPK19c | XP_019893824.1 | NW_004765905.1 | 7663 | 14525 | 6862 | 18478 | III | | | 18478 | |
| MdomPPK19d | XP_011294238.1 | NW_004765905.1 | 33003 | 42419 | 9416 | 4001 | III | | | 4001 | |
| MdomPPK19f | XP_019893825.1 | NW_004765905.1 | 46420 | 52028 | 5608 | 11571 | III | | | 11571 | |
| MdomPPK19e | XP_005187753.2 | NW_004765905.1 | 63599 | 73832 | 10233 | | III | | | | |
| MdomPPK10b | XP_005187879.1 | NW_004765945.1 | 53354 | 67199 | 13845 | | II | | | | |
| MdomPPK22 | XP_019893922.1 | NW_004766008.1 | 47277 | 74209 | 26932 | 9566 | IV | 7 | 2 | 9566 | |
| MdomPPK24 | XP_019893923.1 | NW_004766008.1 | 83775 | 90019 | 6244 | | IV | | | | |

| | | | | | | | | | | |
|---|---|---|---|---|---|---|---|---|---|---|
| MdomPPK2f | XP_011294524.1 | NW_004766068.1 | 58282 | 60037 | 1755 | | V | | | |
| MdomPPK15a | XP_019894349.1 | NW_004766407.1 | 81912 | 82898 | 986 | | I | | | |
| MdomPPK21a | XP_019894437.1 | NW_004766537.1 | 27072 | 28284 | 1212 | 3632 | III | 8 | 5 | 3632 |
| MdomPPK21b | XP_019894438.1 | NW_004766537.1 | 31916 | 43381 | 11465 | 5500 | III | | | 5500 |
| MdomPPK20 | XP_019894440.1 | NW_004766537.1 | 48881 | 59116 | 10235 | 6294 | III | | | 6294 |
| MdomPPK30 | XP_019894441.1 | NW_004766537.1 | 65410 | 70904 | 5494 | 12655 | III | | | 12655 |
| MdomPPK19a | XP_019894439.1 | NW_004766537.1 | 83559 | 91171 | 7612 | | III | | | |
| MdomPPK8a | XP_019894556.1 | NW_004767050.1 | 1699 | 9081 | 7382 | 5064 | V | 9 | 4 | 5064 |
| MdomPPK8b | XP_019894557.1 | NW_004767050.1 | 14145 | 19040 | 4895 | 906 | V | | | 906 |
| MdomPPK28b | XP_005189737.1 | NW_004767050.1 | 19946 | 22081 | 2135 | 2033 | V | | | 2033 |
| MdomPPK28a | XP_005189738.2 | NW_004767050.1 | 24114 | 30914 | 6800 | | V | | | |
| Mdom019894840.1 | XP_019894840.1 | NW_004768977.1 | 8181 | 10289 | 2108 | | III | | | |
| MdomPPK14b | XP_005190919.1 | NW_004769580.1 | 367 | 1476 | 1109 | 7956 | III | 10 | 2 | 7956 |
| MdomPPK7 | XP_005190920.2 | NW_004769580.1 | 9432 | 16128 | 6696 | | III | | | |
| MdomPPK11 | XP_019895075.1 | NW_004770982.1 | 458690 | 464996 | 6306 | 329 | IV | 11 | 3 | 329 |
| MdomPPK16 | XP_019895076.1 | NW_004770982.1 | 465325 | 471554 | 6229 | 32504 | IV | | | 32504 |
| MdomPPK18 | XP_019895070.1 | NW_004770982.1 | 504058 | 506847 | 2789 | | IV | | | |

| Annotation | Original code | Scaffold | Start | Stop | Gene length (bp) | PPK distance (bp) | SF | Cluster N° | Genes per Cluster | Cluster Length (bp) | Exp. |
|---|---|---|---|---|---|---|---|---|---|---|---|
| Mper000000820.1 | MYZPE13164_G006_v1.0_000000820.1 | NW_019100468.1 | 1710025 | 1719014 | 8989 | | V | | | | Hemipteran Exp. |
| Mper000036460.1 | MYZPE13164_G006_v1.0_000036460.1 | NW_019100888.1 | 9094 | 15247 | 6153 | | V | | | | Hemipteran Exp. |
| MperPPK28 | MYZPE13164_G006_v1.0_000060760.1 | NW_019100902.1 | 446633 | 447822 | 1189 | | V | | | | |
| MperPPK23b | MYZPE13164_G006_v1.0_000035270.1 | NW_019101352.1 | 1195397 | 1209624 | **14227** | | VI | | | | |
| MperPPK23a | MYZPE13164_G006_v1.0_000066440.1 | NW_019101471.1 | 479949 | 482433 | 2484 | | VI | | | | |
| MperPPK9b | MYZPE13164_G006_v1.0_000070380.1 | NW_019101569.1 | 107437 | 116391 | 8954 | 12034 | II | 1 | 3 (MIXED) | 34448 | |
| Mper000070390.1 | MYZPE13164_G006_v1.0_000070390.1 | NW_019101569.1 | 128425 | 131260 | 2835 | 940 | I | | | | |
| MperPPK9a | MYZPE13164_G006_v1.0_000070370.1 | NW_019101569.1 | 132200 | 141885 | 9685 | | II | | | | |
| Mper000084580.1 | MYZPE13164_G006_v1.0_000084580.1 | NW_019102003.1 | 36312 | 56291 | 19979 | | I | | | | |
| Mper000138710.1 | MYZPE13164_G006_v1.0_000138710.1 | NW_019102087.1 | 1187 | 10394 | 9207 | | III | | | | |
| Mper000191120.1 | MYZPE13164_G006_v1.0_000191120.1 | NW_019102373.1 | 361416 | 371159 | 9743 | | V | | | | Hemipteran Exp. |
| Mper000138700.1 | MYZPE13164_G006_v1.0_000138700.1 | NW_019103909.1 | 77820 | 87316 | 9496 | 18688 | III | 2 | 2 | 45054 | |
| Mper000154690.1 | MYZPE13164_G006_v1.0_000154690.1 | NW_019103909.1 | 106004 | 122874 | 16870 | | III | | | | |
| Mper000086800.1 | MYZPE13164_G006_v1.0_000086800.1 | NW_019104028.1 | 122336 | 125706 | 3370 | 250 | III | 3 | 2 (MIXED) | 13510 | |
| Mper000095070.1 | MYZPE13164_G006_v1.0_000095070.1 | NW_019104028.1 | 125956 | 135846 | 9890 | | V | | | | Hemipteran Exp. |
| Mper000154700.1 | MYZPE13164_G006_v1.0_000154700.1 | NW_019104098.1 | 41919 | 47181 | 5262 | | III | | | | |

| | | | | | | | | | | | |
|---|---|---|---|---|---|---|---|---|---|---|---|
| Mper000163900 | MYZPE13164_G006_v1.0_000163900.1 | NW_019104105.1 | 24837 | 37244 | 12407 | | I | | | | |
| Mper000164680.1 | MYZPE13164_G006_v1.0_000164680.1 | NW_019104115.1 | 93441 | 102517 | **9076** | | III | | | | |
| Mper000165800.1 | MYZPE13164_G006_v1.0_000165800.1 | NW_019104344.1 | 673716 | 679350 | 5634 | | V | | | | Hemipteran Exp. |
| **Annotation** | **Original code** | **Scaffold** | **Start** | **Stop** | **Gene length (bp)** | **PPK distance (bp)** | **SF** | **Cluster N°** | **Genes per Cluster** | **Cluster Length (bp)** | **Exp.** |
| Pxyl21610.t1 | g21610.t1 | unitig_1196 | 40893 | 46826 | 5933 | | I | | | | |
| Pxyl11832.t1 | g11832.t1 | unitig_12640 | 227494 | 229675 | 2181 | | I | | | | |
| Pxyl20100.t1 | g20100.t1 | unitig_13832 | 59184 | 63491 | 4307 | | I | | | | |
| Pxyl30278.t1 | g30278.t1 | unitig_14556 | 17514 | 24005 | 6491 | | II | | | | |
| PxylPPK23a | g18724.t1 | unitig_14565 | 500578 | 509284 | 8706 | 265 | VI | 1 | 2(MIXED ) | 11491 | |
| Pxyl18725.t1 | g18725.t1 | unitig_14565 | 509549 | 512069 | 2520 | | IV | | | | |
| Pxyl5537.t1 | g5537.t1 | unitig_15281 | 3455 | 8435 | 4980 | 5679 | V | 2 | 2 | 13917 | |
| Pxyl5538.t1 | g5538.t1 | unitig_15281 | 14114 | 17372 | 3258 | | V | | | | |
| PxylPPK23b | g25324.t1 | unitig_15466 | 10517 | 26773 | 16256 | | VI | | | | |
| Pxyl16919.t1 | g16919.t1 | unitig_15784 | 1145715 | 1151390 | 5675 | | IV | | | | |
| Pxyl10292.t1 | g10292.t1 | unitig_15805 | 659350 | 669691 | 10341 | 2245 | V | 3 | 2 | 17570 | |
| Pxyl10294.t1 | g10294.t1 | unitig_15805 | 671936 | 676920 | 4984 | | V | | | | |
| PxylPPK9a | g18897.t1 | unitig_15807 | 145581 | 158882 | 13301 | | II | | | | |
| PxylPPK28a | g35214.t1 | unitig_15817 | 34236 | 44945 | 10709 | 9303 | V | 4 | 2 | 23484 | |
| PxylPPK28b | g35217.t1 | unitig_15817 | 54248 | 57720 | 3472 | | V | | | | |
| Pxyl34278.t1 | g34278.t1 | unitig_15944 | 101480 | 131229 | 29749 | 8142 | IV | 5 | 2 | 45747 | |
| Pxyl34279.t1 | g34279.t1 | unitig_15944 | 139371 | 147227 | 7856 | | IV | | | | |
| PxylPPK17 | g893.t1 | unitig_16006 | 718317 | 735191 | 16874 | | VII | | | | |
| Pxyl26144.t2 | g26144.t2 | unitig_16026 | 375115 | 378983 | 3868 | | IV | | | | |
| PxylPPK6b | g26485.t1 | unitig_16080 | 331017 | 333312 | 2295 | 378 | IV | 6 | 2 | 7105 | |
| PxylPPK6a | g26486.t1 | unitig_16080 | 333690 | 338122 | 4432 | | IV | | | | |
| PxylPPK13a | g16424.t1 | unitig_16100 | 2217428 | 2222992 | 5564 | 24036 | I | 7 | 2 | 33595 | |
| PxylPPK13b | g16431.t1 | unitig_16100 | 2247028 | 2251023 | 3995 | | I | | | | |
| Pxyl10539.t1 | g10539.t1 | unitig_16110 | 244898 | 246496 | 1598 | | IV | | | | |
| Pxyl36750.t1 | g36750.t1 | unitig_18 | 98987 | 99808 | 821 | | I | | | | |
| Pxyl26358.t1 | g26358.t1 | unitig_2459 | 878052 | 882390 | 4338 | | V | | | | |
| PxylPPK25 | g28101.t1 | unitig_2685 | 26966 | 48299 | 21333 | | II | | | | |
| Pxyl36719.t1 | g36719.t1 | unitig_2827 | 144068 | 147714 | 3646 | 292 | V | 8 | 2 | 7812 | |
| Pxyl36720.t1 | g36720.t1 | unitig_2827 | 148006 | 151880 | 3874 | | V | | | | |
| Pxyl12604.t1 | g12604.t1 | unitig_4463 | 96337 | 103748 | 7411 | | V | | | | |
| Pxyl15324.t1 | g15324.t1 | unitig_5355 | 1 | 3091 | 3090 | | I | | | | |
| PxylPPK9b | g31367.t1 | unitig_5670 | 6982 | 8247 | 1265 | | II | | | | |
| Pxyl18724.t1 | g18344.t1 | unitig_686 | 833792 | 839585 | 5793 | | IV | | | | |
| Pxyl15752.t1 | g15752.t1 | unitig_7489 | 1 | 2984 | 2983 | | II | | | | |
| PxylPPK16a | g28744.t1 | unitig_88 | 123443 | 132115 | 8672 | 61 | IV | 9 | 2 | 23467 | |
| PxylPPK16b | g28745.t1 | unitig_88 | 132176 | 146910 | 14734 | | IV | | | | |
| **Annotation** | **Original code** | **Scaffold** | **Start** | **Stop** | **Gene length (bp)** | **PPK distance (bp)** | **SF** | **Cluster N°** | **Genes per Cluster** | **Cluster Length (bp)** | **Exp.** |

| | | | | | | | | | | | |
|---|---|---|---|---|---|---|---|---|---|---|---|
| Rpro000048 | RPRC000048 | KQ034099 | 773670 | 785529 | 11859 | | V | | | | |
| Rpro000341 | RPRC000341 | KQ034159 | 876862 | 883313 | 6451 | | I | | | | |
| Rpro013510 | RPRC013510 | KQ034125 | 1139713 | 1164755 | 25042 | | V | | | | Hemipteran Exp. |
| Rpro014276 | RPRC014276 | KQ034076 | 397336 | 409703 | 12367 | | III | | | | |
| RproPPK23 | RPRC000099 | KQ034059 | 4941909 | 4950584 | 8675 | | VI | | | | |
| RproPPK28 | RPRC000471 | KQ034277 | 672067 | 685665 | 13598 | | V | | | | |
| RproPPK9 | Abs. VectorBase | KQ034064 | 1073569 | 1128473 | 54904 | | I | 1 | 2 | 59890 | |
| RproPPKlike5 | Abs. VectorBase | KQ034064 | 1068583 | 1072274 | 3691 | 1295 | I | | | | |
| RproPPKlike6 | Abs. VectorBase | KQ034335 | 409224 | 414959 | 5735 | | I | | | | |
| RproPPKlike7 | Abs. VectorBase | KQ034607 | 98017 | 111734 | 13717 | | II | | | | |

| **Annotation** | **Original code** | **Scaffold** | **Start** | **Stop** | **Gene length (bp)** | **PPK distance (bp)** | **SF** | **Cluster N°** | **Genes per Cluster** | **Cluster Length (bp)** | **Exp.** |
|---|---|---|---|---|---|---|---|---|---|---|---|
| Sfru031715 | SFRICE026402 | SFRU_RICE_000720 | 278 | 5433 | 5155 | | II | | | | |
| SfruPPK28 | SFRICE001686 | SFRU_RICE_000936 | 26740 | 31033 | 4293 | | V | | | | |
| Sfru034560 | SFRICE030732 | SFRU_RICE_001926 | 6199 | 7548 | 1349 | | V | | | | |
| SfruPPK23 | SFRICE002679 | SFRU_RICE_004623 | 7730 | 11361 | 3631 | | VI | | | | |
| SfruPPK6a | SFRICE012692 | SFRU_RICE_004773 | 12126 | 23265 | 11139 | | IV | | | | |
| Sfru030732 | SFRICE025969 | SFRU_RICE_005059 | 3529 | 7356 | 3827 | | I | | | | |
| Sfru020430 | SFRICE019222 | SFRU_RICE_005421 | 8424 | 12431 | 4007 | | IV | | | | |
| SfruPPK6c | SFRICE034560 | SFRU_RICE_005813 | 260 | 2244 | 1984 | | IV | | | | |
| Sfru009385 | SFRICE009385 | SFRU_RICE_007248 | 10042 | 14301 | 4259 | | I | | | | |
| Sfru023370 | SFRICE020430 | SFRU_RICE_007927 | 19220 | 23277 | 4057 | | II | | | | |
| Sfru002083 | SFRICE002083 | SFRU_RICE_008023 | 71651 | 77520 | 5869 | 4389 | IV | 1 | 2 (MIXED) | 17662 | |
| SfruPPK13a | SFRICE002084 | SFRU_RICE_008023 | 81909 | 89313 | 7404 | | I | | | | |
| Sfru024850 | SFRICE023370 | SFRU_RICE_008308 | 4664 | 10125 | 5461 | | V | | | | |
| SfruPPK6b | SFRICE032362 | SFRU_RICE_011410 | 1177 | 16678 | 15501 | | IV | | | | |
| Sfru019222 | SFRICE018167 | SFRU_RICE_011675 | 8206 | 13128 | 4922 | | IV | | | | |
| Sfru032362 | SFRICE030048 | SFRU_RICE_012279 | 27 | 5462 | 5435 | | IV | | | | |
| SfruPPK25a | SFRICE017730 | SFRU_RICE_012413 | 1702 | 9757 | 8055 | | II | | | | |
| Sfru017730 | SFRICE016175 | SFRU_RICE_013775 | 8212 | 14404 | 6192 | | IV | | | | |
| Sfru035107 | SFRICE031715 | SFRU_RICE_015948 | 208 | 7364 | 7156 | | IV | | | | |
| SfruPPK13c | SFRICE008543 | SFRU_RICE_01650 | 6546 | 10271 | 3725 | 7827 | I | 2 | 2 | 13222 | |

| | | | | | | | |
|---|---|---|---|---|---|---|---|
| | | 9 | | | | | |
| SfruPPK13b | SFRICE008547 | SFRU_RICE_016509 | 18098 | 19768 | 1670 | | I |
| SfruPPK25b | SFRICE024850 | SFRU_RICE_019662 | 10788 | 16979 | 6191 | | II |
| Sfru023811 | SFRICE020475 | SFRU_RICE_021761 | 1336 | 5690 | 4354 | | V |
| SfruPPK17 | SFRICE023811 | SFRU_RICE_025934 | 2599 | 7857 | 5258 | | VII |
| SfruPPK9 | SFRICE035107 | SFRU_RICE_027176 | 190 | 2327 | 2137 | | II |

| **Annotation** | **Original code** | **Scaffold** | **Start** | **Stop** | **Gene length (bp)** | **PPK distance (bp)** | **SF** | **Cluster N°** | **Genes per Cluster** | **Cluster Length (bp)** | **Exp.** |
|---|---|---|---|---|---|---|---|---|---|---|---|
| TC012955 | TC012955 | LG10 | 79.375 | 84.993 | 5.618 | | V | | | | Coleopteran Exp. |
| Tcas001349 | TC001349 | ChLG2 | 19581203 | 19582637 | 1434 | | I | | | | Coleopteran Exp. |
| TcasPPK3 | TC004432 | ChLG2 | 6506795 | 6508631 | 1836 | | II | | | | |
| Tcas003563 | TC003563 | ChLG3 | 9753485 | 9757996 | 4511 | | V | | | | Coleopteran Exp. |
| Tcas002633 | TC002633 | ChLG3 | 23275372 | 23281135 | 5763 | 5417 | V | 1 | 4 | 49331 | Coleopteran Exp. |
| Tcas002632 | TC002632 | ChLG3 | 23286552 | 23297273 | 10721 | 3784 | V | | | | Coleopteran Exp. |
| Tcas002631 | TC002631 | ChLG3 | 23301057 | 23306206 | 5149 | 5197 | V | | | | Coleopteran Exp. |
| Tcas002630 | TC002630 | ChLG3 | 23311403 | 23324703 | 13300 | | V | | | | Coleopteran Exp. |
| Tcas002554 | TC002554 | ChLG3 | 26801513 | 26808830 | 7317 | | V | | | | Coleopteran Exp. |
| TcasPPK16 | TC013324 | ChLG5 | 14000637 | 14004226 | 3589 | | IV | | | | |
| TcasPPK27a | TC015541 | ChLG6 | 3416400 | 3417670 | 1270 | 5264 | IV | 2 | 2 | 5264 | |
| TcasPPK27b | TC015167 | ChLG6 | 3412406 | 3413659 | 1253 | | IV | | | | |
| Tcas009302 | TC009302 | ChLG7 | 2278516 | 2281226 | 2710 | | IV | | | | |
| TcasPPK13 | TC006095 | ChLG8 | 2339481 | 2349457 | 9976 | | I | | | | |
| Tcas006569 | TC006569 | ChLG8 | 14028979 | 14030981 | 2002 | 3070 | V | 3 | 9 | 118686 | |
| Tcas006570 | TC006570 | ChLG8 | 14034051 | 14036020 | 1969 | 4673 | V | | | | |
| Tcas006571 | TC006571 | ChLG8 | 14040693 | 14044923 | 4230 | 168 | V | | | | Coleopteran Exp. |
| Tcas006572 | TC006572 | ChLG8 | 14045091 | 14049320 | 4229 | 1281 | V | | | | Coleopteran Exp. |
| Tcas006573 | TC006573 | ChLG8 | 14050601 | 14053427 | 2826 | 2963 | V | | | | Coleopteran Exp. |
| Tcas006574 | TC006574 | ChLG8 | 14056390 | 14063498 | 7108 | 1462 | V | | | | Coleopteran Exp. |
| Tcas006575 | TC006575 | ChLG8 | 14064960 | 14067385 | 2425 | 469 | V | | | | Coleopteran Exp. |
| Tcas005542 | TC005542 | ChLG8 | 14067854 | 14069796 | 1942 | 75001 | V | | | | Coleopteran Exp. |
| Tcas006581 | TC006581 | ChLG8 | 14144797 | 14147665 | 2868 | | V | | | | Coleopteran Exp. |
| TcasPPK17 | TC005481 | ChLG8 | 15168356 | 15169929 | 1573 | | VII | | | | Coleopteran Exp. |

| | | | | | | | |
|---|---|---|---|---|---|---|---|
| TcasPPK23 | TC012362 | ChLG9 | 8472347 | 8475659 | 3312 | | VI |
| TcasPPK28 | TC002095 | NW_015452015.1 | 14.023 | 17.902 | 3.879 | | V |
| TcasPPK9 | TC002260 | NW_015452015.1 | 731636 | 737466 | 5830 | | II |

**Table S5. Results of likelihood ratio test and parameter estimates under the best-fitting model for each orthogroup.** ω-values were obtained with the M0 model. LR = Likelihood ratio (2ΔL). Estimated parameters were obtained under M8 model (when significant, marked with *): P0 = proportion of sites that follow a beta distribution with 10 omega classes (0 ≤ ω ≥ 1); P1 = proportion of sites in the extra class with ω ≥ 1; location of predicted sites under positive selection and their posterior probability in parentheses.

| Orthogroup | ω | M8 vs M7 LR | M8 vs M8a LR | Estimated parameters | Location of predicted sites under positive selection (posterior probability) |
|---|---|---|---|---|---|
| *ppk2* | 0.15 | -0.003 | - | | |
| *ppk3* | 0.05 | 5.48 | - | | |
| *ppk6* | 0.1 | 30.29 * | 31.56* | $P_0$ = 0.97; $P_1$ = 0.03; $\omega_1$ = 4.8 | 1076 (0.96); 1160 (0.99) |
| *ppk9* | 0.06 | 3.21 | - | | |
| *ppk10* | 0.07 | -0.001 | - | | |
| *ppk13* | 0.02 | 5.9 | - | | |
| *ppk15* | 0.04 | -0.001 | - | | |
| *ppk16* | 0.16 | -0.005 | - | | |
| *ppk17* | 0.01 | 6.12 | - | | |
| *ppk19* | 0.15 | 3.11 | - | | |
| *ppk23* | 0.15 | 1 | - | | |
| *ppk25* | 0.1 | 56.33* | 56.21* | $P_0$ = 0.93; $P_1$ = 0.07; $\omega_1$ = 60.1 | 329 (0.98), 565 (0.98), 567 (0.98) |
| *ppk27* | 0.11 | 50.56* | 43.87* | $P_0$ = 0.98; $P_1$ = 0.02; $\omega_1$ = 314.1 | 10 (0.96), 17 (0.99), 19 (0.98) |
| *ppk28* | 0.08 | 3.7 | - | | |
| *ppk29* | 0.08 | -0.002 | - | | |
| *ppk31* | 0.03 | 0.8 | - | | |
| Coleopteran expansion | 0.08 | 2.43 | - | | |
| *Bl. germanica* expansion | 0.21 | 10.75* | 3.87 | | |
| Culicid expansion | 0.14 | 6.33 | - | | |
| Hemipteran expansion | 0.2 | 8.11 | - | | |

### Table S6. Functional information reported for PPKs

| Subfamily | Gene | Function and expression reported |
|---|---|---|
| I | *ppk13* | Expressed in hairs from the legs, tarsi and wings of the *D. melanogaster.* Also expressed in one terminal and four dorsal organ neurons in *D. melanogaster* larva (L. Liu, Leonard, et al., 2003) |
| | *ppk15* | Unknown function and expression data is absent |
| | *ppk29* | - Postsynaptic regulation of excitatory neurotransmission in larvae of *D. melanogaster* (A. Hill et al., 2017)<br>- Regulates neuronal excitability in *D. melanogaster* (Zheng et al., 2014)<br>- Expressed with *ppk23* and *ppk25* in the antennae, proboscis and legs. They are involved in sensing *Drosophila* cuticular hydrocarbons pheromones and mediate sexual behavior (T. Liu et al., 2020; Thistle et al., 2012; Vijayan et al., 2014)<br>- Expressed in whole fly, larval and adult carcass, larval central nervous system and in male accessory, thoracicoabdominal, salivary glands and midgut of *D. melanogaster* adults (Zelle et al., 2013)<br>- Mediates high salt taste with *ppk21* and *ppk27* in *D. melanogaster* (Lee et al., 2017) |
| | *ppk31* | Unknown function. It is expressed in larval midgut, adult carcass, male accessory, and fat body, salivary glands and crop of *D. melanogaster* adults (Zelle et al., 2013) |
| II | *ppk3* | Unknown function. Expressed in larval and adult tubule (Zelle et al., 2013) and in the reproductive female system (Wasbrough et al., 2010) |
| | *ppk9* | Unknown function and it is expressed in midgut, salivary glands, fat body and hindgut of larval, in adult carcass, male accessory, fat body, hindgut, midgut, crop, salivary gland and thoracicoabdominal of *D. melanogaster* adults (Zelle et al., 2013) |
| | *ppk10* | - Expressed in *D. melanogaster* wing and potentially involved in pheromone reception (He et al., 2019; G. Wang et al., 2010)<br>- Expressed in larval fat body, testis and salivary glands of adults (Zelle et al., 2013)<br>- Wing morphogenesis in *D. melanogaster* (George et al., 2019) |
| | *ppk25* | - High expressed in the *D. melanogaster* male appendages (legs, wings, and antennae) and it is responsible for gustatory and olfactory detection of female pheromones (Lin et al., 2005; Menuz et al., 2014; Starostina et al., 2012)<br>- Expressed with *ppk23* and *ppk29* in the antennae, proboscis and legs. They are involved in sensing *Drosophila* cuticular hydrocarbons pheromones and mediate sexual behavior (T. Liu et al., 2020; Thistle et al., 2012; Toda et al., 2012; Vijayan et al., 2014)<br>- Amplification of olfactory response by CBMs and *fruitless* (Ng et al., 2019; Zhang, Ng, Neville, Goodwin, Su, et al., 2020)<br>- Wing morphogenesis in *D. melanogaster* (George et al., 2019) |
| III | *ppk7* | Expressed in tracheal system of *D. melanogaster* larvae (L. Liu, Johnson, et al., 2003) |
| | *ppk14* | Expressed in tracheal system of *D. melanogaster* larvae (L. Liu, Johnson, et al., 2003) |
| | *ppk19* | - Sensing of low Na+ and K+ salts in *D. melanogaster* with *ppk11* (L. Liu, Leonard, et al., 2003)<br>- Wing morphogenesis in *D. melanogaster* (George et al., 2019)<br>- Expressed in tracheal system (Liu et al. 2003b) and testis of *D. melanogaster* adults (Zelle et al., 2013) |
| | *ppk20* | Unknown function. Expressed in the wing disc of the larval stage (Organista et al., 2015) |
| | *ppk21* | - Mediate high salt taste with *ppk27* and *ppk29* in *D. melanogaster* (Lee et al., 2017) |
| | *ppk30* | - Proprioceptive movement and mechanical nociception in *D. melanogaster* larvae. Also it is activated by lowering extracellular pH (Jang et al., 2019)<br>- Wing morphogenesis in *D. melanogaster* (George et al., 2019)<br>- Expressed in salivary glands of *D. melanogaster* adults (Zelle et al., 2013) |
| IV | *ppk4* | Expressed in tracheal system and it is involved in liquid clearance (L. Liu, Johnson, et al., 2003) |
| | *ppk6* | Wing morphogenesis in *D. melanogaster* (George et al., 2019). Expressed in larval carcass and hindgut, salivary glands and hindgut in adult (Zelle et al., 2013) |
| | *ppk11* | - Expressed in tracheal system and it is involved in liquid clearance (L. Liu, Johnson, et al., 2003)<br>- Presynaptic homeostatic plasticity with *ppk16* and *ppk1* (Orr et al., 2017; Younger et al., 2013)<br>- Sensing of low Na+ and K+ salts in *D. melanogaster* with *ppk19* (L. Liu, Leonard, et al., 2003) |
| | *ppk16* | Presynaptic homeostatic plasticity with *ppk1* and *ppk11* ((Orr et al., 2017; Younger et al., 2013). Expressed in larval and adult salivary glands, adult carcass, male accessory and hindgut of adults (Zelle et al., 2013) |
| | *ppk18* | Unknown function and expression data is absent |
| | *ppk22* | Unknown function. It is expressed in the whole fly, larval trachea, fat body and hindgut; adult carcass and male accessory of *D. melanogaster* (Zelle et al., 2013) |
| | *ppk24* | Unknown function and expression data is absent |
| | *ppk27* | - It is expressed in adult carcass, thoracicoabdominal and crop of *D. melanogaster* adults (Zelle et al., 2013).<br>- Mediates high salt taste with *ppk21* and *ppk29* in *D. melanogaster* (Lee et al., 2017) |
| V | *ppk1* (*ppk*) | - Mechanosensation in *D. melanogaster* larvae with *ppk2* ((Adams et al., 1998; Zelle et al., 2013)<br>- Mechanical nociception stimuli in *D. melanogaster* larval ((Zhong et al., 2010)<br>- Mediate larval aversion to dry surface environments with Painless (Johnson & Carder, 2012) |

| | | |
|---|---|---|
| | | - Presynaptic homeostatic plasticity with *ppk16* and *ppk11* (Orr et al., 2017; Younger et al., 2013)<br>- Wing morphogenesis in *D. melanogaster* (George et al., 2019)<br>- Proper locomotion in *D. melanogaster* larvae with *ppk26* (Gorczyca et al., 2014)<br>- Acid sensing (pH between 9 and 15) in *D. melanogaster* larvae (Boiko et al., 2012)<br>- Expressed in larval carcass of *D. melanogaster* (Zelle et al., 2013) |
| | *ppk2 (rpk)* | - Sense of touch in *Drosophila* larvae (Tsubouchi et al., 2012)<br>- Mechanosensation in *D. melanogaster* larvae with *ppk1* (Adams et al., 1998)<br>- Wing morphogenesis in *D. melanogaster* (George et al., 2019)<br>- Expressed in whole fly, larval fat body and central nervous system, adult carcass, virgin spermatheca, fat body and ovary, and salivary glands of *D. melanogaster* adults (Zelle et al., 2013) |
| | *ppk5* | Unknown function. Expressed in virgin spermatheca of *D. melanogaster* (Zelle et al., 2013) |
| | *ppk8* | Unknown function. Expressed in adult fat body (Xu et al., 2011) and in salivary glands of *D. melanogaster* (Zelle et al., 2013) |
| | *ppk12* | Mechanosensation in *D. melanogaster* larvae with *ppk2* (Adams et al., 1998; Zelle et al., 2013). Expressed in larval hindgut, male accessory and adult salivary glands (Zelle et al., 2013). |
| | *ppk26* | - Mechanosensation in *D. melanogaster* larvae with *ppk1* ((Zelle et al., 2013)<br>- Mechanical nocioception in *D. melanogaster* with *ppk1* ((Y. Guo et al., 2014)<br>- Proper locomotion in *D. melanogaster* larvae with *ppk1* (Gorczyca et al., 2014)<br>- Expressed in larval and adult carcass, thoracicoabdominal and salivary glands of *D. melanogaster* adults (Zelle et al., 2013) |
| | *ppk28 (ppk301 )* | - Expressed in labellum, legs and wing margins and it is involved in gustatory water reception in *D. melanogaster* (Cameron et al., 2010; Chen et al., 2010)<br>- Expressed in the tracheal system of *D. melanogaster* larvae (L. Liu, Johnson, et al., 2003)Liu et al. 2003b)<br>- Expressed in larval central nervous system and thoracicoabdominal and salivary glands of *D. melanogaster* adults (Zelle et al., 2013)<br>- Expressed in proboscis and tarsi and it controls egg-laying initiation and choice in *Ae. aegypti* (Matthews et al., 2019) |
| VI | *ppk23* | - Expressed in gustatory neurons of proboscis in both adult sexes and considered a contact receptor that mediates with *ppk29* the detection of cuticular hydrocarbons involved in sexual behavior of *D. melanogaster* (Thistle et al., 2012)<br>- Expressed in antennae with *ppk25* and *ppk29* and they are necessary for *Drosophila* females to become receptive to mating (Vijayan et al., 2014)<br>- Expressed in male legs and mediates: 1) male anti-aphrodisiac pheromones detection with *fruitless* in the "M" cells 2) female aphrodisiac pheromone detection with *ppk25, ppk29* and *fruitless* in the "F" cells ((Thistle et al., 2012; Toda et al., 2012; Vijayan et al., 2014)<br>- Expressed in the fly labellum and it detects low salt concentrations (Jaeger et al., 2018).<br>- Expressed in whole, adult carcass, mated spermatheca, virgin spermatheca, salivary glands, crop and midgut of *D. melanogaster* adults (Zelle et al., 2013).<br>- Amplification of olfactory response mediated by *fruitless* (Zhang, Ng, Neville, Goodwin, & Su, 2020) |
| VII | *ppk17* | Wing morphogenesis in *D. melanogaster* (George et al., 2019). It is expressed in the larval carcass, tubule, hindgut and salivary glands; and hindgut, midgut, crop and salivary glands of *D. melanogaster* adults (Zelle et al., 2013). |

**Table S7. Number of sensory receptor families from different insect species.** Data of OR, GR and IR genes from *Myzus persicae* are not available. ND: No data,

| Insect | ORs | GRs | IRs | PPKs | Total | Reference |
|---|---|---|---|---|---|---|
| *Aedes aegypti* | 117 | 72 | 135 | 32 | 356 | Matthews et al. (2018) |
| *Aedes albopictus* | 82 | 30 | 60 | 49 | 221 | Lombardo et al. (2017) |
| *Anopheles gambiae* | 79 | 76 | 46 | 27 | 228 | C. A. Hill et al, (2002); Jason Pitts et al. (2017) |
| *Culex quinquefasciatus* | 180 | 123 | 59 | 48 | 410 | Arensburger et al. (2010); Leal et al. (2013) |
| *Drosophila melanogaster* | 62 | 68 | 66 | 31 | 227 | Robertson et al., 2003) |
| *Glossina morsitans* | 46 | 14 | 19 | 15 | 94 | Watanabe et al., 2014) |
| *Musca domestica* | 86 | 103 | 110 | 59 | 358 | Scott et al., 2014) |
| *Bombyx mori* | 70 | 76 | 25 | 18 | 189 | Gouin et al., 2017; H. Guo et al. (2017); van Schooten et al. (2016); Wanner & Robertson (2008) |
| *Danaus plexippus* | 64 | 47 | 27 | 18 | 156 | Zhan et al. (2011) |
| *Plutella Xylotella* | 95 | 69 | 37 | 36 | 237 | Engsontia et al. (2014) |
| *Spodoptera frugiperda* | 69 | 231 | 43 | 25 | 368 | Gouin et al. (2017) |
| *Anoplophora glabripennis* | 132 | 234 | 72 | 17 | 455 | McKenna et al. (2016) |
| *Dendroctonus ponderosae* | 86 | 57 | 60 | 14 | 217 | Andersson et al. (2019) |
| *Leptinotarsa decemlineata* | 75 | 144 | 27 | 10 | 256 | Schoville et al. (2018) |
| *Tribolium castaneum* | 264 | 219 | 72 | 27 | 582 | Abdel-latief, (2007); Engsontia et al. (2008) |
| *Atta cephalotes* | 376 | 89 | 18 | 6 | 489 | Engsontia et al. (2015); Koch et al. (2013) |
| *Apis mellifera* | 177 | 14 | 10 | 8 | 209 | Brand & Ramírez (2017) |
| *Bombus impatiens* | 159 | 24 | ND | 9 | 192 | Sadd et al. (2015) |
| *Camponotus floridanus* | 352 | 11 | 31 | 8 | 402 | Bonasio et al. (2010); Zhou et al. (2012) |
| *Pediculus humanus* | 10 | 8 | 12 | 1 | 31 | Kirkness et al. (2010) |
| *Acyrthosiphon pisum* | 79 | 77 | 11 | 21 | 188 | Smadja et al. (2009) |
| *Cimex lectularius* | 49 | 36 | 30 | 10 | 125 | Benoit et al. (2016) |
| *Myzus persicae* | ND | ND | ND | 19 | 19 | - |
| *Rhodnius prolixus* | 111 | 30 | 27 | 10 | 178 | Mesquita et al. (2015) |
| *Locusta migratoria* | 142 | 75 | 32 | 19 | 268 | X. Wang et al. (2014); Z. Wang et al. (2015) |
| *Blattella germanica* | 134 | 545 | 897 | 41 | 1617 | Robertson et al. (2018) |

Abdel-latief, M. (2007). A Family of Chemoreceptors in Tribolium castaneum (Tenebrionidae: Coleoptera). *PLoS ONE*, *2*(12), e1319. https://doi.org/10.1371/journal.pone.0001319

Adams, C. M., Anderson, M. G., Motto, D. G., Price, M. P., Johnson, W. a, & Welsh, M. J. (1998). Ripped Pocket and Pickpocket, Novel. *Cell*, *140*(1), 143–152.

Andersson, M. N., Keeling, C. I., & Mitchell, R. F. (2019). Genomic content of chemosensory genes correlates with host range in wood-boring beetles (Dendroctonus ponderosae, Agrilus planipennis, and Anoplophora glabripennis). *BMC Genomics*, *20*(1), 690. https://doi.org/10.1186/s12864-019-6054-x

Arensburger, P., Megy, K., Waterhouse, R. M., Abrudan, J., Amedeo, P., Antelo, B., Bartholomay, L., Bidwell, S., Caler, E., Camara, F., Campbell, C. L., Campbell, K. S., Casola, C., Castro, M. T., Chandramouliswaran, I., Chapman, S. B., Christley, S., Costas, J., Eisenstadt, E., … Atkinson, P. W. (2010). Sequencing of Culex quinquefasciatus establishes a platform for mosquito comparative genomics. *Science*, *330*(6000), 86–88. https://doi.org/10.1126/science.1191864

Benoit, J. B., Adelman, Z. N., Reinhardt, K., Dolan, A., Poelchau, M., Jennings, E. C., Szuter, E. M., Hagan, R. W., Gujar, H., Shukla, J. N., Zhu, F., Mohan, M., Nelson, D. R., Rosendale, A. J., Derst, C., Resnik, V., Wernig, S., Menegazzi, P., Wegener, C., … Richards, S. (2016). Unique features of a global human ectoparasite identified through sequencing of the bed bug genome. *Nature Communications*, *7*(1), 22. https://doi.org/10.1038/ncomms10165

Boiko, N., Kucher, V., Stockand, J. D., & Eaton, B. A. (2012). Pickpocket1 is an ionotropic molecular sensory transducer. *Journal of Biological Chemistry*, *287*(47), 39878–39886. https://doi.org/10.1074/jbc.M112.411736

Bonasio, R., Zhang, G., Ye, C., Mutti, N. S., Fang, X., Qin, N., Donahue, G., Yang, P., Li, Q., Li, C., Zhang, P., Huang, Z., Berger, S. L., Reinberg, D., Wang, J., & Liebig, J. (2010). Genomic comparison of the ants Camponotus floridanus and Harpegnathos saltator. *Science*, *329*(5995), 1068–1071. https://doi.org/10.1126/science.1192428

Brand, P., & Ramírez, S. R. (2017). The evolutionary dynamics of the odorant receptor gene family in corbiculate bees. *Genome Biology and Evolution*, *9*(8), 2023–2036. https://doi.org/10.1093/gbe/evx149

Cameron, P., Hiroi, M., Ngai, J., & Scott, K. (2010). The molecular basis for water taste in Drosophila. *Nature*, *465*(7294), 91–95. https://doi.org/10.1038/nature09011

Chen, Z., Wang, Q., & Wang, Z. (2010). The amiloride-sensitive epithelial Na+ channel PPK28 is essential for Drosophila gustatory water reception. *Journal of Neuroscience*, *30*(18), 6247–6252. https://doi.org/10.1523/JNEUROSCI.0627-10.2010

Engsontia, P., Sanderson, A. P., Cobb, M., Walden, K. K. O., Robertson, H. M., & Brown, S. (2008). The red flour beetle's large nose: An expanded odorant receptor gene family in Tribolium castaneum. *Insect Biochemistry and Molecular Biology*, *38*(4), 387–397. https://doi.org/10.1016/j.ibmb.2007.10.005

Engsontia, P., Sangket, U., Chotigeat, W., & Satasook, C. (2014). Molecular evolution of the odorant and gustatory receptor genes in lepidopteran insects: Implications for their adaptation and speciation. *Journal of Molecular Evolution*, *79*(1–2), 21–39. https://doi.org/10.1007/s00239-014-9633-0

Engsontia, P., Sangket, U., Robertson, H. M., & Satasook, C. (2015). Diversification of the ant odorant receptor gene family and positive selection on candidate cuticular hydrocarbon receptors. *BMC Research Notes*, *8*(1), 1–13. https://doi.org/10.1186/s13104-015-1371-x

George, L. F., Pradhan, S. J., Mitchell, D., Josey, M., Casey, J., Belus, M. T., Fedder, K. N., Raj Dahal, G., & Bates, E. A. (2019). Ion channel contributions to wing development in Drosophila melanogaster. *G3: Genes, Genomes, Genetics*, *9*(4), 999–1008. https://doi.org/10.1534/g3.119.400028

Gorczyca, D. A., Younger, S., Meltzer, S., Kim, S. E., Cheng, L., Song, W., Lee, H. Y., Jan, L. Y., & Jan, Y. N. (2014). Identification of Ppk26, a DEG/ENaC Channel Functioning with Ppk1 in a Mutually Dependent Manner to Guide Locomotion Behavior in Drosophila. *Cell Reports*, *9*(4), 1446–1458. https://doi.org/10.1016/j.celrep.2014.10.034

Gouin, A., Bretaudeau, A., Nam, K., Gimenez, S., Aury, J. M., Duvic, B., Hilliou, F., Durand, N., Montagné, N., Darboux, I., Kuwar, S., Chertemps, T., Siaussat, D., Bretschneider, A., Moné, Y., Ahn, S. J., Hänniger, S., Grenet, A. S. G., Neunemann, D., … Fournier, P. (2017). Two genomes of highly polyphagous lepidopteran pests (Spodoptera frugiperda, Noctuidae) with different host-plant ranges. *Scientific Reports*, *7*(1), 1–12. https://doi.org/10.1038/s41598-017-10461-4

Guo, H., Cheng, T., Chen, Z., Jiang, L., Guo, Y., Liu, J., Li, S., Taniai, K., Asaoka, K., Kadono-Okuda, K., Arunkumar, K. P., Wu, J., Kishino, H., Zhang, H., Seth, R. K., Gopinathan, K. P., Montagné, N., Jacquin-Joly, E., Goldsmith, M. R., … Mita, K. (2017). Expression map of a complete set of gustatory receptor genes in chemosensory organs of Bombyx mori. *Insect Biochemistry and Molecular Biology*, *82*, 74–82. https://doi.org/10.1016/j.ibmb.2017.02.001

Guo, Y., Wang, Y., Wang, Q., & Wang, Z. (2014). The Role of PPK26 in Drosophila Larval Mechanical Nociception. *Cell Reports*, *9*(4), 1183–1190. https://doi.org/10.1016/j.celrep.2014.10.020

He, Z., Luo, Y., Shang, X., Sun, J. S., & Carlson, J. R. (2019). Chemosensory sensilla of the Drosophila wing express a candidate ionotropic pheromone receptor. *PLoS Biology*, *17*(5), e2006619. https://doi.org/10.1371/journal.pbio.2006619

Hill, A., Zheng, X., Li, X., McKinney, R., Dickman, D., & Ben-Shahar, Y. (2017). The Drosophila postsynaptic DEG/ENaC channel ppk29 contributes to excitatory neurotransmission. *Journal of Neuroscience*, *37*(12), 3171–3180. https://doi.org/10.1523/JNEUROSCI.3850-16.2017

Hill, C. A., Fox, A. N., Pitts, R. J., Kent, L. B., Tan, P. L., Chrystal, M. A., Cravchik, A., Collins, F. H., Robertson, H. M., & Zwiebel, L. J. (2002). G protein-coupled receptors in Anopheles gambiae. *Science*, *298*(5591), 176–178. https://doi.org/10.1126/science.1076196

Jaeger, A. H., Stanley, M., Weiss, Z. F., Musso, P. Y., Chan, R. C. W., Zhang, H., Feldman-Kiss, D., & Gordon, M. D. (2018). A complex peripheral code for salt taste in drosophila. *ELife*, *7*, 1–30. https://doi.org/10.7554/eLife.37167

Jang, W., Lee, S., Choi, S. I., Chae, H. S., Han, J., Jo, H., Hwang, S. W., Park, C. S., & Kim, C. (2019). Impairment of proprioceptive movement and mechanical nociception in Drosophila melanogaster larvae lacking Ppk30, a Drosophila member of the Degenerin/Epithelial Sodium Channel family. *Genes, Brain and Behavior*, *18*(5), 1–8. https://doi.org/10.1111/gbb.12545

Jason Pitts, R., Derryberry, S. L., Zhang, Z., & Zwiebel, L. J. (2017). Variant Ionotropic Receptors in the Malaria Vector Mosquito Anopheles gambiae Tuned to Amines and Carboxylic Acids. *Scientific Reports*, *7*(September 2016), 1–11. https://doi.org/10.1038/srep40297

Johnson, W. A., & Carder, J. W. (2012). Drosophila nociceptors mediate larval aversion to dry surface environments utilizing both the painless TRP Channel and the DEG/

ENaC subunit, PPK1. *PLoS ONE*, *7*(3). https://doi.org/10.1371/journal.pone.0032878

Kirkness, E. F., Haas, B. J., Sun, W., Braig, H. R., Perotti, M. A., Clark, J. M., Lee, S. H., Robertson, H. M., Kennedy, R. C., Elhaik, E., Gerlach, D., Kriventseva, E. V., Elsik, C. G., Graur, D., Hill, C. A., Veenstra, J. A., Walenz, B., Tubío, J. M. C., Ribeiro, J. M. C., … Pittendrigh, B. R. (2010). Genome sequences of the human body louse and its primary endosymbiont provide insights into the permanent parasitic lifestyle. *Proceedings of the National Academy of Sciences of the United States of America*, *107*(27), 12168–12173. https://doi.org/10.1073/pnas.1003379107

Koch, S. I., Groh, K., Vogel, H., Hannson, B. S., Kleineidam, C. J., & Grosse-Wilde, E. (2013). Caste-specific expression patterns of immune response and chemosensory related genes in the leaf-cutting ant, Atta vollenweideri. *PLoS ONE*, *8*(11), e81518. https://doi.org/10.1371/journal.pone.0081518

Leal, W. S., Choo, Y. M., Xu, P., Da Silva, C. S. B., & Ueira-Vieira, C. (2013). Differential expression of olfactory genes in the southern house mosquito and insights into unique odorant receptor gene isoforms. *Proceedings of the National Academy of Sciences of the United States of America*, *110*(46), 18704–18709. https://doi.org/10.1073/pnas.1316059110

Lee, M. J., Sung, H. Y., Jo, H., Kim, H. W., Choi, M. S., Kwon, J. Y., & Kang, K. J. (2017). Ionotropic receptor 76b is required for gustatory aversion to excessive na+ in drosophila. *Molecules and Cells*, *40*(10), 787–795. https://doi.org/10.14348/molcells.2017.0160

Lin, H., Mann, K. J., Starostina, E., Kinser, R. D., & Pikielny, C. W. (2005). A Drosophila DEG/ENaC channel subunit is required for male response to female pheromones. *Proceedings of the National Academy of Sciences of the United States of America*, *102*(36), 12831–12836. https://doi.org/10.1073/pnas.0506420102

Liu, L., Johnson, W. A., & Welsh, M. J. (2003). Drosophila DEG/ENaC pickpocket genes are expressed in the tracheal system, where they may be involved in liquid clearance. *Proceedings of the National Academy of Sciences of the United States of America*, *100*(4), 2128–2133. https://doi.org/10.1073/pnas.252785099

Liu, L., Leonard, A. S., Motto, D. G., Feller, M. A., Price, M. P., Johnson, W. A., & Welsh, M. J. (2003). Contribution of Drosophila DEG/ENaC genes to salt taste. *Neuron*, *39*(1), 133–146. https://doi.org/10.1016/S0896-6273(03)00394-5

Liu, T., Wang, Y., Tian, Y., Zhang, J., Zhao, J., & Guo, A. (2020). The receptor channel formed by ppk25, ppk29 and ppk23 can sense the *Drosophila* female pheromone 7,11-heptacosadiene. *Genes, Brain and Behavior*, *19*(2), e12529. https://doi.org/10.1111/gbb.12529

Lombardo, F., Salvemini, M., Fiorillo, C., Nolan, T., Zwiebel, L. J., Ribeiro, J. M., & Arcà, B. (2017). Deciphering the olfactory repertoire of the tiger mosquito Aedes albopictus. *BMC Genomics*, *18*(1). https://doi.org/10.1186/s12864-017-4144-1

Matthews, B. J., Dudchenko, O., Kingan, S. B., Koren, S., Antoshechkin, I., Crawford, J. E., Glassford, W. J., Herre, M., Redmond, S. N., Rose, N. H., Weedall, G. D., Wu, Y., Batra, S. S., Brito-Sierra, C. A., Buckingham, S. D., Campbell, C. L., Chan, S., Cox, E., Evans, B. R., … Vosshall, L. B. (2018). Improved reference genome of Aedes aegypti informs arbovirus vector control. *Nature*, *563*(7732), 501–507. https://doi.org/10.1038/s41586-018-0692-z

Matthews, B. J., Younger, M. A., & Vosshall, L. B. (2019). The ion channel ppk301 controls freshwater egg-laying in the mosquito aedes aegypti. *ELife*, *8*, 1–27. https://doi.org/10.7554/eLife.43963

McKenna, D. D., Scully, E. D., Pauchet, Y., Hoover, K., Kirsch, R., Geib, S. M., Mitchell, R. F., Waterhouse, R. M., Ahn, S. J., Arsala, D., Benoit, J. B., Blackmon, H., Bledsoe, T., Bowsher, J. H., Busch, A., Calla, B., Chao, H., Childers, A. K., Childers, C., … Richards, S. (2016). Genome of the Asian longhorned beetle (Anoplophora glabripennis), a globally significant invasive species, reveals key functional and evolutionary innovations at the beetle-plant interface. *Genome Biology*, *17*(1), 227. https://doi.org/10.1186/s13059-016-1088-8

Menuz, K., Larter, N. K., Park, J., & Carlson, J. R. (2014). An RNA-Seq Screen of the Drosophila Antenna Identifies a Transporter Necessary for Ammonia Detection. *PLoS Genetics*, *10*(11). https://doi.org/10.1371/journal.pgen.1004810

Mesquita, R. D., Vionette-Amaral, R. J., Lowenberger, C., Rivera-Pomar, R., Monteiro, F. A., Minx, P., Spieth, J., Carvalho, A. B., Panzera, F., Lawson, D., Torres, A. Q., Ribeiro, J. M. C., Sorgine, M. H. F., Waterhouse, R. M., Montague, M. J., Abad-Franch, F., Alves-Bezerra, M., Amaral, L. R., Araujo, H. M., … Oliveira, P. L. (2015). Genome of Rhodnius prolixus, an insect vector of Chagas disease, reveals unique adaptations to hematophagy and parasite infection. *Proceedings of the National Academy of Sciences of the United States of America*. https://doi.org/10.1073/pnas.1506226112

Ng, R., Salem, S. S., Wu, S. T., Wu, M., Lin, H. H., Shepherd, A. K., Joiner, W. J., Wang, J. W., & Su, C. Y. (2019). Amplification of Drosophila Olfactory Responses by a DEG/ENaC Channel. *Neuron*, *104*(5), 947-959.e5. https://doi.org/10.1016/j.neuron.2019.08.041

Organista, M. F., Martín, M., de Celis, J. M., Barrio, R., López-Varea, A., Esteban, N., Casado, M., & de Celis, J. F. (2015). The Spalt Transcription Factors Generate the Transcriptional Landscape of the Drosophila melanogaster Wing Pouch Central Region. *PLOS Genetics*, *11*(8), e1005370. https://doi.org/10.1371/journal.pgen.1005370

Orr, B. O., Gorczyca, D., Younger, M. A., Jan, L. Y., Jan, Y. N., & Davis, G. W. (2017). Composition and Control of a Deg/ENaC Channel during Presynaptic Homeostatic Plasticity. *Cell Reports*, *20*(8), 1855–1866. https://doi.org/10.1016/j.celrep.2017.07.074

Robertson, H. M., Baits, R. L., Walden, K. K. O., Wada-Katsumata, A., & Schal, C. (2018). Enormous expansion of the chemosensory gene repertoire in the omnivorous German cockroach *Blattella germanica*. *Journal of Experimental Zoology Part B: Molecular and Developmental Evolution*, *330*(5), 265–278. https://doi.org/10.1002/jez.b.22797

Robertson, H. M., Warr, C. G., & Carlson, J. R. (2003). Molecular evolution of the insect chemoreceptor gene superfamily in Drosophila melanogaster. *Proceedings of the National Academy of Sciences of the United States of America*, *100*(24), 14537–14542. https://doi.org/10.1073/pnas.2335847100

Sadd, B. M., Barribeau, S. M., Bloch, G., de Graaf, D. C., Dearden, P., Elsik, C. G., Gadau, J., Grimmelikhuijzen, C. J. P., Hasselmann, M., Lozier, J. D., Robertson, H. M., Smagghe, G., Stolle, E., Van Vaerenbergh, M., Waterhouse, R. M., Bornberg-Bauer, E., Klasberg, S., Bennett, A. K., Câmara, F., … Worley, K. C. (2015). The genomes of two key bumblebee species with primitive eusocial organization. *Genome Biology*, *16*(1), 76. https://doi.org/10.1186/s13059-015-0623-3

Schoville, S. D., Chen, Y. H., Andersson, M. N., Benoit, J. B., Bhandari, A., Bowsher, J. H., Brevik, K., Cappelle, K., Chen, M. J. M., Childers, A. K., Childers, C., Christiaens, O., Clements, J., Didion, E. M., Elpidina, E. N., Engsontia, P., Friedrich, M., García-Robles, I., Gibbs, R. A., … Richards, S. (2018). A model species for agricultural pest genomics: The genome of the Colorado potato beetle, Leptinotarsa decemlineata

(Coleoptera: Chrysomelidae). *Scientific Reports*, *8*(1), 1–18. https://doi.org/10.1038/s41598-018-20154-1

Scott, J. G., Warren, W. C., Beukeboom, L. W., Bopp, D., Clark, A. G., Giers, S. D., Hediger, M., Jones, A. K., Kasai, S., Leichter, C. A., Li, M., Meisel, R. P., Minx, P., Murphy, T. D., Nelson, D. R., Reid, W. R., Rinkevich, F. D., Robertson, H. M., Sackton, T. B., … Liu, N. (2014). Genome of the house fly, Musca domestica L., a global vector of diseases with adaptations to a septic environment. *Genome Biology*, *15*(10), 466. https://doi.org/10.1186/s13059-014-0466-3

Smadja, C., Shi, P., Butlin, R. K., & Robertson, H. M. (2009). Large Gene Family Expansions and Adaptive Evolution for Odorant and Gustatory Receptors in the Pea Aphid, Acyrthosiphon pisum. *Molecular Biology and Evolution*, *26*(9), 2073–2086. https://doi.org/10.1093/molbev/msp116

Starostina, E., Liu, T., Vijayan, V., Zheng, Z., Siwicki, K. K., & Pikielny, C. W. (2012). A drosophila DEG/ENaC subunit functions specifically in gustatory neurons required for male courtship behavior. *Journal of Neuroscience*, *32*(13), 4665–4674. https://doi.org/10.1523/JNEUROSCI.6178-11.2012

Thistle, R., Cameron, P., Ghorayshi, A., Dennison, L., & Scott, K. (2012). Contact chemoreceptors mediate male-male repulsion and male-female attraction during drosophila courtship. *Cell*, *149*(5), 1140–1151. https://doi.org/10.1016/j.cell.2012.03.045

Toda, H., Zhao, X., & Dickson, B. J. (2012). The Drosophila Female Aphrodisiac Pheromone Activates ppk23+ Sensory Neurons to Elicit Male Courtship Behavior. *Cell Reports*, *1*(6), 599–607. https://doi.org/10.1016/j.celrep.2012.05.007

Tsubouchi, A., Caldwell, J. C., & Tracey, W. D. (2012). Dendritic filopodia, ripped pocket, NOMPC, and NMDARs contribute to the sense of touch in Drosophila larvae. *Current Biology*, *22*(22), 2124–2134. https://doi.org/10.1016/j.cub.2012.09.019

van Schooten, B., Jiggins, C. D., Briscoe, A. D., & Papa, R. (2016). Genome-wide analysis of ionotropic receptors provides insight into their evolution in Heliconius butterflies. *BMC Genomics*, *17*(1), 254. https://doi.org/10.1186/s12864-016-2572-y

Vijayan, V., Thistle, R., Liu, T., Starostina, E., & Pikielny, C. W. (2014). Drosophila Pheromone-Sensing Neurons Expressing the ppk25 Ion Channel Subunit Stimulate Male Courtship and Female Receptivity. *PLoS Genetics*, *10*(3), e1004238. https://doi.org/10.1371/journal.pgen.1004238

Wang, G., Carey, A. F., Carlson, J. R., & Zwiebel, L. J. (2010). Molecular basis of odor coding in the malaria vector mosquito Anopheles gambiae. *Proceedings of the National Academy of Sciences of the United States of America*, *107*(9), 4418–4423. https://doi.org/10.1073/pnas.0913392107

Wang, X., Fang, X., Yang, P., Jiang, X., Jiang, F., Zhao, D., Li, B., Cui, F., Wei, J., Ma, C., Wang, Y., He, J., Luo, Y., Wang, Z., Guo, X., Guo, W., Wang, X., Zhang, Y., Yang, M., … Kang, L. (2014). The locust genome provides insight into swarm formation and long-distance flight. *Nature Communications*, *5*(1), 2957. https://doi.org/10.1038/ncomms3957

Wang, Z., Yang, P., Chen, D., Jiang, F., Li, Y., Wang, X., & Kang, L. (2015). Identification and functional analysis of olfactory receptor family reveal unusual characteristics of the olfactory system in the migratory locust. *Cellular and Molecular Life Sciences*, *72*(22), 4429–4443. https://doi.org/10.1007/s00018-015-2009-9

Wanner, K. W., & Robertson, H. M. (2008). The gustatory receptor family in the silkworm moth Bombyx mori is characterized by a large expansion of a single lineage of putative bitter receptors. *Insect Molecular Biology*, *17*(6), 621–629. https://doi.org/10.1111/j.1365-2583.2008.00836.x

Wasbrough, E. R., Dorus, S., Hester, S., Howard-Murkin, J., Lilley, K., Wilkin, E., Polpitiya, A., Petritis, K., & Karr, T. L. (2010). The Drosophila melanogaster sperm proteome-II (DmSP-II). *Journal of Proteomics*, *73*(11), 2171–2185. https://doi.org/10.1016/j.jprot.2010.09.002

Watanabe, J., Hattori, M., Berriman, M., Lehane, M. J., Hall, N., Solano, P., Aksoy, S., Hide, W., Touré, Y., Attardo, G. M., Darby, A. C., Toyoda, A., Hertz-Fowler, C., Larkin, D. M., Cotton, J. A., Sanders, M. J., Swain, M. T., Quail, M. A., Inoue, N., … Kawahara, Y. (2014). Genome sequence of the tsetse fly (Glossina morsitans ): Vector of African trypanosomiasis. *Science*, *344*(6182), 380–386. https://doi.org/10.1126/science.1249656

Xu, K., DiAngelo, J. R., Hughes, M. E., Hogenesch, J. B., & Sehgal, A. (2011). The circadian clock interacts with metabolic physiology to influence reproductive fitness. *Cell Metabolism*, *13*(6), 639–654. https://doi.org/10.1016/j.cmet.2011.05.001

Younger, M. A., Müller, M., Tong, A., Pym, E. C., & Davis, G. W. (2013). A presynaptic ENaC channel drives homeostatic plasticity. *Neuron*, *79*(6), 1183–1196. https://doi.org/10.1016/j.neuron.2013.06.048

Zelle, K. M., Lu, B., Pyfrom, S. C., & Ben-Shahar, Y. (2013). The genetic architecture of degenerin/epithelial sodium channels in drosophila. *G3: Genes, Genomes, Genetics*, *3*(3), 441–450. https://doi.org/10.1534/g3.112.005272

Zhan, S., Merlin, C., Boore, J. L., & Reppert, S. M. (2011). The monarch butterfly genome yields insights into long-distance migration. *Cell*, *147*(5), 1171–1185. https://doi.org/10.1016/j.cell.2011.09.052

Zhang, Y., Ng, R., Neville, M. C., Goodwin, S. F., & Su, C. Y. (2020). Distinct Roles and Synergistic Function of FruM Isoforms in Drosophila Olfactory Receptor Neurons. *Cell Reports*, *33*(11), 108516. https://doi.org/10.1016/j.celrep.2020.108516

Zhang, Y., Ng, R., Neville, M. C., Goodwin, S. F., Su, C., Zhang, Y., Ng, R., Neville, M. C., Goodwin, S. F., & Su, C. (2020). Article Distinct Roles and Synergistic Function of Fru M Isoforms in Drosophila Olfactory Receptor Neurons ll ll Distinct Roles and Synergistic Function of Fru M Isoforms in Drosophila Olfactory Receptor Neurons. *CellReports*, *33*(11), 108516. https://doi.org/10.1016/j.celrep.2020.108516

Zheng, X., Valakh, V., DiAntonio, A., & Ben-Shahar, Y. (2014). Natural antisense transcripts regulate the neuronal stress response and excitability. *ELife*, *2014*(3). https://doi.org/10.7554/eLife.01849

Zhong, L., Hwang, R. Y., & Tracey, W. D. (2010). Pickpocket Is a DEG/ENaC Protein Required for Mechanical Nociception in Drosophila Larvae. *Current Biology*, *20*(5), 429–434. https://doi.org/10.1016/j.cub.2009.12.057

Zhou, X., Slone, J. D., Rokas, A., Berger, S. L., Liebig, J., Ray, A., Reinberg, D., & Zwiebel, L. J. (2012). Phylogenetic and Transcriptomic Analysis of Chemosensory Receptors in a Pair of Divergent Ant Species Reveals Sex-Specific Signatures of Odor Coding. *PLoS Genetics*, *8*(8), e1002930. https://doi.org/10.1371/journal.pgen.1002930
